# Directional Gene-Level Concordance and Methodological Constraints in Blood Transcriptomic and DNA Methylation Studies of Parkinson’s Disease

**DOI:** 10.64898/2026.05.17.725808

**Authors:** Chiragh Dewan, Inayat Chauhan, Kanika Sharma, Sangeeta Sharma, Rupinderjeet Kaur

**Author notes:** Corresponding author: Rupinderjeet Kaur, Department of Biotechnology, DAV College, Sector 10, Chandigarh.

## Abstract

Assessing reproducibility across different molecular profiling studies is a persistent methodological challenge (Zhang et al., 2009; Sweeney et al., 2017; Ioannidis, 2005). Differences in platform technology, cohort composition, analytical pipelines, and feature definitions often make it difficult to interpret cross-study comparisons based solely on gene-identity overlap.

In this study, we conducted a retrospective computational analysis of seven publicly available analytical datasets (including alternative analytical pipelines applied to the same cohort) derived from five biologically independent peripheral blood transcriptomic and DNA methylation cohorts, comprising 3,487 samples (1,824 Parkinsons disease cases and 1,663 controls). Reproducibility was evaluated using gene-identity overlap, enrichment-based comparisons, and a permutation-based framework to assess directional consistency of effect estimates across datasets. We also tested the robustness of results by varying false discovery rate thresholds and applying alternative probe-to-gene collapsing strategies. All analyses were performed using reproducible workflows implemented in R and Python with fixed random seeds.

Across independent cohorts, gene-identity overlap was generally limited, with enrichment ratios close to one, especially when datasets were generated using different platforms. In several datasets, limited numbers of statistically significant features further constrained overlap-based comparisons. In contrast, directional consistency showed greater stability. High levels of directional consistency were observed across independent cohort comparisons when restricted to overlapping statistically significant features and remained stable across statistical thresholds (90.0% at FDR < 0.05 and 82.8% at FDR < 0.10). When evaluated across the full shared gene universe without conditioning on statistical significance, directional consistency was substantially lower (∼30 to 32%) but remained significantly above permutation-based null expectations. Permutation testing confirmed that the observed directional consistency exceeded what would be expected by chance. A combined analysis including methodological replicates (n ≥ 3 datasets) showed 98.3% directional consistency; however, this estimate includes non-independent analytical pipelines applied to the same cohort and reflects analytical stability rather than independent biological replication. Rather than introducing a new statistical method, this study examines how commonly used reproducibility metrics behave under crossstudy heterogeneity and identifies their practical limitations and appropriate use boundaries.

## INTRODUCTION

Reproducibility across independent molecular profiling studies is a central challenge in computational biology (Ioannidis, 2005). High-throughput transcriptomic and epigenomic datasets are generated using diverse platforms, analytical workflows, and cohort designs, resulting in heterogeneous feature definitions and statistical properties (Irizarry et al., 2005; MAQC Consortium, 2006). These differences complicate cross-study comparisons and frequently lead to limited gene-level replication, even when studies nominally investigate similar biological conditions. Recent multi-omic investigations have further emphasized the complexity and heterogeneity of molecular signatures in Parkinson’s disease (Dayan et al., 2025).

Gene-identity overlap is commonly used to assess reproducibility; however, overlap-based metrics are highly sensitive to analytical thresholds, sample size, platform-specific measurement characteristics, and the underlying feature universe (Zhang et al., 2009; Sweeney et al., 2017). These limitations are particularly pronounced for DNA methylation studies, where large numbers of CpG loci are mapped to genes, inflating apparent overlap and obscuring meaningful comparisons (Chen et al., 2013; Pidsley et al., 2013). As a result, gene-identity overlap may provide an unstable or misleading representation of reproducibility across heterogeneous datasets.

Directional consistency of effect estimates offers an alternative framework for evaluating reproducibility that is less dependent on feature identity (Cahill et al., 2018). By focusing on concordance in effect direction rather than exact feature overlap, this approach captures stability in statistical signal across studies despite platform and analytical heterogeneity. However, the behavior of directional consistency under varying analytical conditions and annotation strategies has not been systematically evaluated.

In this study, we conducted a methodological assessment of reproducibility across publicly available transcriptomic and DNA methylation datasets using Parkinson’s disease as a case study (Kurvits et al., 2021; Dayan et al., 2025; Chen-Plotkin, 2018; Scherzer et al., 2007; Shamir et al., 2017). Although seven datasets were processed, not all contributed equally to inferential analyses. Some datasets yielded no statistically significant features under prespecified thresholds, and two analyses (GSE6613 and GSE6613-CEL) represent alternative analytical pipelines applied to the same underlying cohort. Consequently, effective independent cohort comparisons are restricted to four primary datasets. All references to “seven datasets” reflect processed analytical inputs rather than independent biological replications. Importantly, the disease context serves solely as an analytical anchor. We compare overlap-based and directional reproducibility metrics and evaluate robustness through permutation-based inference and sensitivity analyses. All conclusions are methodological and do not constitute biological interpretation.

Several approaches for assessing cross-study reproducibility have been proposed, including effect-size meta-analysis and vote-counting strategies (Sweeney et al., 2017). The present study does not introduce a new estimator but provides the first systematic empirical evaluation of how overlap-based and directional reproducibility metrics behave across both transcriptomic and epigenomic platforms under real-world analytical heterogeneity. By benchmarking these metrics against permutation-calibrated null expectations and testing their robustness to threshold and annotation perturbations, we identify conditions under which each metric succeeds or fails.

Our objective is not to identify reproducible Parkinson’s disease biomarkers, but to characterize failure modes, deterministic artifacts, and scenarios in which apparent reproducibility can arise without biological replication. By doing so, we aim to provide practical guardrails for the safe interpretation of overlap, enrichment and directional consistency analyses in large-scale omics studies.

## MATERIAL AND METHODS

### Study Design

This study was conducted as a retrospective computational analysis of publicly available transcriptomic and DNA methylation datasets. The analytical framework was designed to evaluate statistical reproducibility under heterogeneous analytical and platform conditions rather than to generate or validate biological hypotheses. Throughout the manuscript, the term directional consistency refers specifically to agreement in the sign of effect estimates across datasets and is used consistently to avoid terminological ambiguity.

All datasets were obtained from the Gene Expression Omnibus (GEO) database (Edgar et al., 2002). Seven publicly available datasets were included, encompassing a total of 3,487 samples, comprising 1,824 Parkinson’s disease cases and 1,663 control samples.

### Data Selection and Characteristics

A total of seven datasets were analyzed, derived from five biologically independent cohorts (GSE6613 and GSE6613-CEL share underlying samples). These included three transcriptomic expression datasets and four DNA methylation datasets generated using Affymetrix and Illumina platforms. Expression datasets included GSE6613, GSE99039, and GSE165083 (GPL11154), while methylation datasets included GSE111629, GSE72774, GSE145361, and GSE165083 (methylation platform). All datasets were derived from peripheral blood samples.

For example, GSE6613-CEL included 25 Parkinson’s disease cases and 37 controls (n = 62), whereas GSE99039 comprised 191 cases and 212 controls (n = 403). The largest methylation dataset, GSE145361, included 1,889 samples (959 cases and 930 controls) and interrogated approximately 485,512 CpG loci. Unless otherwise specified, all references to GSE6613 in comparative analyses refer to the GSE6613-CEL pipeline for consistency. Quality control considerations for cross-study integration were informed by established meta-analysis frameworks (Kang et al., 2012). Feature counts reflect post-quality-control analytical universes and therefore varied across platforms and datasets.

### Data Processing

Expression datasets were normalized using platform-appropriate procedures (robust multiarray average [RMA] for Affymetrix arrays and standard library-size normalization for RNA-sequencing data), and low-variance genes were filtered prior to downstream analysis. Gene identifiers were standardized to ensure consistent annotation across datasets. After normalization and filtering of low-abundance features, expression datasets retained approximately 8,000 to 21,000 genes for differential analysis, depending on platform characteristics. To account for potential latent structure and technical variation, standard normalization and quality control considerations were applied, as commonly addressed in high-dimensional genomic studies (Price et al., 2010).

DNA methylation beta-value matrices were obtained as pre-processed data from GEO. Beta-values were clipped to the interval [10^−6^, 1−10^−6^] to avoid boundary instability and then transformed to M-values using M = log_2_(β/(1−β)) for linear modelling (Aryee et al., 2014; Fortin et al., 2017; Du et al., 2010). Probe-level quality control included filtering of low-variance and high-missingness probes (Pidsley et al., 2013; Chen et al., 2013). Cross-reactive probes and probes overlapping common single-nucleotide polymorphisms were not removed, as datasets were obtained as pre-processed matrices rather than reprocessed from raw IDAT files. This is acknowledged as a limitation. Proportions of major blood cell types were estimated (using EpiDISH) but not included as covariates in differential analyses. Following quality control, methylation datasets retained approximately 393,000 to 485,000 CpG sites per dataset. The reduced feature count for GSE111629 (393,217 CpGs vs 485,512 in other 450K datasets) reflects upstream submitter-specific quality control and probe filtering applied prior to GEO deposition.

For microarray datasets, differential analysis was performed at the gene level following probe-to-gene mapping and collapsing of multiple probes per gene. Consequently, probe-level feature counts reported in dataset descriptions (e.g., 9,779 probes for GSE99039) differ from the gene-level universes used in DEG analyses (e.g., 2,670 unique genes after mapping and filtering). Cell-type fraction estimates were available only for GSE165083-methylation (Table S1) (Houseman et al., 2012; Newman et al., 2015). No other datasets included these estimates, and therefore no cross-dataset covariate harmonization was possible.

Differentially methylated regions were mapped to genes for diagnostic comparison; however, these mappings were interpreted cautiously due to their broad and uneven genomic coverage. As a result, DMR-mapped gene universes frequently exceeded 22,000 genes, highlighting the structural asymmetry between expression-based and methylation-based feature definitions.

### Implementation and Software

All analyses were performed using R and Python (Gentleman et al., 2004). Differential expression and methylation analyses were conducted using ordinary least squares (OLS) regression with covariate adjustment, implemented in Python using the statsmodels library (Seabold and Perktold, 2010). Variance moderation was applied using a simplified empirical Bayes shrinkage procedure to stabilize variance estimates across features. DNA methylation preprocessing and region-level analyses were performed using standard Bioconductor workflows. Region-level methylation analysis was performed using DMRcate with default parameters (bandwidth λ = 1000 bp, scaling factor C = 2, minimum 3 CpGs per region), and genomic annotation of regions was conducted using ChIPseeker (Peters et al., 2015; Yu et al., 2015). Cell-type fraction estimation was performed using EpiDISH where applicable (Teschendorff et al., 2017). Genomic interval operations were implemented using the pyranges library (Stovner and Sætrom, 2020). Permutation testing, robustness analyses, and sensitivity analyses were implemented using custom scripts. Full computational reproducibility was ensured through the use of fixed random seeds.

### Differential Analysis

Differential expression analyses were conducted primarily using ordinary least squares regression with empirical Bayes variance shrinkage (Ritchie et al., 2015; Smyth, 2004). For diagnostic comparison, standard two-sample t-tests were also computed; however, all primary results and gene universe definitions reported in the manuscript are based on the OLS regression-derived statistics unless explicitly stated otherwise. The OLS regression path applied stricter feature-level quality control than the t-test path (e.g., 9,779 versus 43,088 features for GSE99039), resulting in smaller analytical universes; false discovery rate correction was applied independently within each method. Differential methylation analyses were conducted using analogous linear modeling frameworks appropriate to array-based methylation data.

Where valid numeric covariate data existed, models included age and sex as covariates (GSE72774 and GSE111629); sex only was included for GSE6613-CEL, GSE145361, GSE165083, and GSE165083-GPL11154; batch was additionally included for GSE99039. The specific covariates available per dataset are documented in Supplementary Table S1. Cell-type fraction estimates were computed for GSE165083-methylation but were not included as covariates in any differential analysis. DMR analyses used unadjusted models with disease state as the sole predictor. When specific covariates were unavailable for a given dataset, models were fit using the maximal covariate set supported by that dataset; no covariate imputation was performed.

Multiple-testing correction was applied using the False Discovery Rate (Benjamini and Hochberg, 1995). Platform-specific thresholds were used to accommodate differences in signal distributions and feature densities (FDR < 0.10 for microarray expression datasets; FDR < 0.05 for RNA-sequencing and methylation datasets). These thresholds were selected to balance sensitivity and specificity rather than to influence biological interpretation. The primary effect-size threshold was |log_2_FC| ≥ 0.05, selected to retain features with small but directionally consistent signals given the diagnostic focus of this study.

### Directional Consistency Analysis

To evaluate reproducibility beyond gene-identity overlap, a permutation-based framework was used to test whether agreement in effect direction across datasets exceeded random expectation. Formal inferential testing was restricted to comparisons involving at least three analytical datasets (n ≥ 3); comparisons involving only two datasets were treated as descriptive.

Only features present in at least three independent datasets were included in directional consistency analyses, where feature presence was defined as being measured and retained after quality control within a dataset, independent of statistical significance. Directional consistency was defined as concordant up- or down-regulation (or hyper- or hypomethylation) across datasets for a given feature. Directional consistency was computed as D = n_concordant / (n_concordant + n_discordant), excluding neutrals (effect size = 0) from the denominator.

Within dataset and between dataset comparisons are reported separately and serve distinct descriptive purposes; no biological replication claims are made for unmatched cohort comparisons.

### Cross-Dataset Concordance

Overlap enrichment analyses and rank-based concordance assessments were conducted to describe statistical behavior across datasets. All analyses were restricted to dataset-specific feature universes to avoid artifacts arising from platform heterogeneity.

### Robustness Evaluation

Robustness of directional findings was evaluated using permutation-based resampling procedures and sensitivity analyses. Empirical null distributions were generated using 10,000 resampling iterations, in which effect direction labels were randomly reassigned without replacement to approximate the distribution expected under the null hypothesis of no reproducible directionality (Zhang et al., 2009). Pairwise null calibration analyses used label permutation without replacement. Permutation p-values for the primary directional consistency analysis were computed using a mid-p correction, defined as (count + 0.5)/(n + 1), whereas pairwise null calibration used the standard formula (count + 1)/(n + 1). Both approaches yield conservative p-values; the choice does not materially affect reported significance levels. 95% confidence intervals for directional consistency estimates were computed using the Wilson score interval for binomial proportions; bootstrap resampling (10,000 iterations) was used for sensitivity validation. These permutation procedures condition on marginal direction frequencies but assume gene-level independence. Co-expression and correlated gene structure may violate this assumption and could modestly influence null distribution behavior. All stochastic procedures were executed using a fixed random seed (set to 42) to ensure full computational reproducibility.

### Threshold Sensitivity Analysis

To assess whether directional consistency results were dependent on specific threshold choices, pairwise concordance analyses were repeated across FDR thresholds of 0.01, 0.05, and 0.10 using a fixed effect size threshold (|log_2_ fold-change| ≥ 0.1), which is stricter than the primary |log_2_FC| ≥ 0.05 cutoff and was selected to reduce the noise floor in cross-dataset comparisons, where small-effect features are most susceptible to sign instability. For independent cohort comparisons between GSE6613-CEL and GSE99039, directional consistency remained high despite threshold relaxation, with 90.0% agreement at FDR < 0.05 (n = 30 genes) and 82.8% agreement at FDR < 0.10 (n = 58 genes). Although overlap size increased with relaxed thresholds, directional consistency remained stable, indicating that concordance patterns were not driven by arbitrary threshold selection.

### Gene Annotation Sensitivity Analysis

For expression datasets in which multiple probes mapped to a single gene, probe-level results were collapsed to gene-level summaries. The primary analysis used selection of the probe with maximum absolute log_2_ fold-change per gene. Sensitivity analyses were conducted using two alternative strategies: selection based on the lowest adjusted p-value per gene and calculation of the median log_2_ fold-change per gene. All three approaches yielded comparable full-universe directional consistency (30.5%–32.4%). These values reflect concordance across the shared analytical universe rather than intersection-restricted significant features and therefore represent unconditional directional consistency rates.

### Meta-Analytic and Multi-Omic Procedures

Between-dataset heterogeneity was quantified using descriptive metrics. All meta-analytic outputs were treated strictly as descriptive summaries rather than evidence of replication. Multi-omic integration and pathway enrichment analyses were performed solely as methodological demonstrations and were not interpreted as biological findings. No data recalibration or inflation correction procedures were applied. Batch was included as a covariate in the GSE99039 linear model where a valid batch variable was available; no post-hoc batch correction was performed on any dataset.

## RESULTS

### Dataset Overview and Quality Assessment

Following quality control and preprocessing, all seven datasets were retained for downstream analyses, comprising a total of 3,487 samples across transcriptomic and DNA methylation modalities. Expression datasets ranged in size from 26 to 403 samples, while methylation datasets ranged from 28 to 1,889 samples, reflecting differences in cohort design and platform coverage. All datasets were derived from peripheral blood samples of Parkinson’s disease cases and matched controls (Scherzer et al., 2007; Shamir et al., 2017).

Global data distributions were broadly comparable across datasets after preprocessing (Figures 2 and 4). Principal component analysis revealed dataset-specific clustering patterns consistent with platform and cohort differences, without evidence of extreme outlier samples (Figures 3 and 5). These observations indicate adequate data quality for downstream comparative analyses while highlighting expected inter-dataset heterogeneity.

**Figure 1.**
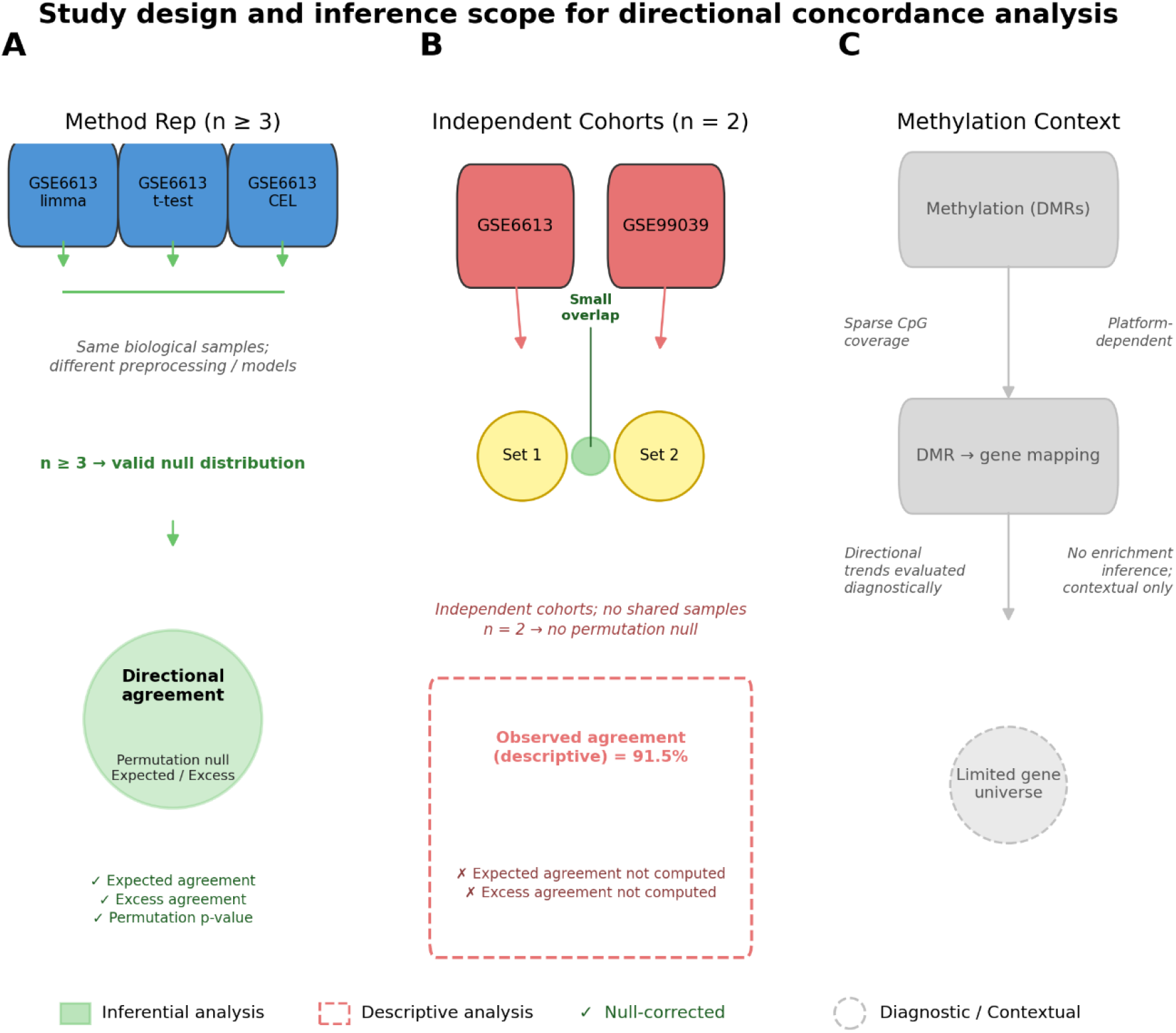
Study design and inference scope for directional consistency analysis. Schematic overview of the reproducibility framework. (A) Method-replication analysis (n ≥ 3 datasets) using permutation-based inference to estimate excess directional consistency. (B) Independent cohort comparison (n = 2) reporting descriptive directional consistency only. (C) Contextual considerations for methylation data, including probe-to-gene mapping constraints and platform-dependent annotation effects.

**Figure 2.**
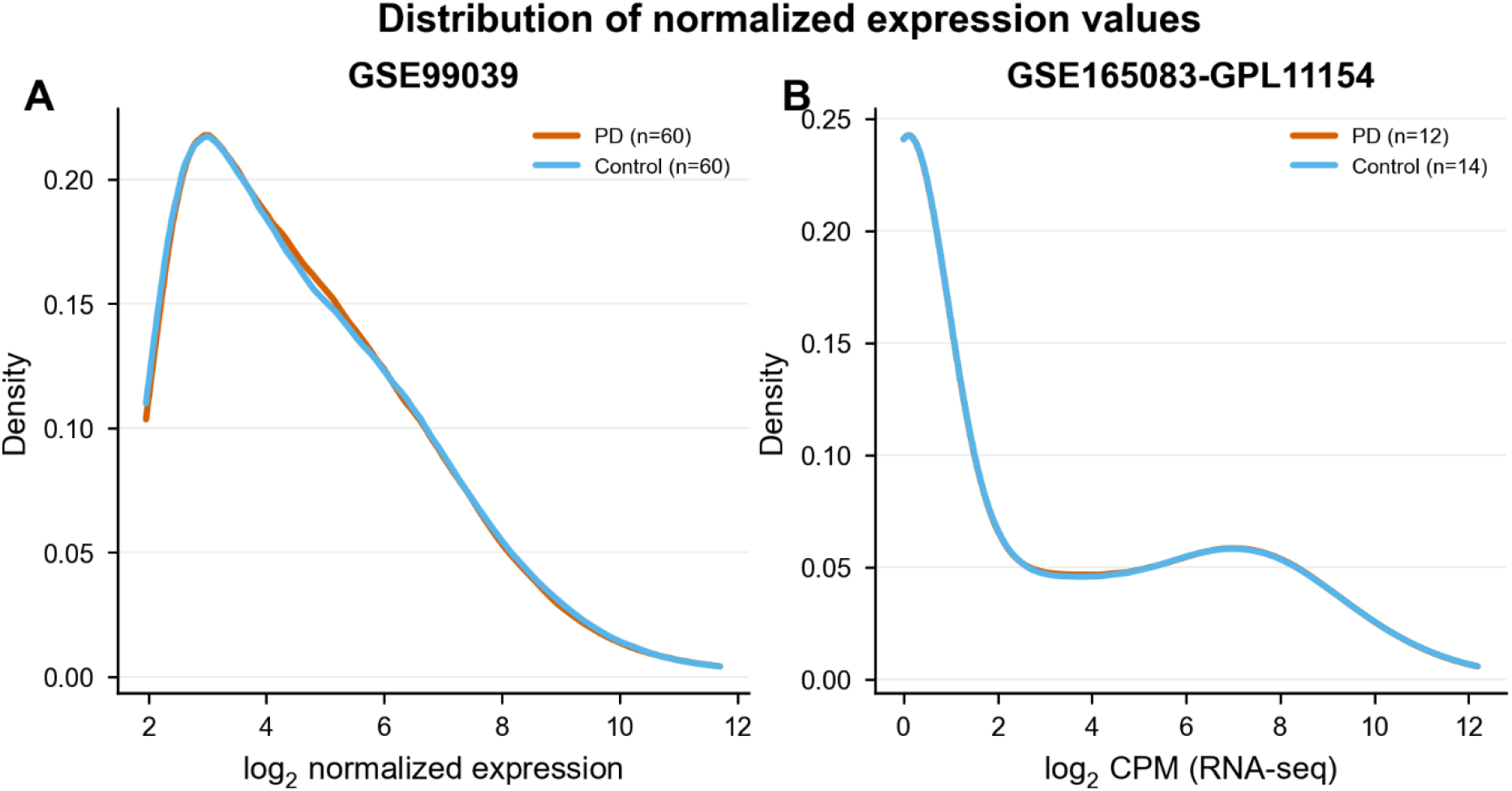
Distribution of normalized expression values. Density plots of normalized expression values for representative microarray (GSE99039) and RNA-seq (GSE165083-GPL11154) datasets. Thin lines show persample densities (subsampled); thick lines show within-group medians (PD = vermillion, Control = sky blue).

**Figure 3.**
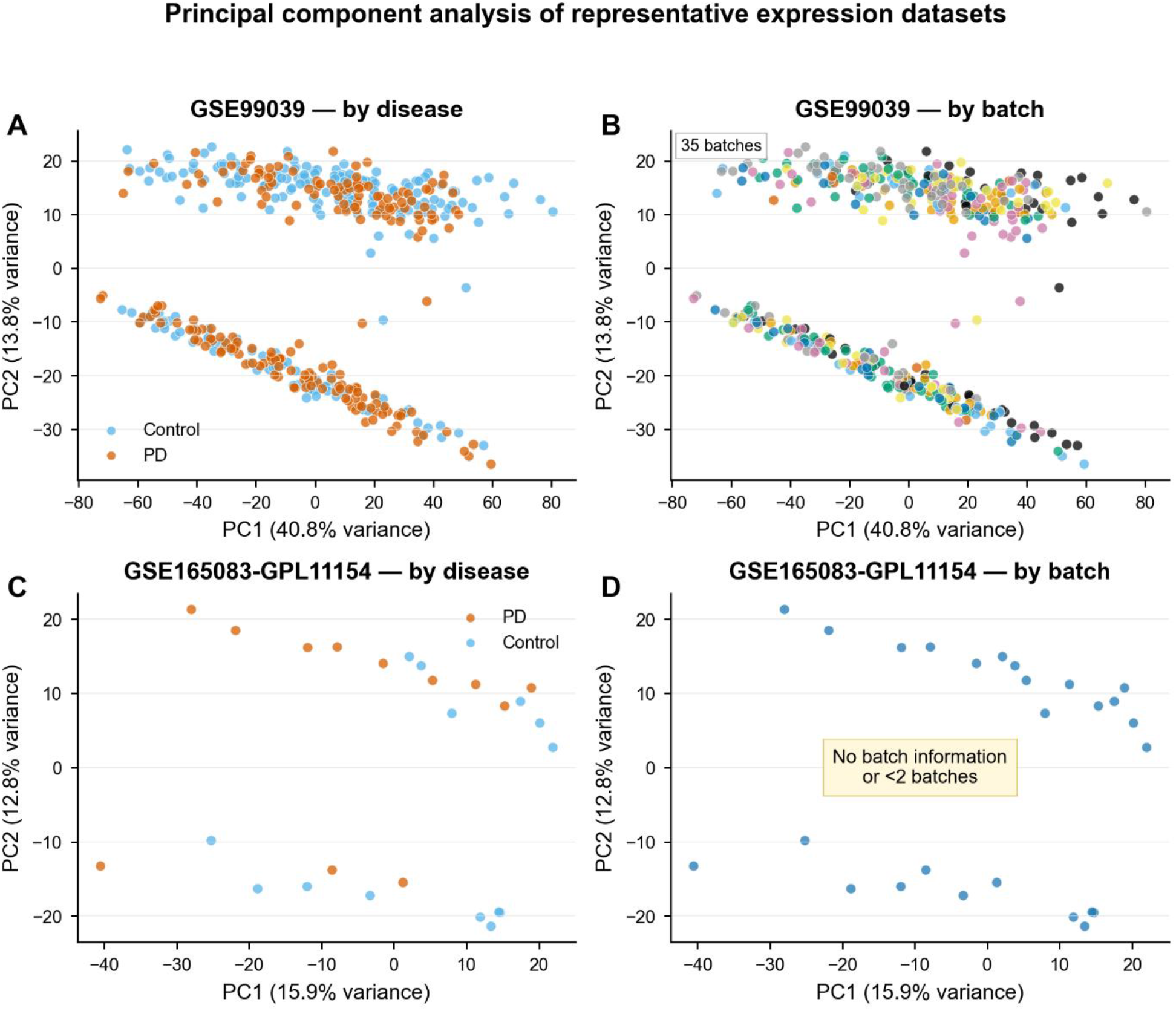
Principal component analysis of representative expression datasets. PCA of top-1000 variance features in GSE99039 and GSE165083-GPL11154. Samples are colored by disease status (left column) and batch (right column, where available). Axes indicate variance explained. Disease-related separation is modest relative to overall variance structure.

**Figure 4.**
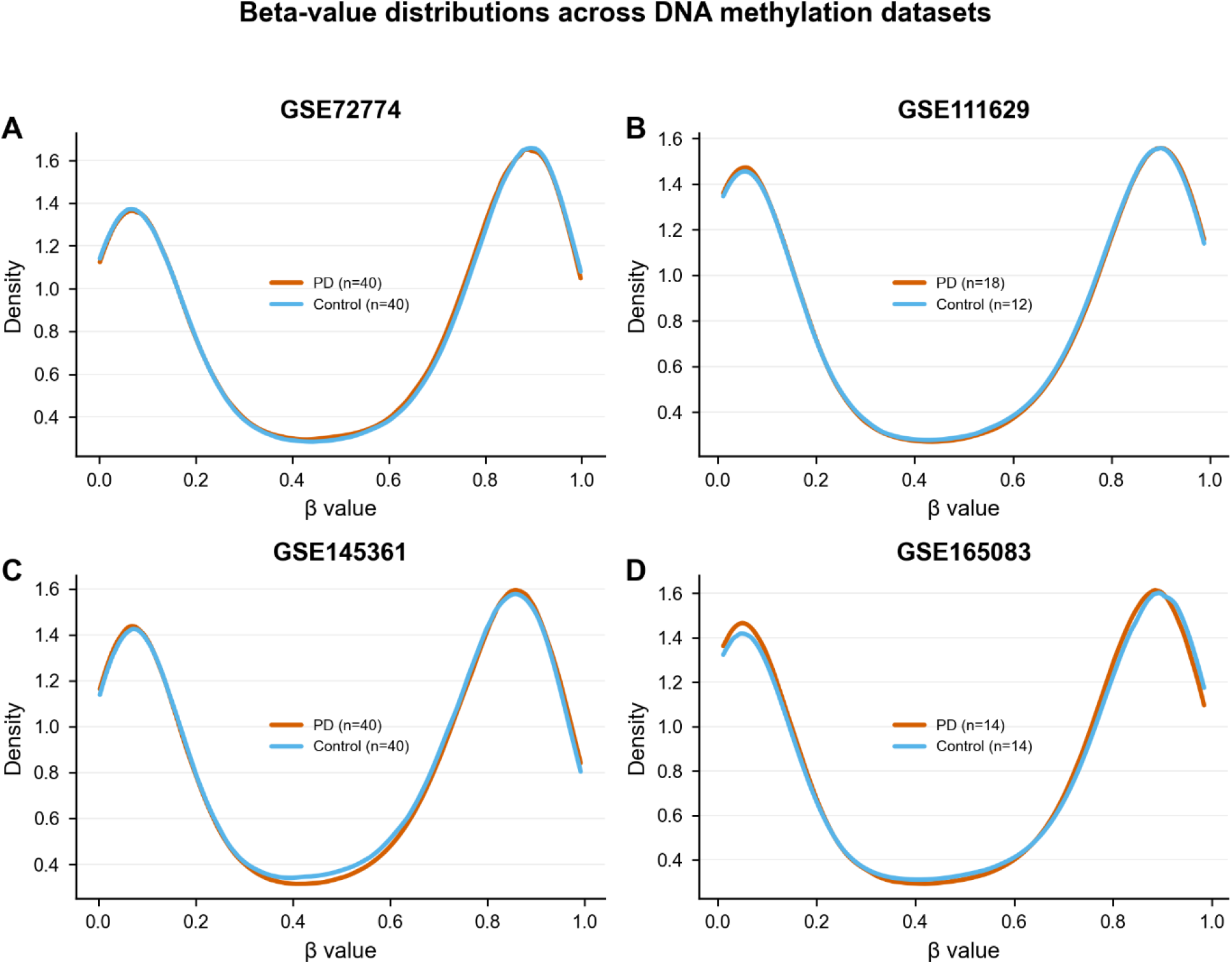
Beta-value distributions across DNA methylation datasets. Density plots showing distribution of CpG methylation β values across all four methylation datasets. Thin lines show per-sample densities; thick lines show within-group medians. All datasets exhibit the characteristic bimodal distribution of methylation levels.

**Figure 5a.**
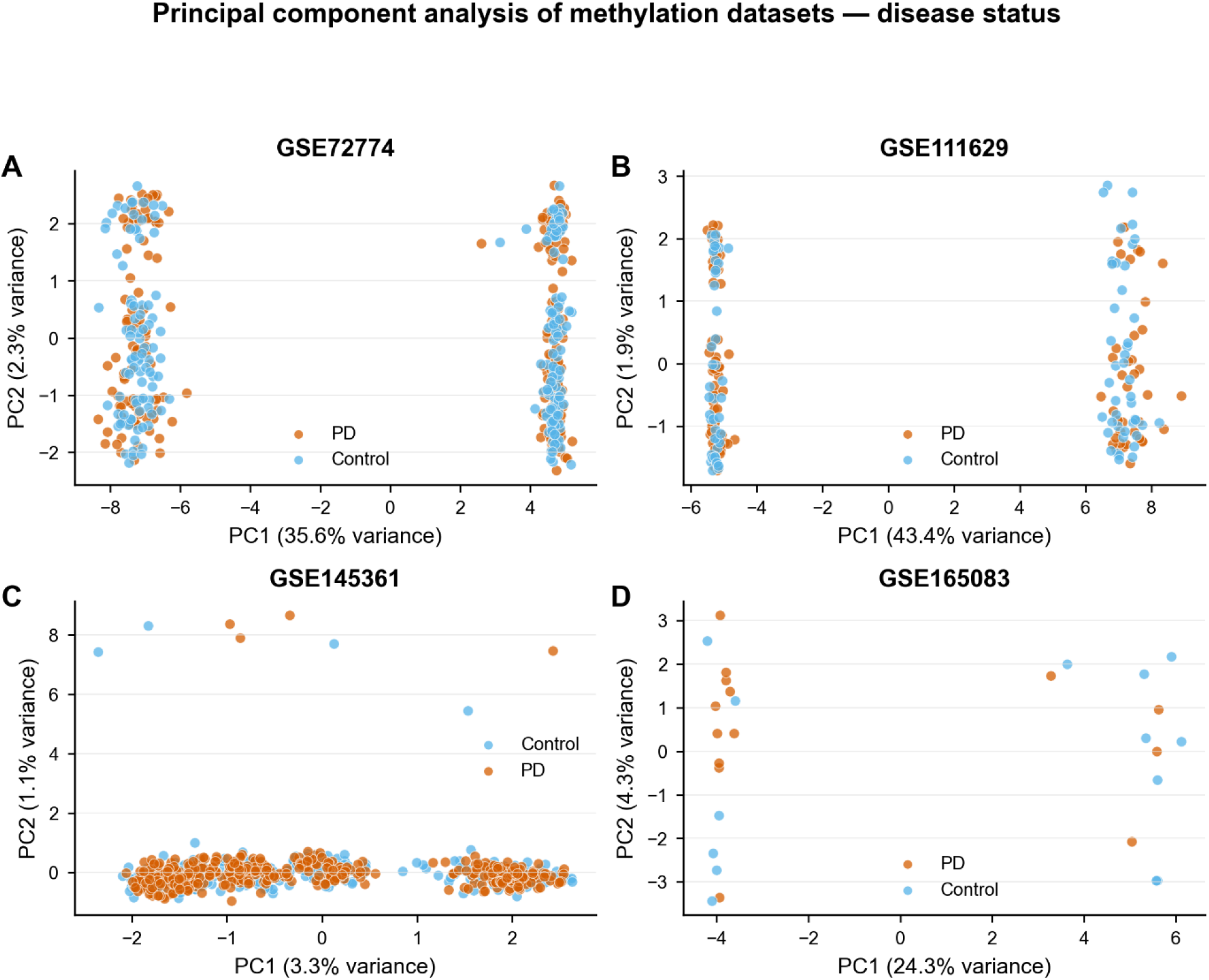
PCA of methylation datasets — disease status. PCA based on top-1000 variance CpGs within each dataset. Samples colored by PD (vermillion) vs Control (sky blue). Dataset-local; not directly comparable across cohorts.

**Figure 5b.**
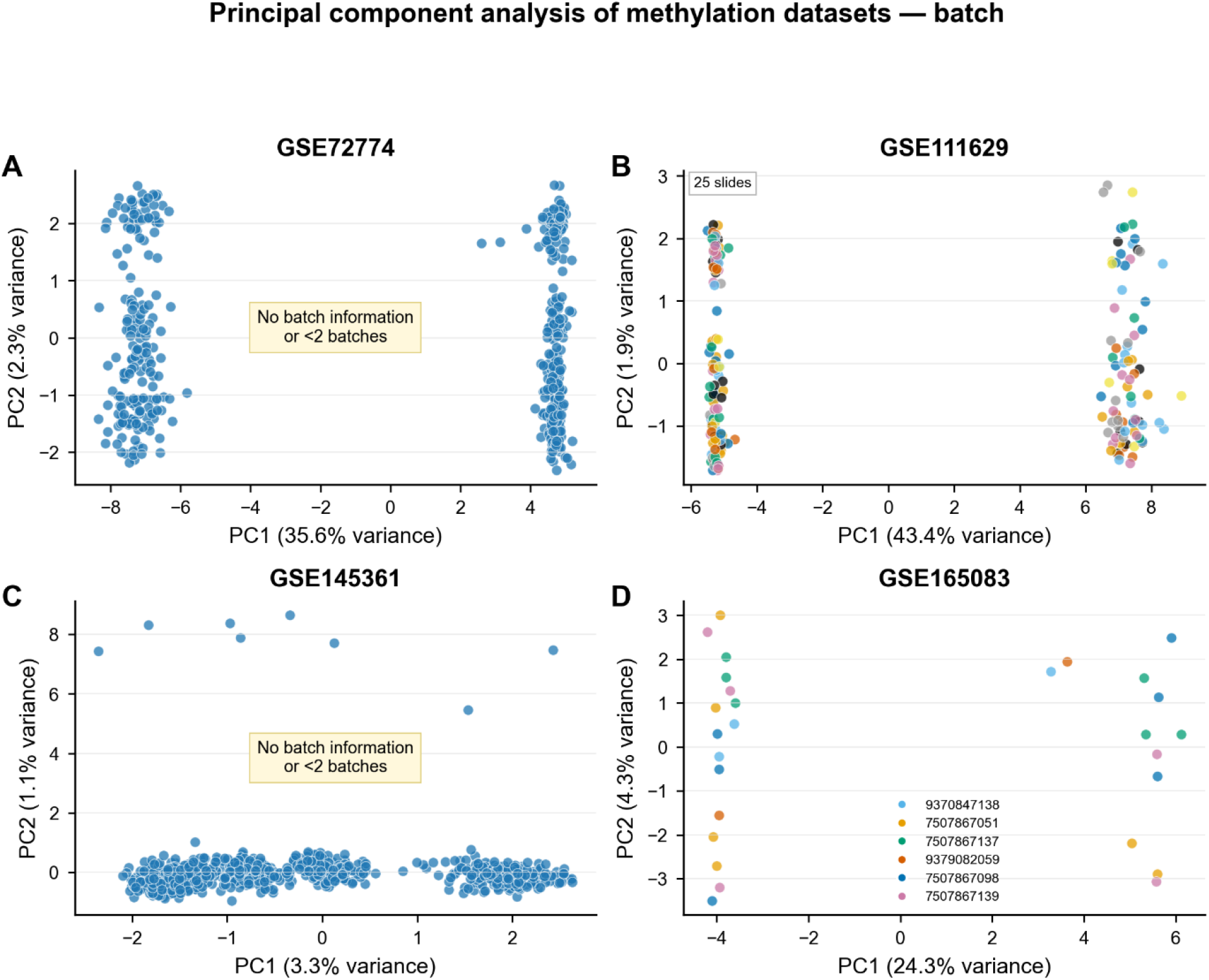
PCA of methylation datasets — batch. Same PCA projections as Figure 5a, with samples colored by Sentrix slide ID where available.

Genomic inflation factors were reported as diagnostic indicators of large-scale correlation structure and annotation density rather than as measures requiring correction; all downstream analyses were conducted without recalibration (Devlin and Roeder, 1999). At the individual CpG level, genomic inflation was well-controlled across all methylation datasets (GSE145361 λ=1.04, GSE72774 λ=0.99, GSE111629 λ=1.04, GSE165083 λ=1.00), confirming that the substantially elevated region-level λGC values (123.1 and 114.7 respectively) reflect the aggregation of correlated CpG signals within DMRs rather than systematic miscalibration at the probe level.

### Differential Analysis Summary

Differential expression and methylation analyses identified variable numbers of statistically significant features across datasets, reflecting differences in sample size, platform technology, and analytical universe. Expression datasets identified between 0 and 1,596 significant features, depending on dataset and analytical pipeline, whereas methylation datasets yielded substantially larger sets of differentially methylated region (DMR)-mapped genes, frequently exceeding 18,000 genes per dataset (Table 2) (Vallerga et al., 2020; Lie et al., 2025). These disparities underscore the structural asymmetry between expression-based and methylation-based feature definitions and motivated subsequent analyses focused on reproducibility metrics that are robust to heterogeneous feature universes.

**Table 1:**
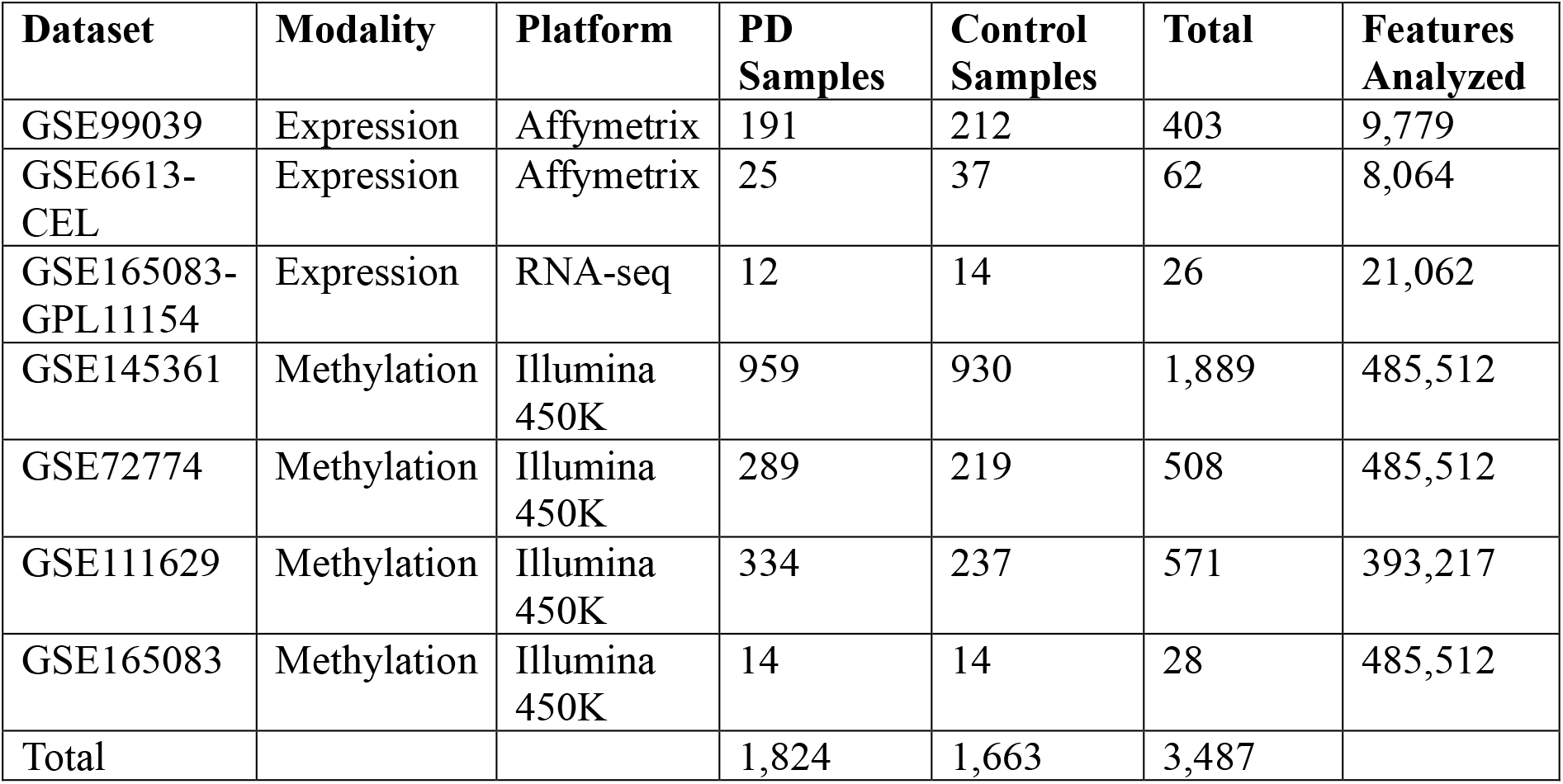
Summary of included datasets. Characteristics of seven peripheral blood transcriptomic and DNA methylation datasets analyzed, including platform, sample counts, and features retained after preprocessing. In total, 3,487 samples (1,824 PD and 1,663 controls) were included. GSE6613-CEL represents the same underlying cohort as GSE6613, reprocessed from raw CEL files using an alternative analytical pipeline. GSE99039 metadata contained 557 entries; 119 samples with unknown disease status were excluded, and a further 35 samples were removed during quality control. GSE6613-CEL metadata contained 105 samples; 33 samples labelled ‘Neurological Control’ were merged with 22 regular controls, and 43 samples were excluded during quality control.

**Table 2:**
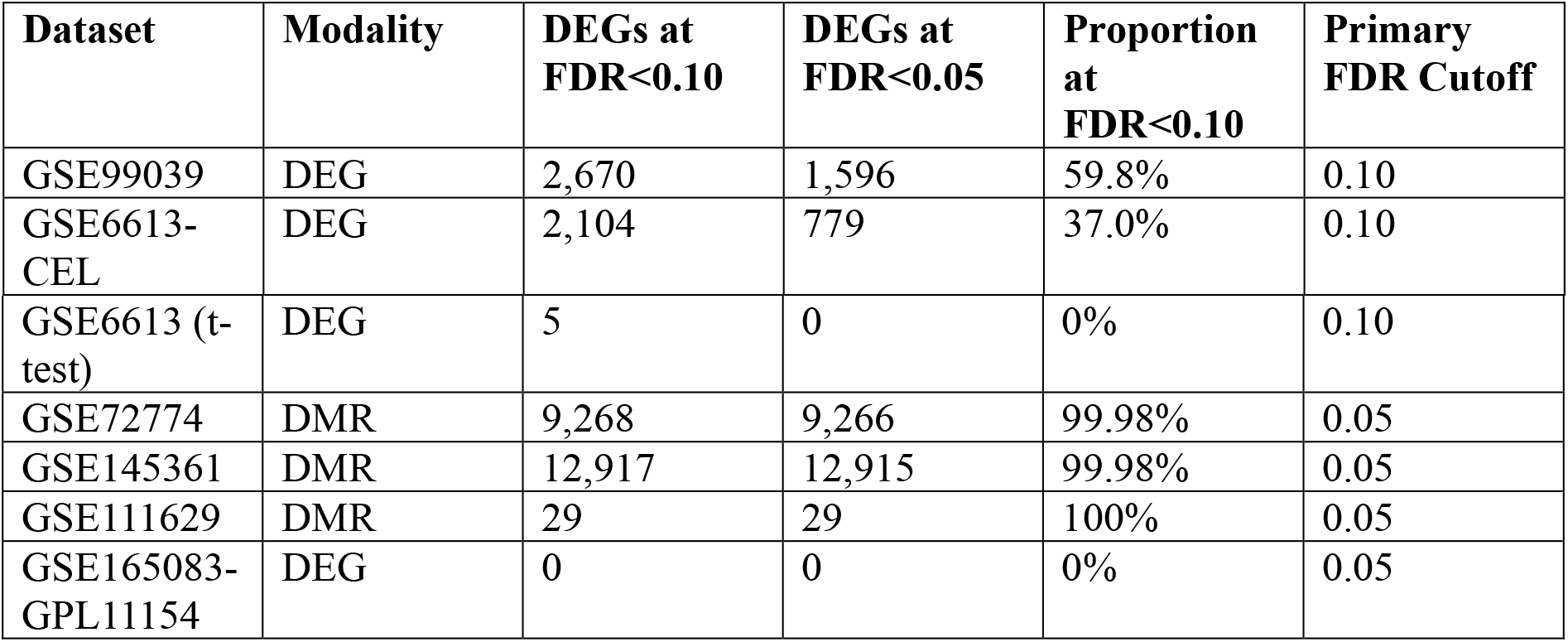
Differential analysis summary per dataset. Post-quality-control analytical universes, number of statistically significant features, and corresponding FDR thresholds. Differences in the proportion of significant features reflect platform-specific feature definitions rather than biological interpretation. The GSE6613 (t-test) row is a methodological diagnostic comparison included for illustrative comparison only.

The OLS regression framework was informed by the limma methodology (Ritchie et al., 2015; Smyth, 2004) but was implemented independently in Python using statsmodels; it does not call the R limma package directly.

### Overlap Enrichment Analysis

Within-dataset methodological comparisons demonstrated strong overlap enrichment, indicating internal analytical consistency. In contrast, independent cohort comparisons exhibited enrichment ratios generally close to unity, reflecting limited gene-identity replication across datasets. Methylation datasets showed near-complete overlap across comparisons, attributable to the expansive and deterministic nature of DMR-based gene mappings rather than shared biological signal (Figure 6, Table 3) (Henderson-Smith et al., 2019; Moore et al., 2014; Du et al., 2010). Although some overlaps reached nominal statistical significance, enrichment ratios were generally close to unity.

**Table 3:**
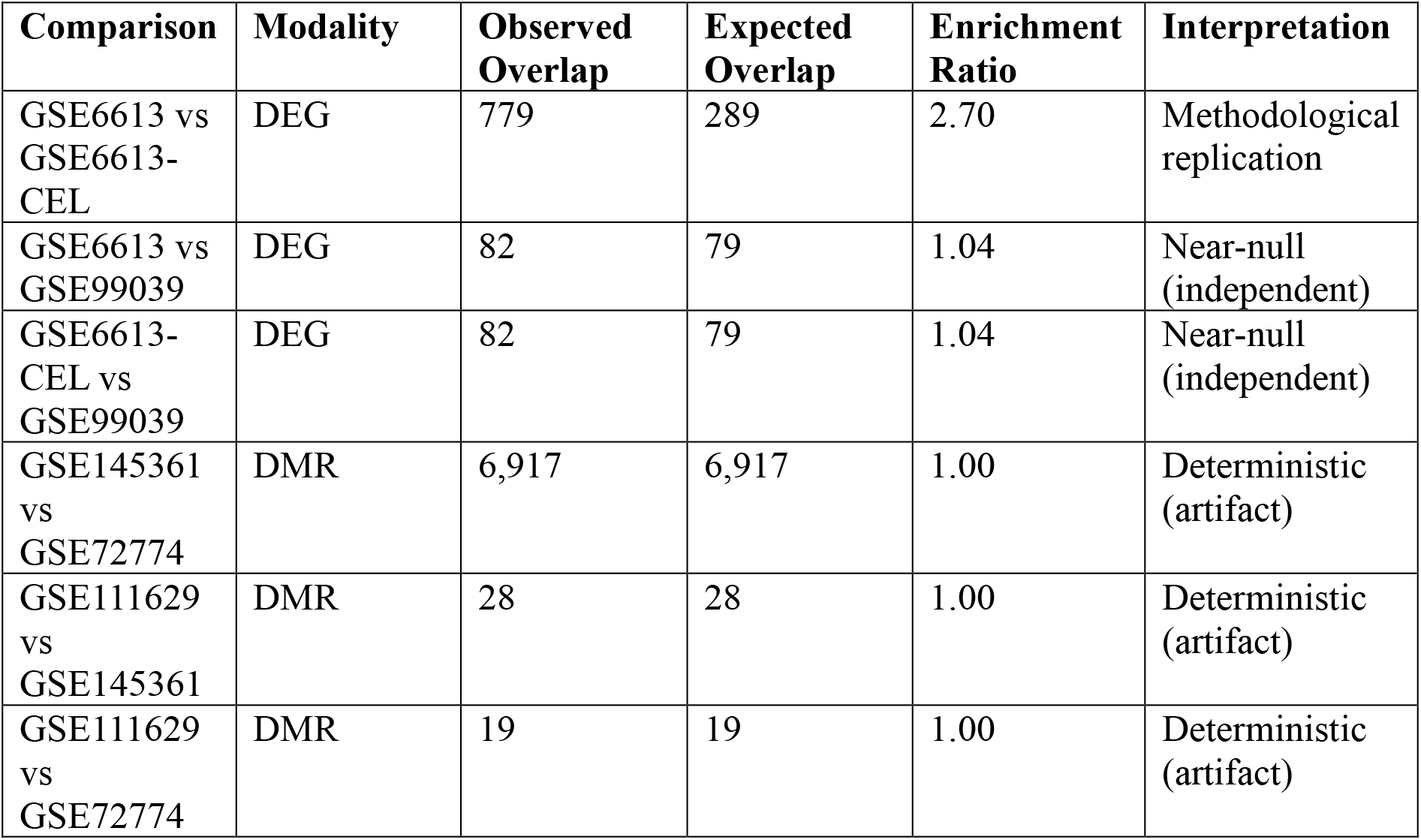
Overlap enrichment analysis. Observed and expected overlaps computed using dataset-specific feature universes. Enrichment ratios near unity indicate limited gene-identity replication across independent cohorts. Deterministic overlaps observed for methylation datasets reflect large and highly overlapping DMR-mapped gene universes rather than shared biological signal.

**Figure 6.**
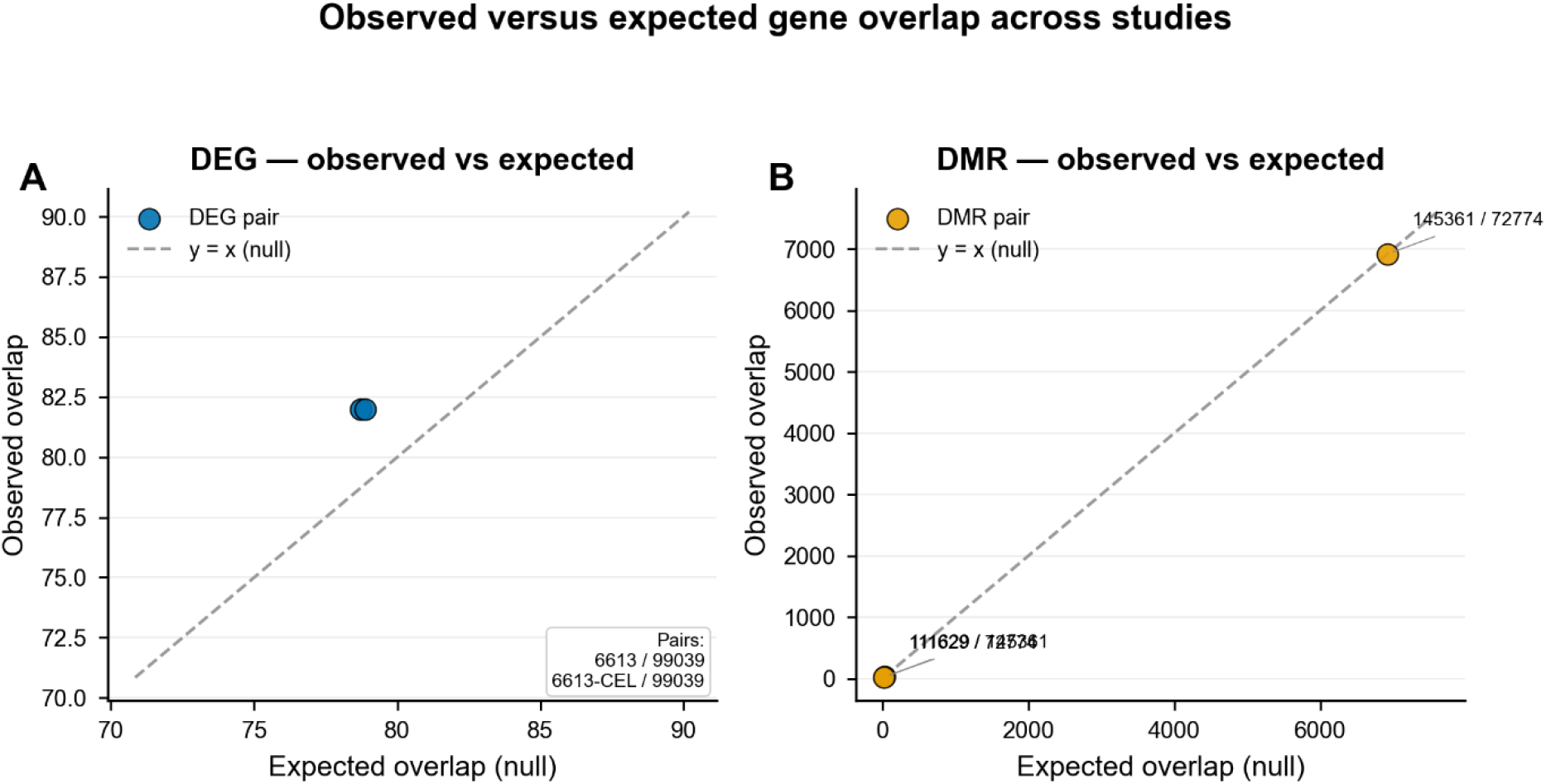
Observed versus expected gene overlap across studies. Scatter plots comparing observed overlap of significant features to null-expected overlap derived from permutation-based models. (A) DEG overlap. (B) DMR overlap. Dashed line: y = x (null expectation). Methylation overlaps approach near-complete gene coverage due to DMR-to-gene mapping expansion.

### Directional Consistency Analysis

Directional consistency analyses demonstrated substantially greater stability than gene-identity overlap. Alternative analytical methods applied to the same dataset showed complete directional consistency among significant features. Descriptive comparisons between independent cohorts also demonstrated high directional consistency (Table 4), despite limited overlap at the gene-identity level.

**Table 4:**
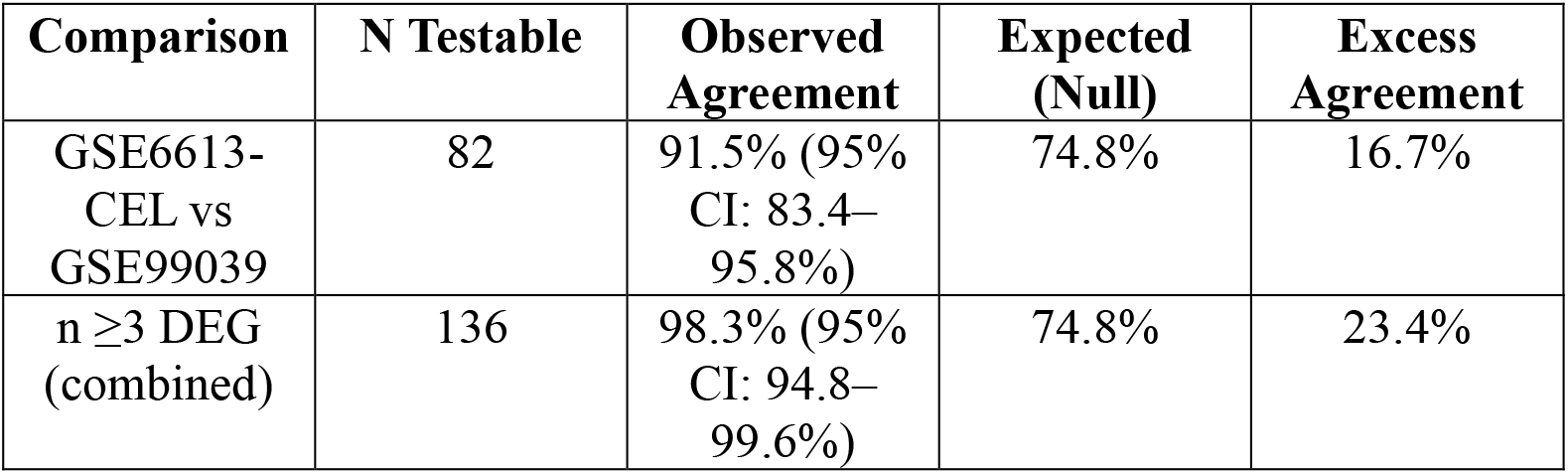
Directional consistency analysis results. DEG, differentially expressed gene. Directional consistency was defined as concordant direction of effect (up–up or down–down) across datasets. Expected agreement was estimated from permutation-based null models. The n ≥3 dataset analysis includes two analytical pipelines applied to the same underlying cohort and therefore reflects analytical stability rather than independent biological replication. The two rows use different analytical frameworks and are not directly comparable.

**Table 5:**
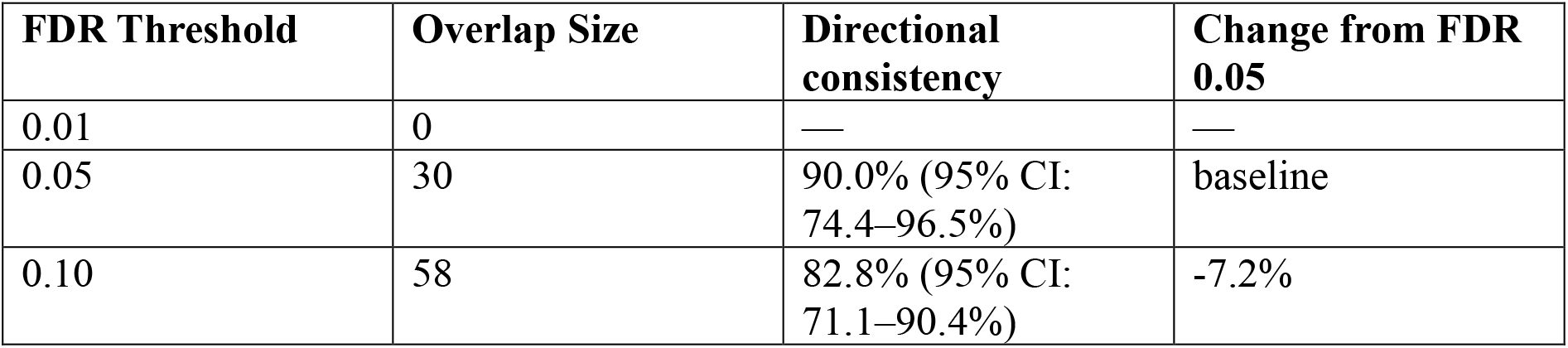
Threshold sensitivity of directional consistency. Sensitivity of directional consistency to FDR threshold selection for the independent cohort comparison. Relaxation of the FDR threshold from 0.05 to 0.10 resulted in near doubling of overlapping features (30 to 58), while directional consistency decreased modestly. No overlapping features were observed at the most stringent threshold (FDR < 0.01).

Inferential analysis involving at least three analytical results (including alternative pipelines applied to the same cohort) revealed significant excess directional consistency beyond random expectations. Because these comparisons combine two independent cohorts with one methodological reanalysis, they reflect analytical stability rather than three independent biological replications and should be interpreted accordingly.

Permutation testing confirmed that observed directional consistency significantly exceeded null expectations in all evaluable comparisons (aggregate permutation p < 0.001 based on 10,000 permutations). These p-values reflect aggregate directional consistency statistics across shared gene sets, not per-gene permutation tests, and support the statistical robustness of the directional consistency metric at the dataset level.

Directional consistency is interpreted exclusively as a measure of analytical stability and not as evidence of shared disease mechanisms or etiological consistency (Dayan et al., 2025; Cahill et al., 2018). Directional consistency reflects consistency in effect sign and does not imply concordance in effect magnitude or biological relevance.

Because GSE6613 and GSE6613-CEL represent alternative analytical pipelines applied to the same underlying biological cohort, the n ≥ 3 directional stability estimate (98.3%) is partially driven by non-independence. A gene identified in both pipelines does not constitute independent biological replication. Therefore, this estimate reflects methodological robustness rather than three independent cohort confirmations.

When evaluated across the full shared gene universe without conditioning on joint statistical significance, directional consistency between GSE6613-CEL and GSE99039 was 30.5% (95% CI: 25.1–36.5%) (Table 7). The higher 91.5% estimate reflects conditioning on genes meeting significance criteria in both datasets and therefore represents a restricted intersection rather than a fixed-universe concordance rate.

### Robustness and Sensitivity Analyses

Permutation-based resampling confirmed that observed directional consistency exceeded random expectation under null models with appropriate Type I error control. Directional consistency metrics remained stable across a wide range of analytical perturbations, including threshold variation and annotation strategy changes.

#### Threshold Sensitivity

Directional consistency remained high across multiple false discovery rate thresholds. For independent cohort comparisons between GSE6613-CEL and GSE99039, concordance was 90.0% at FDR < 0.05 (30 overlapping genes) and 82.8% at FDR < 0.10 (58 overlapping genes) (Figure 8, Table 5). Although overlap size increased with threshold relaxation, the persistence of high directional consistency indicates that concordance patterns are not driven by arbitrary threshold selection.

**Figure 7.**
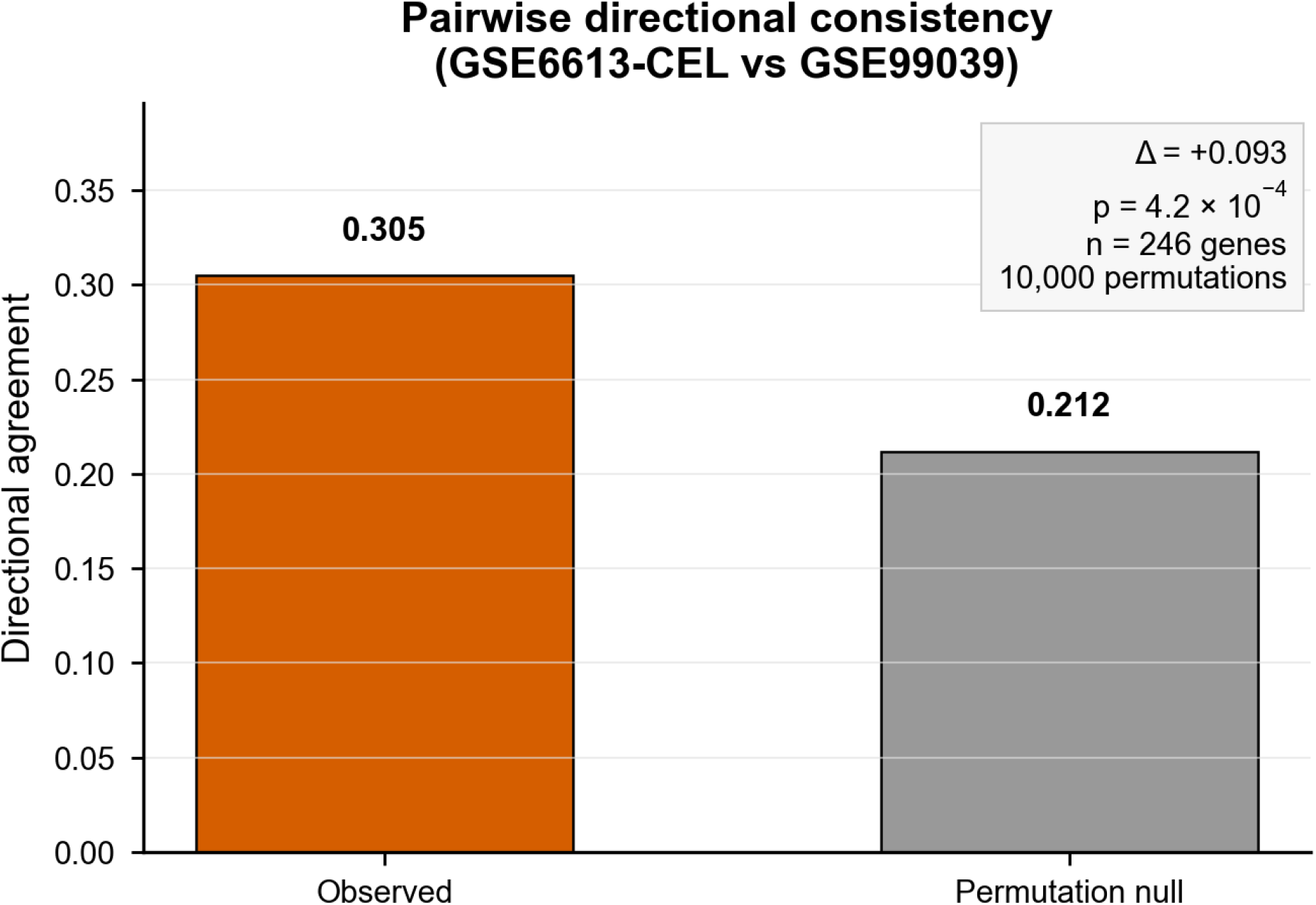
Pairwise directional consistency between GSE6613-CEL and GSE99039. Observed directional consistency (0.305) across n = 246 testable genes exceeded the permutation-derived null expectation (0.212), yielding excess agreement Δ = +0.093 (p = 4.2 × 10^−4^, 10,000 permutations). Analysis retains one-sided cases in the denominator.

**Figure 8.**
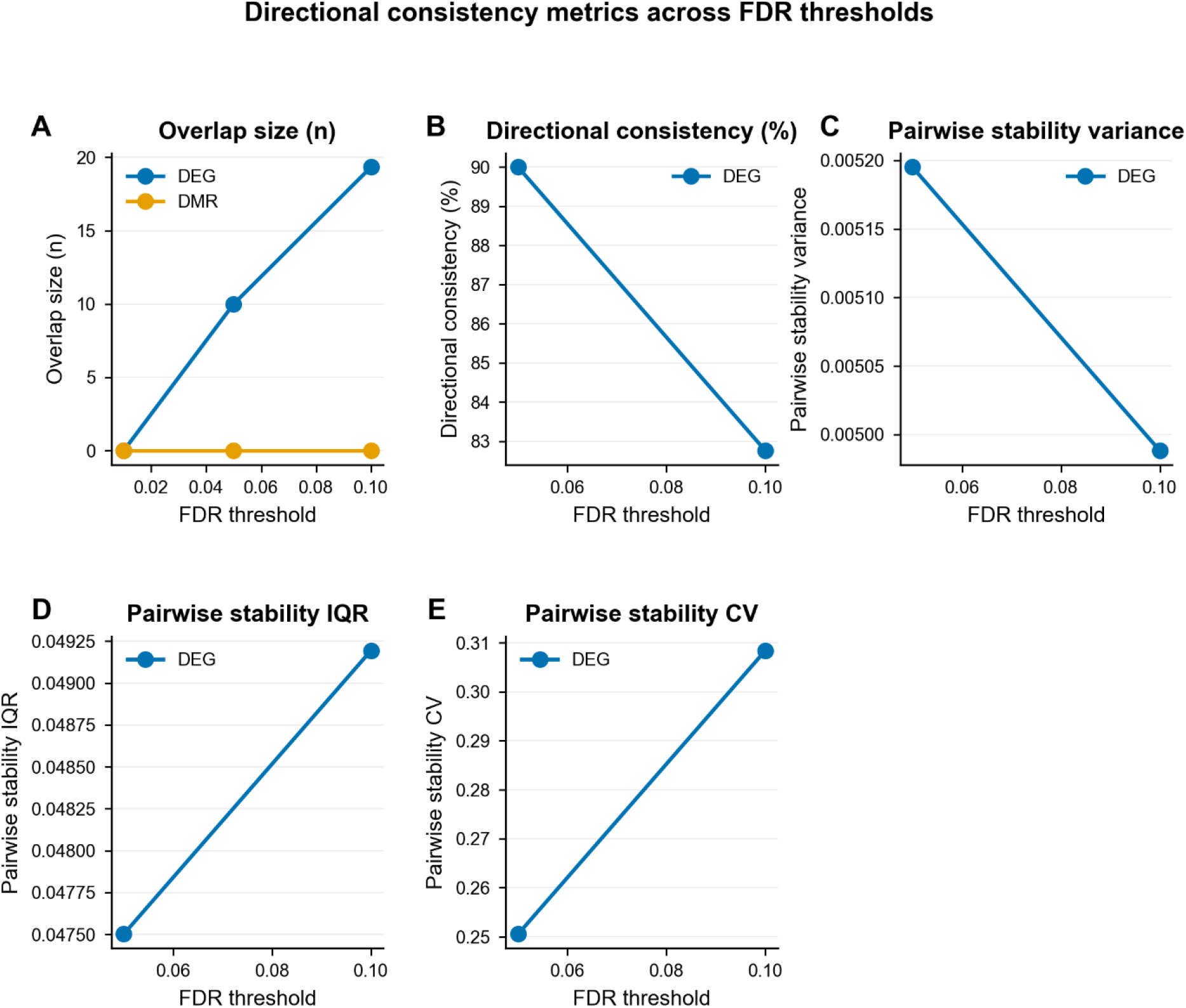
Directional consistency metrics across FDR thresholds. Six-panel sensitivity analysis of pairwise concordance statistics across FDR thresholds (0.01–0.10) for DEGs (blue) and DMRs (orange). Panels: (A) overlap size, (B) directional consistency, (C) variance, (D) interquartile range, (E) coefficient of variation. DEG overlap grows with relaxed thresholds while consistency declines modestly (90% → 83%); DMR overlap is absent across all thresholds.

In contrast, all methylation pairwise comparisons yielded zero overlapping genes meeting both false discovery rate and effect-size thresholds across FDR values of 0.01, 0.05 and 0.10. This absence of overlap, despite extreme inflation in individual datasets, underscores the structural asymmetry between within-dataset significance and cross-dataset replication in DMR-mapped analyses.

#### Gene Annotation Sensitivity

Sensitivity analyses assessing probe-to-gene collapsing strategies demonstrated comparable directional consistency across methods. Maximum absolute log_2_ fold-change selection, lowest adjusted p-value selection, and median log_2_ fold-change approaches yielded directional consistency ranging from 30.5% to 32.4% (Figure 9, Table 6). These results indicate that directional consistency metrics are robust to reasonable variations in gene annotation methodology.

**Table 6:**
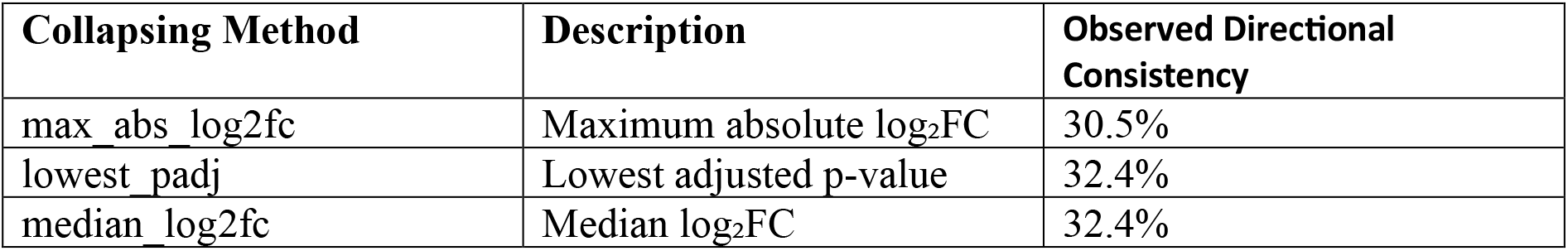
Gene annotation sensitivity analysis. Robustness of directional consistency estimates to alternative probe-to-gene collapsing strategies. All three collapsing methods produced highly comparable directional consistency values (30.5%–32.4%), confirming that results are not driven by probe annotation or collapsing choices.

| Collapsing Method | Description | Observed Directional Consistency |
| --- | --- | --- |
| max_abs_log2fc | Maximum absolute $\log_2$FC | 30.5% |
| lowest_padj | Lowest adjusted p-value | 32.4% |
| median_log2fc | Median $\log_2$FC | 32.4% |

**Table 7:**
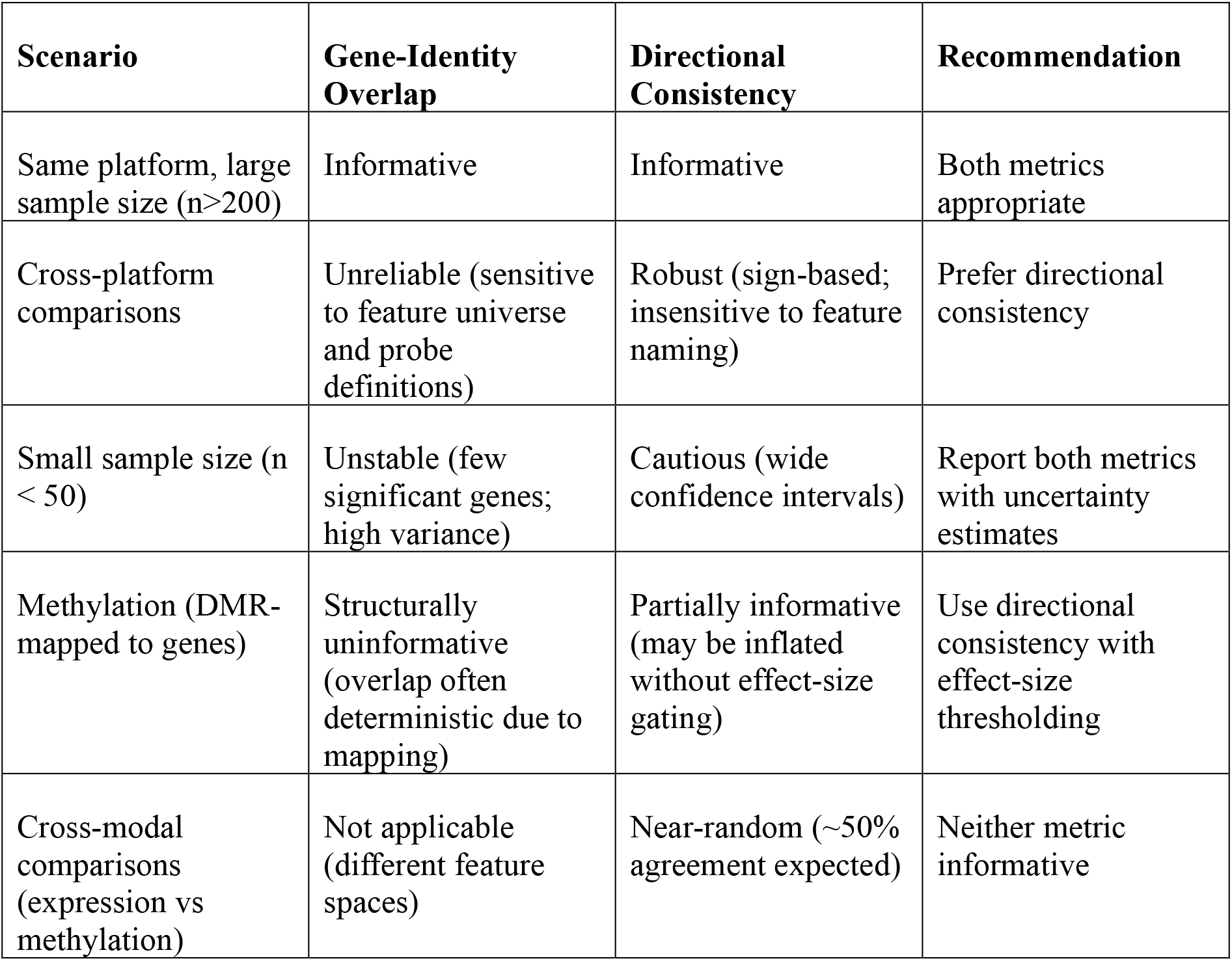
Practical guidance for selecting reproducibility metrics under common crossstudy comparison scenarios. Practical guidance for selecting reproducibility metrics under common cross-study comparison scenarios. These recommendations are derived from the empirical findings of this study and are intended as practical guardrails rather than definitive prescriptions.

**Figure 9.**
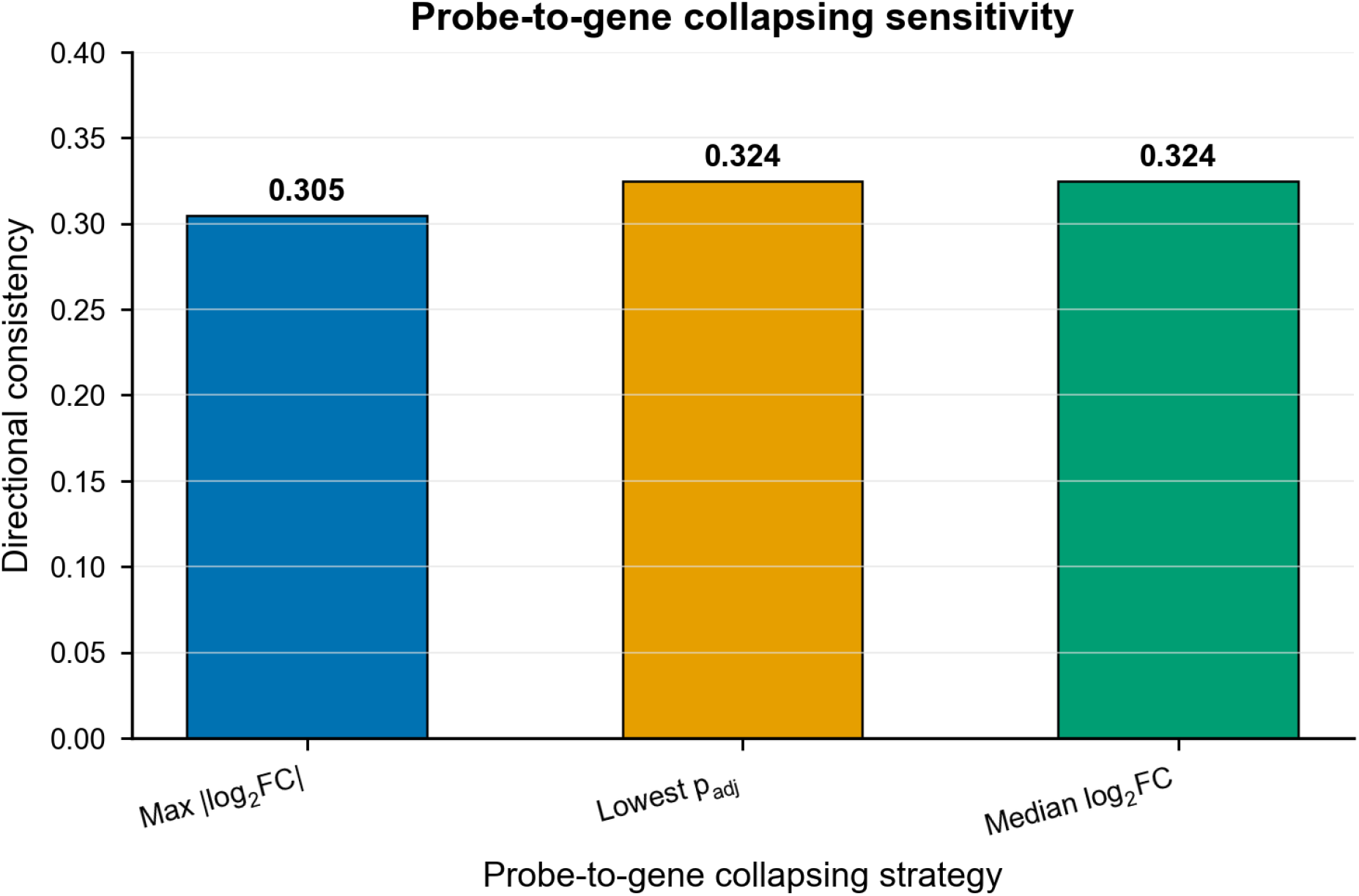
Directional consistency is robust to probe-to-gene collapsing strategy. Observed directional consistency under three collapsing methods: maximum |log_2_FC| (0.305), lowest adjusted p-value (0.324), and median log_2_FC (0.324). Narrow range confirms robustness to annotation choice.

### Cross-Modal Directional Consistency

Cross-modal comparison between expression-derived and methylation-derived gene-level effects revealed no consistent directional consistency across modalities. Agreement rates approximated random expectation (∼50%), indicating limited stability of gene-level signals across transcriptomic and epigenomic layers. These findings further underscore the modality-specific nature of reproducibility patterns.

### Meta-Analytic and Multi-Omic Diagnostics

Between-dataset heterogeneity metrics revealed substantial variability across studies, consistent with differences in platform, cohort composition, and analytical universe. Multiomic integration and pathway enrichment analyses yielded extensive apparent enrichment; however, these findings were interpreted strictly as methodological artifacts arising from large and overlapping gene universes rather than evidence of biological replication. All metaanalytic and integrative outputs are presented as descriptive diagnostics and are not interpreted as biological findings.

## DISCUSSION

This study presents a systematic methodological evaluation of reproducibility across heterogeneous transcriptomic and DNA methylation datasets. Differential expression consistency across datasets has been widely studied in the context of meta-analysis frameworks (Wang et al., 2012). Classical gene-identity overlap was frequently limited across independent cohorts, particularly when studies differed in platform technology and feature definitions (Zhang et al., 2009; Irizarry et al., 2005; MAQC Consortium, 2006). These findings highlight the inherent limitations of overlap-based metrics in cross-dataset reproducibility assessments.

Existing cross-study reproducibility frameworks such as MetaQC, MetaDE, and rank–rank hypergeometric overlap (RRHO) primarily focus on rank concordance, meta-analytic aggregation, or cross-platform harmonization (Kang et al., 2012; Wang et al., 2012; Johnson et al., 2007; Plaisier et al., 2010). In contrast, the present analysis isolates directional stability independent of magnitude ranking, providing a complementary diagnostic perspective focused strictly on effect-sign reproducibility under heterogeneous analytical conditions.

For methylation datasets, genomic inflation factors exceeding 100 reflect the dense correlation structure of CpG probes and region-based aggregation procedures rather than classical population stratification (Devlin and Roeder, 1999; Price et al., 2010). Under such conditions, raw significance counts become nearly deterministic functions of feature density and correlation structure. Importantly, directional consistency analyses rely on the sign of effect estimates rather than p-value magnitude and therefore remain mathematically well-defined even when calibration of absolute significance is compromised. Consequently, methylation overlap enrichment results are interpreted as structurally uninformative rather than as empirical evidence of replication failure.

Near-complete significance in DMR-mapped gene sets reflects the expansion of gene coverage produced by region-to-gene mapping combined with correlated CpG structure rather than independent biological signal. LambdaGC calculations assume a 1-degree-of-freedom chisquared distribution; however, region-level statistics derived from kernel smoothing procedures (e.g., DMRcate) may not strictly conform to this assumption. Consequently, λ estimates for methylation datasets should be interpreted cautiously as descriptive diagnostics rather than calibrated inflation metrics.

Notably, 16.1% of DMR-associated genes exhibited direction discordance across datasets, indicating weaker directional robustness for methylation compared to expression features. This estimate includes all genes with non-zero effects regardless of statistical significance; direction estimates for genes not reaching significance thresholds may be unreliable and could inflate apparent discordance. This further highlights structural differences between transcriptomic and epigenomic reproducibility behavior.

In contrast, directional consistency emerged as a more stable and informative metric for evaluating reproducibility under heterogeneous conditions (Sweeney et al., 2017; Cahill et al., 2018). Directional consistency remained high across independent cohorts, analytical methods, significance thresholds, and gene annotation strategies, indicating that concordance in effect direction captures reproducible statistical behavior that is less sensitive to platform-specific feature definitions (Dayan et al., 2025; Lie et al., 2025).

These findings are consistent with recent work demonstrating that even ostensibly reproducible Parkinson’s disease transcriptomic signatures exhibit limited clinical utility, with cross-dataset classification performance near chance levels (Dayan et al., 2025). Our results suggest a potential mechanistic explanation: if reproducibility assessments relied primarily on gene-identity overlap — a metric we show to be unstable across heterogeneous datasets — then signatures validated using overlap-based criteria may overestimate true replicability. Directional consistency, which we find to be more robust under the same conditions, may provide a more reliable foundation for evaluating whether molecular signatures generalize across independent cohorts.

However, rank-based concordance across independent cohorts was minimal (median Spearman ρ ≈ 0.04–0.045 across datasets for differential expression and −0.15 to 0.26 for methylation datasets). This indicates that while direction of effect is often stable, relative effect magnitude ordering is largely inconsistent across studies. Consequently, directional consistency does not imply prioritization concordance and should not be interpreted as ranking robustness.

The robustness analyses further demonstrate that directional consistency is not driven by arbitrary analytical choices. Threshold sensitivity analyses showed that directional consistency persists despite substantial increases in overlap size, while gene annotation sensitivity analyses confirmed stability across alternative probe-to-gene collapsing strategies. These findings support directional consistency as a statistically tractable reproducibility metric in multi-dataset settings.

Several limitations should be noted. Feature universes differed substantially across platforms, particularly between expression and methylation data, restricting direct cross-modal comparisons (Ritchie et al., 2015; Aryee et al., 2014; Fortin et al., 2017). Statistical power varied substantially across datasets, with some cohorts having modest sample sizes (e.g., GSE165083 n=26/28; GSE111629 yielding only 29 significant genes). Limited power may contribute to instability in overlap metrics and reduced replication probability. GSE6613-CEL exhibited a high proportion of significant features (37.0%) alongside substantial genomic inflation (λGC = 5.03), which may reflect platform-specific signal characteristics and should be interpreted cautiously. Additionally, inferential directional analyses necessarily included methodological replications within the same cohort, limiting conclusions to statistical reproducibility rather than independent biological validation.

Estimated proportions of major blood cell types were computed using EpiDISH where applicable but were not included as covariates in differential analyses (Andersen et al., 2023; Houseman et al., 2012; Newman et al., 2015). This represents a substantive limitation for methylation results specifically: blood DNA methylation profiles are strongly influenced by inter-individual variation in immune cell-type composition (Houseman et al., 2012), and unadjusted DMR analyses may capture cell-proportion differences between PD cases and controls rather than disease-specific epigenetic changes. As the objectives of this study are methodological rather than biological, re-running cell-composition-adjusted models was outside the scope of the current analysis and would not alter the primary conclusions. Additionally, sex chromosome probes were retained in methylation analyses; sex was included as a covariate where available to mitigate potential confounding, but residual sex-linked effects cannot be excluded.

All datasets were derived from peripheral blood samples. As Parkinson’s disease is a neurodegenerative disorder primarily affecting the central nervous system, peripheral molecular signatures may not fully reflect brain-specific pathophysiology. This limitation does not affect the methodological conclusions but restricts biological interpretation.

Importantly, this study does not aim to identify Parkinson’s disease biomarkers, pathways, or molecular mechanisms (Kurvits et al., 2021; Scherzer et al., 2007; Shamir et al., 2017). All analyses are explicitly methodological and are intended to characterize the statistical behavior of reproducibility metrics under heterogeneous analytical conditions. Simulation studies were not performed because the goal was not to validate a new estimator under controlled conditions, but to assess real-world behavior under the heterogeneity that motivates the need for diagnostic guardrails.

## CONCLUSION

Gene-identity overlap alone provides an unstable and often misleading measure of reproducibility across heterogeneous molecular datasets. Directional consistency offers a more robust alternative that remains stable across platforms, analytical thresholds, and annotation strategies. All conclusions drawn in this study are methodological in nature and do not constitute claims regarding Parkinson’s disease biology.

## Supporting information

Supplementary image files

## ACKNOWLEDGMENTS

The authors would like to express their sincere gratitude to the DAV Managing Committee, Delhi and the Department of Biotechnology, DAV College, Sector 10, Chandigarh for providing the necessary facilities, resources and support to carry out this work.

## AUTHORS’ CONTRIBUTIONS

Inayat Chauhan designed the study, acquired raw data, and prepared the initial draft of the manuscript. Chiragh Dewan performed the bioinformatics analysis. Kanika Sharma provided valuable inputs for compiling the manuscript, editing, and critical review of the study. Sangeeta Sharma and Rupinderjeet Kaur supervised the study.

## DATA, SCRIPT, CODE, AND SUPPLEMENTARY INFORMATION AVAILABILITY

All analysis scripts were executed in R (version 4.5.1) and Python (version 3.13.7). Reproducible workflows and configuration files are [PLACEHOLDER: Scripts to be deposited on Zenodo prior to submission. Replace this text with: Scripts available at https://doi.org/10.5281/zenodo.XXXXXXX (Zenodo DOI to be added before submission)]. All analyses were performed with fixed random seeds (seed = 42) to ensure reproducibility.

## SUPPLEMENTARY MATERIALS

Supplementary figures and tables provide detailed quality control diagnostics, calibration assessments, direction-stratified overlap statistics, and robustness analyses that support the main methodological conclusions of the study. These materials are intended to document analytical validity and parameter sensitivity rather than biological interpretation.

**Supplementary Table 1:**
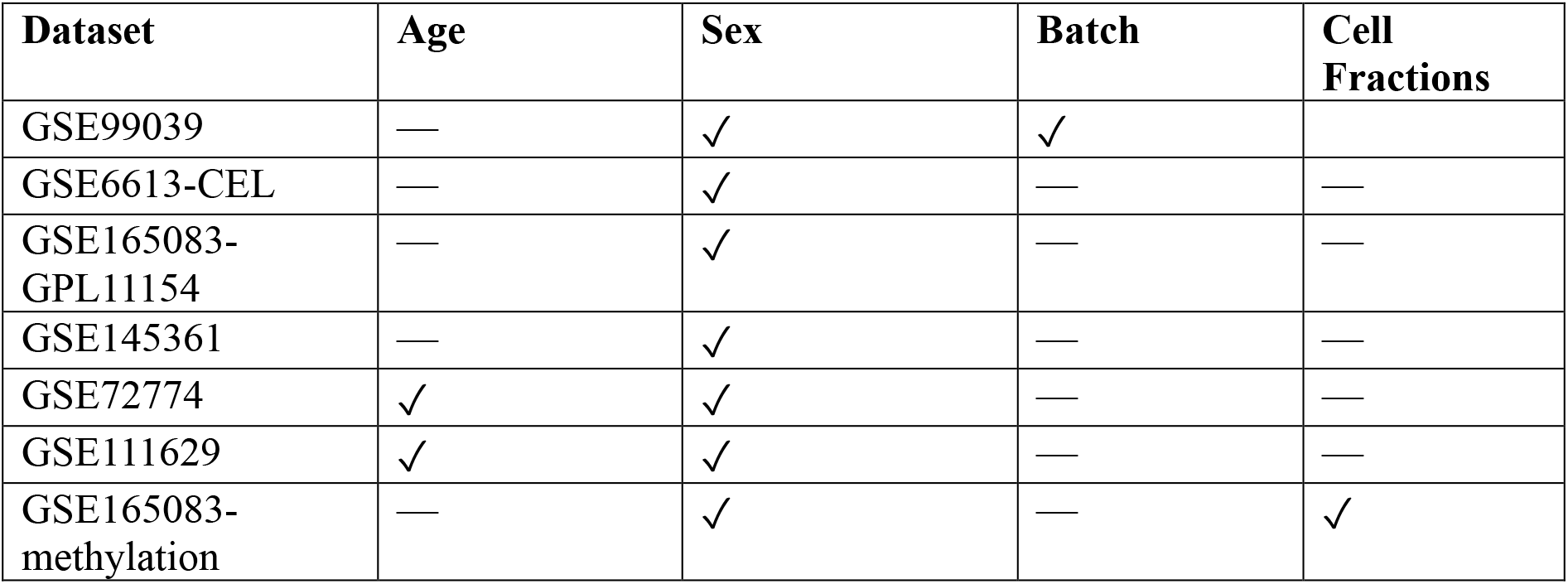
Covariate Availability Across Datasets. Availability of demographic and technical covariates. ✓ indicates valid numeric covariate values present for the majority of samples. — indicates absent or missing values. Cell-type fraction estimates (EpiDISH-derived) were available for GSE165083-methylation only but were not included as covariates in any differential analysis.

**Supplementary Table 2:**
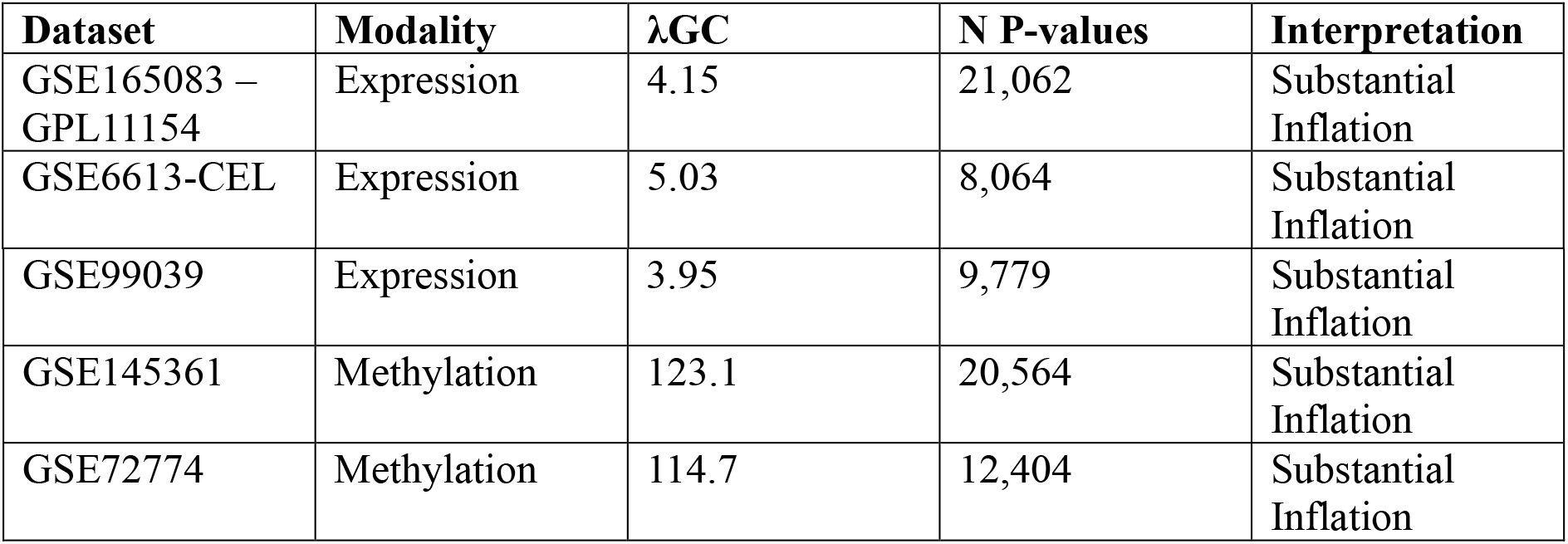
Genomic Inflation Factors and Calibration Diagnostics. For methylation datasets, region-level λGC values are shown; CpG-level λGC values (1.04 and 0.99 respectively) are reported in the main text. GSE111629 and GSE165083-methylation produced insufficient region-level p-values for stable λGC estimation and are omitted from region-level diagnostics.

| Dataset | Modality | $\lambda_{GC}$ | N P-values | Interpretation |
| --- | --- | --- | --- | --- |
| GSE165083 – GPL11154 | Expression | 4.15 | 21,062 | Substantial Inflation |
| GSE6613-CEL | Expression | 5.03 | 8,064 | Substantial Inflation |
| GSE99039 | Expression | 3.95 | 9,779 | Substantial Inflation |
| GSE145361 | Methylation | 123.1 | 20,564 | Substantial Inflation |
| GSE72774 | Methylation | 114.7 | 12,404 | Substantial Inflation |

**Supplementary Table 3:**
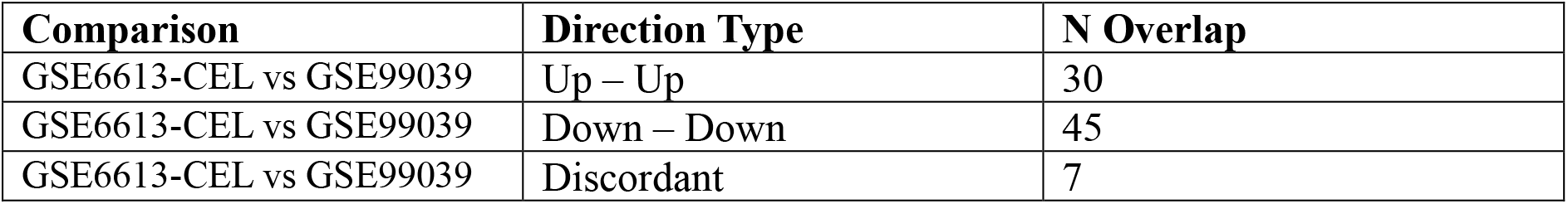
Direction-Stratified Overlap Counts. Counts of overlapping features stratified by direction of effect for pairwise dataset comparisons. These counts underlie the directional consistency metrics reported in the main text.

**Supplementary Table 4:**
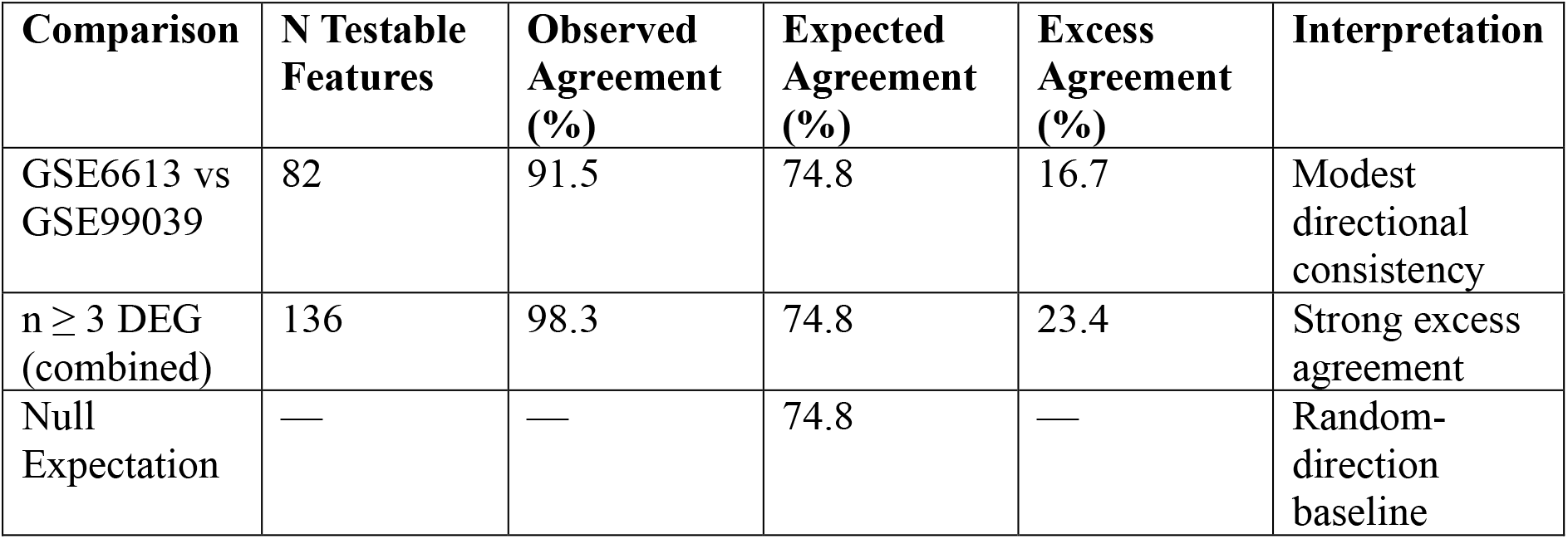
Pairwise Null-Model Context for Directional Consistency. Permutation-based null-model statistics. Expected agreement was derived from permutation procedures using fixed random seeds and 10,000 iterations.

Supplementary Figures S1–S12: Quality control diagnostics, calibration QQ plots, gene-mapping expansion plots, distributional properties of directional consistency metrics, direction-stratified overlap heatmaps, and rank concordance heatmaps (to be provided as separate figure files).

