## Supplementary material for "Directional Gene-Level Concordance and Methodological Constraints in Blood Transcriptomic and DNA Methylation Studies of Parkinson’s Disease": Code files: PD_paper_PCI_MCB_final.docx

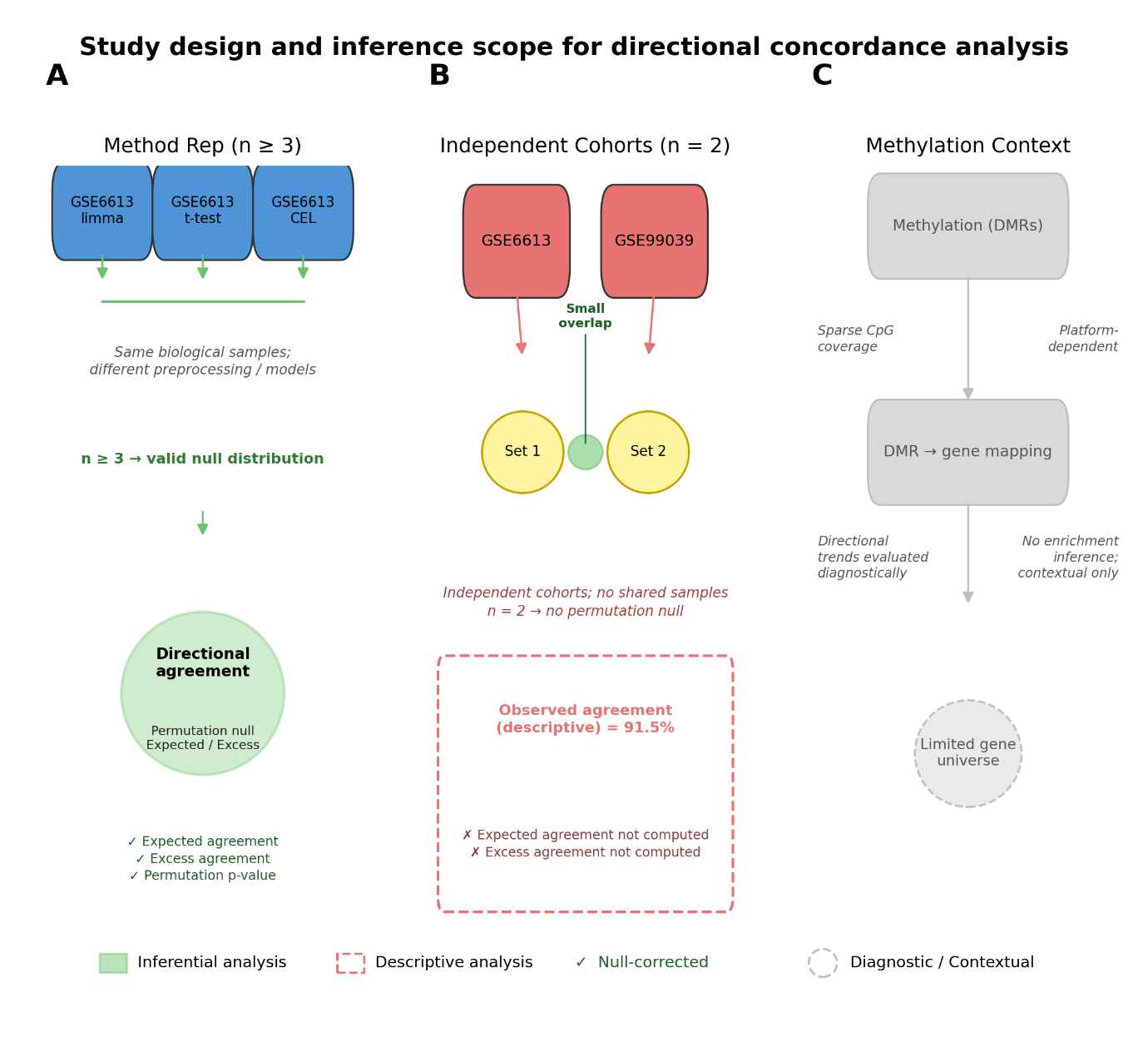
*Figure 1. Study design and inference scope for directional consistency analysis. Schematic overview of the reproducibility framework. (A) Method-replication analysis (n ≥ 3 datasets) using permutation-based inference to estimate excess directional consistency. (B) Independent cohort comparison (n = 2) reporting descriptive directional consistency only. (C) Contextual considerations for methylation data, including probe-to-gene mapping constraints and platform-dependent annotation effects.*

**Data Selection and Characteristics**

A total of seven datasets were analyzed, derived from five biologically independent cohorts (GSE6613 and GSE6613-CEL share underlying samples). These included three transcriptomic expression datasets and four DNA methylation datasets generated using Affymetrix and Illumina platforms. Expression datasets included GSE6613, GSE99039, and GSE165083 (GPL11154), while methylation datasets included GSE111629, GSE72774, GSE145361, and GSE165083 (methylation platform). All datasets were derived from peripheral blood samples.

For example, GSE6613-CEL included 25 Parkinson’s disease cases and 37 controls (n = 62), whereas GSE99039 comprised 191 cases and 212 controls (n = 403). The largest methylation dataset, GSE145361, included 1,889 samples (959 cases and 930 controls) and interrogated approximately 485,512 CpG loci. Unless otherwise specified, all references to GSE6613 in comparative analyses refer to the GSE6613-CEL pipeline for consistency. Quality control considerations for cross-study integration were informed by established meta-analysis frameworks (Kang et al., 2012). Feature counts reflect post-quality-control analytical universes and therefore varied across platforms and datasets.

| **Dataset** | **Modality** | **Platform** | **PD Samples** | **Control Samples** | **Total** | **Features Analyzed** |
| --- | --- | --- | --- | --- | --- | --- |
| GSE99039 | Expression | Affymetrix | 191 | 212 | 403 | 9,779 |
| GSE6613-CEL | Expression | Affymetrix | 25 | 37 | 62 | 8,064 |
| GSE165083-GPL11154 | Expression | RNA-seq | 12 | 14 | 26 | 21,062 |
| GSE145361 | Methylation | Illumina 450K | 959 | 930 | 1,889 | 485,512 |
| GSE72774 | Methylation | Illumina 450K | 289 | 219 | 508 | 485,512 |
| GSE111629 | Methylation | Illumina 450K | 334 | 237 | 571 | 393,217 |
| GSE165083 | Methylation | Illumina 450K | 14 | 14 | 28 | 485,512 |
| Total |  |  | 1,824 | 1,663 | 3,487 |  |

**Table 1: Summary of included datasets**

*Table 1: Characteristics of seven peripheral blood transcriptomic and DNA methylation datasets analyzed, including platform, sample counts, and features retained after preprocessing. In total, 3,487 samples (1,824 PD and 1,663 controls) were included. GSE6613-CEL represents the same underlying cohort as GSE6613, reprocessed from raw CEL files using an alternative analytical pipeline. GSE99039 metadata contained 557 entries; 119 samples with unknown disease status were excluded, and a further 35 samples were removed during quality control. GSE6613-CEL metadata contained 105 samples; 33 samples labelled ‘Neurological Control’ were merged with 22 regular controls, and 43 samples were excluded during quality control.*

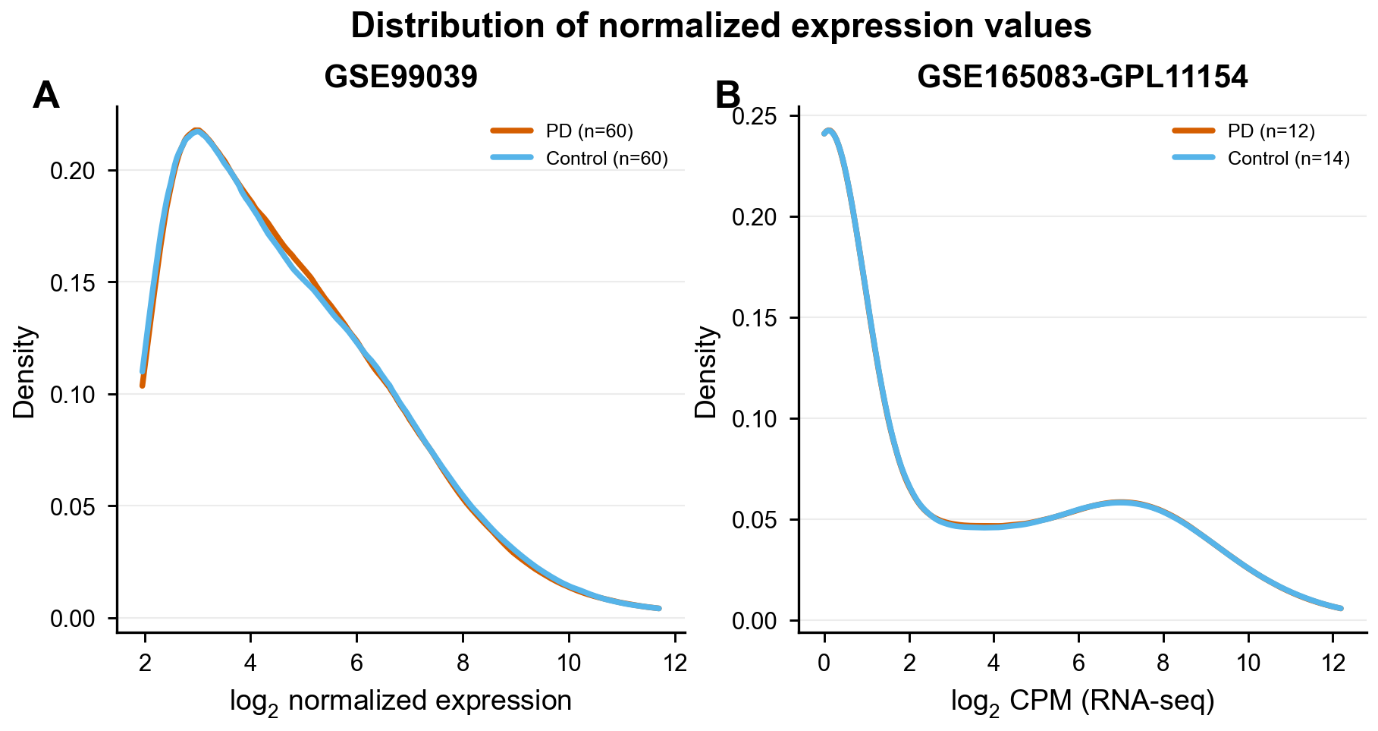

*Figure 2. Distribution of normalized expression values. Density plots of normalized expression values for representative microarray (GSE99039) and RNA-seq (GSE165083-GPL11154) datasets. Thin lines show per-sample densities (subsampled); thick lines show within-group medians (PD = vermillion, Control = sky blue).*

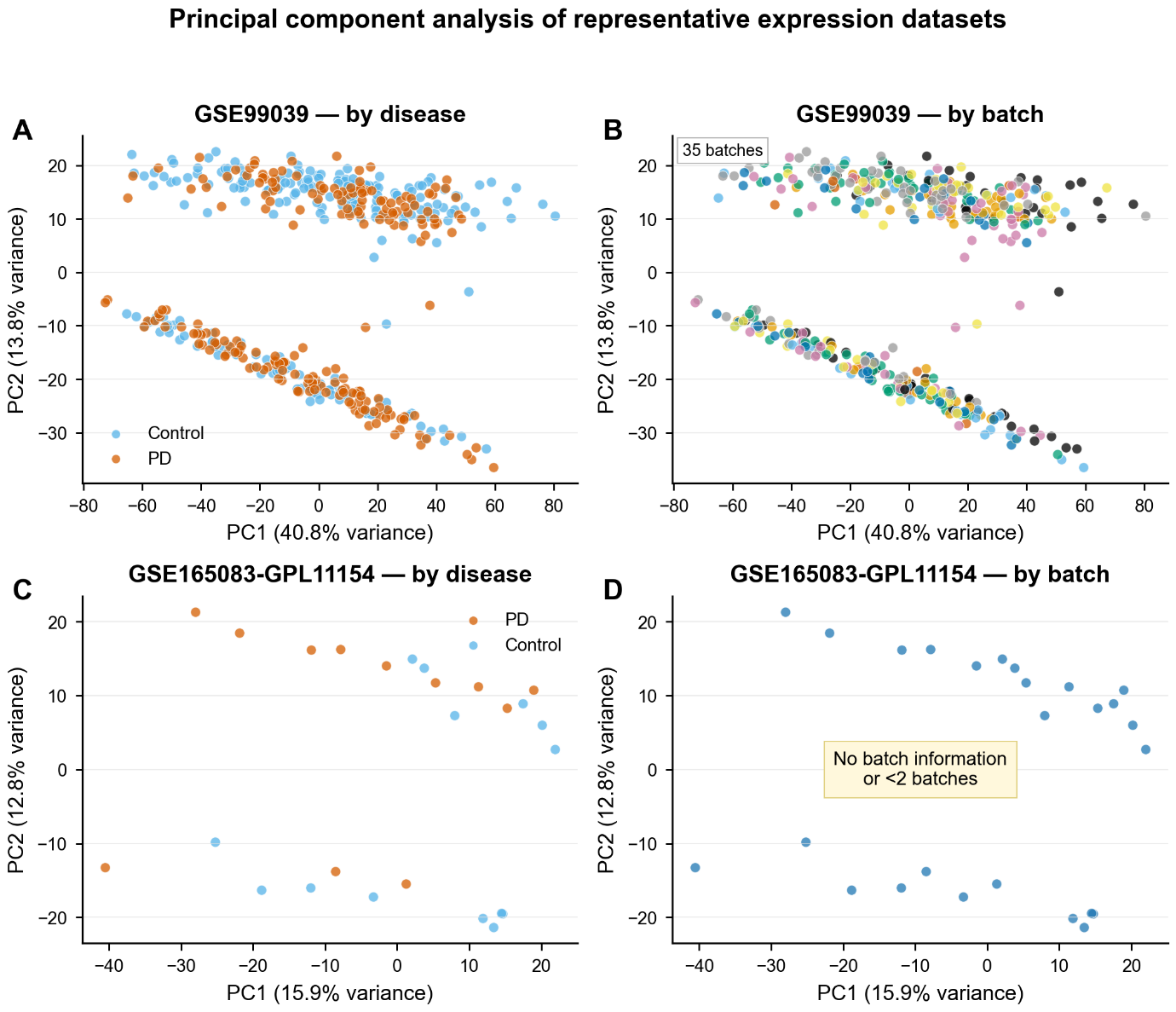

*Figure 3. Principal component analysis of representative expression datasets. PCA of top-1000 variance features in GSE99039 and GSE165083-GPL11154. Samples are colored by disease status (left column) and batch (right column, where available). Axes indicate variance explained. Disease-related separation is modest relative to overall variance structure.*

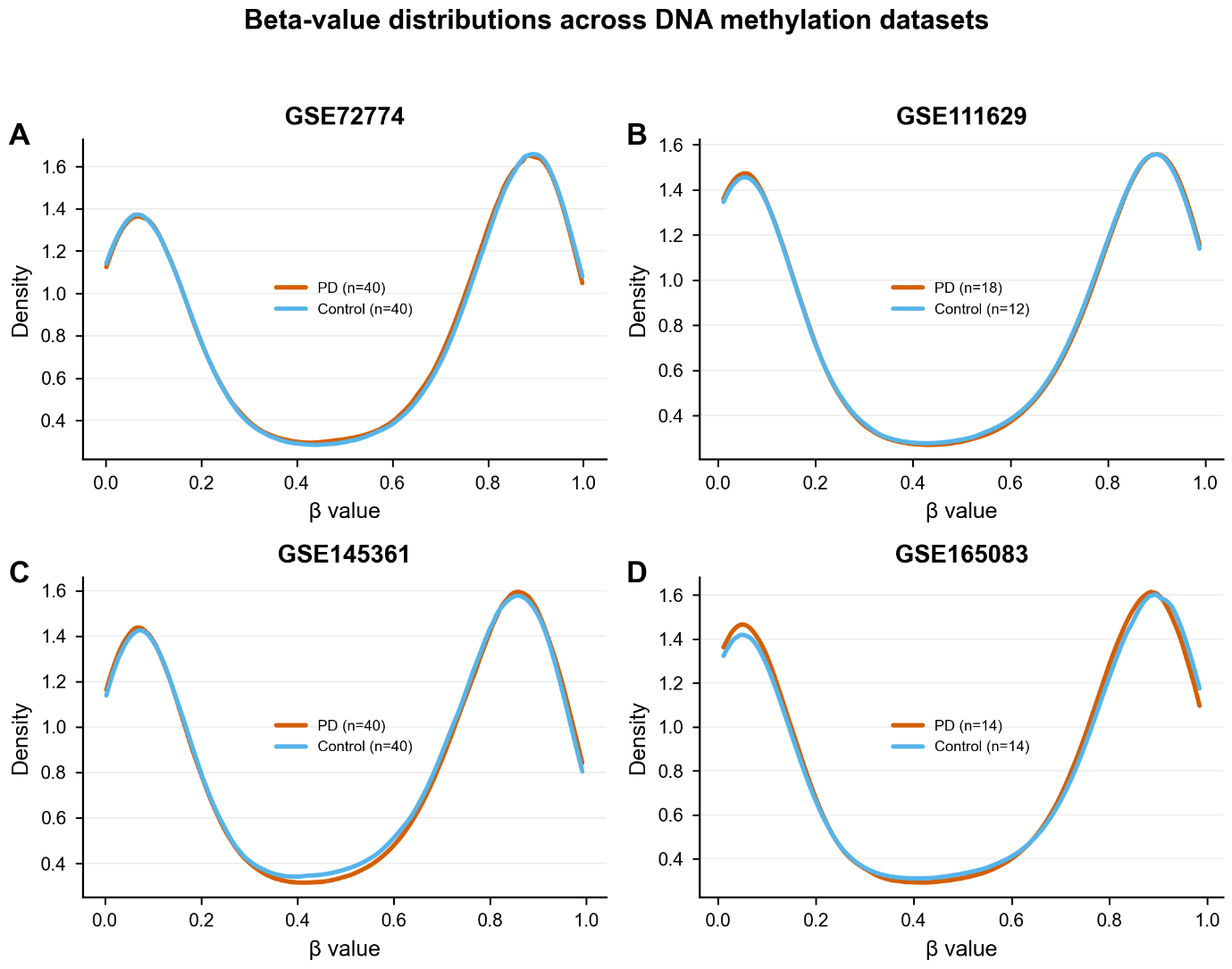

*Figure 4. Beta-value distributions across DNA methylation datasets. Density plots showing distribution of CpG methylation β values across all four methylation datasets. Thin lines show per-sample densities; thick lines show within-group medians. All datasets exhibit the characteristic bimodal distribution of methylation levels.*

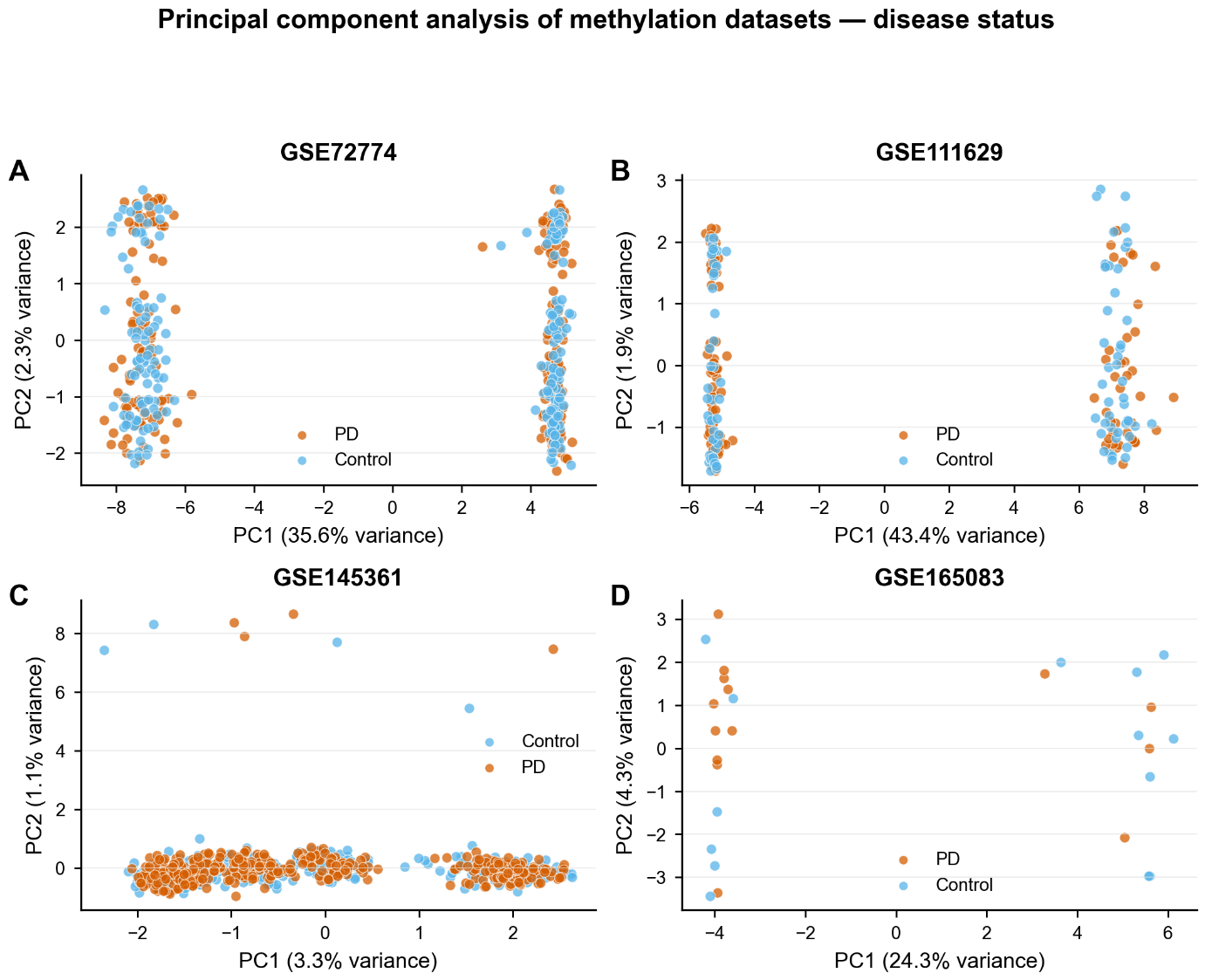

*Figure 5a. PCA of methylation datasets — disease status. PCA based on top-1000 variance CpGs within each dataset. Samples colored by PD (vermillion) vs Control (sky blue). Dataset-local; not directly comparable across cohorts.*

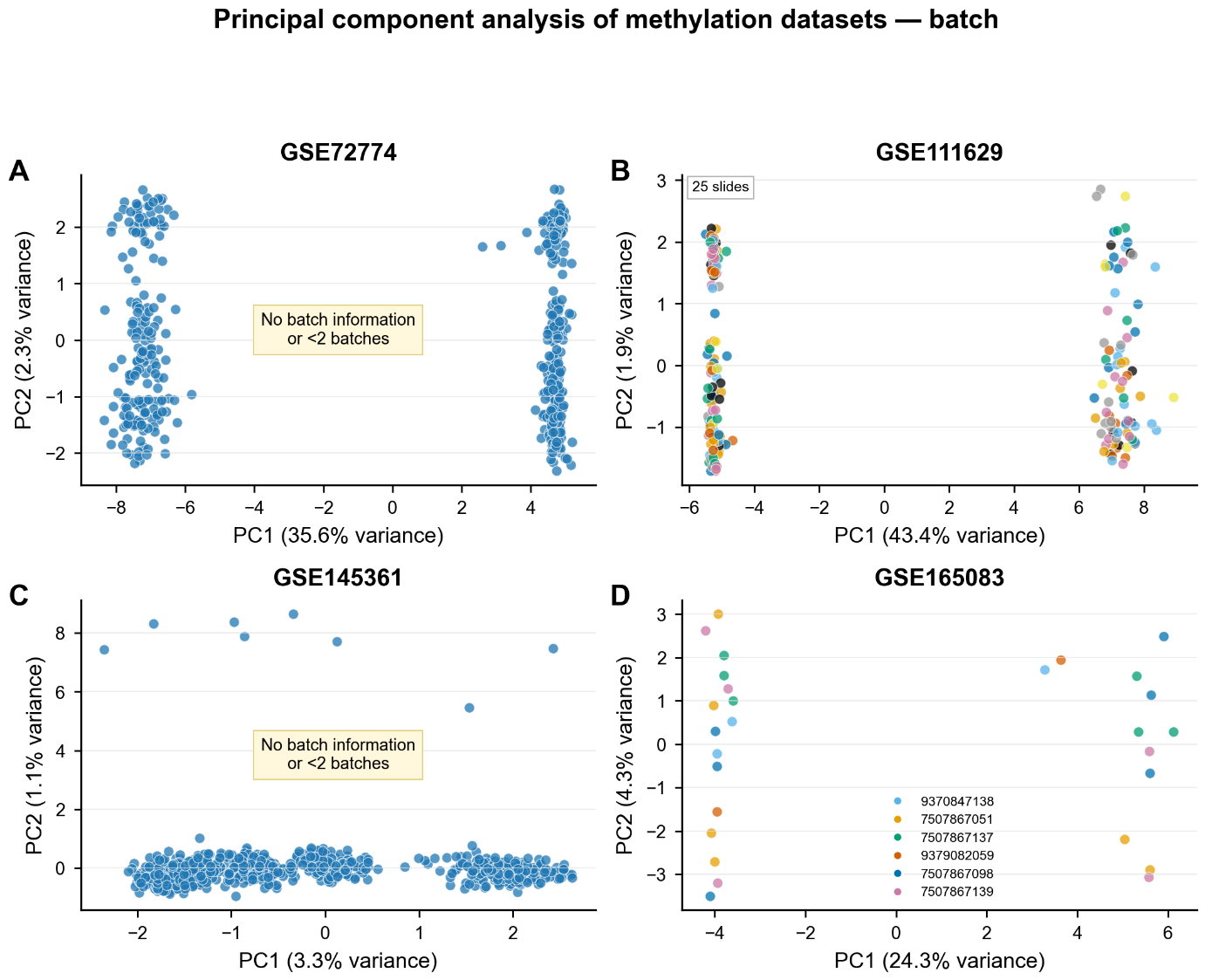

*Figure 5b. PCA of methylation datasets — batch. Same PCA projections as Figure 5a, with samples colored by Sentrix slide ID where available.*

**Differential Analysis Summary**

Differential expression and methylation analyses identified variable numbers of statistically significant features across datasets, reflecting differences in sample size, platform technology, and analytical universe. Expression datasets identified between 0 and 1,596 significant features, depending on dataset and analytical pipeline, whereas methylation datasets yielded substantially larger sets of differentially methylated region (DMR)-mapped genes, frequently exceeding 18,000 genes per dataset (Table 2) (Vallerga et al., 2020; Lie et al., 2025). These disparities underscore the structural asymmetry between expression-based and methylation-based feature definitions and motivated subsequent analyses focused on reproducibility metrics that are robust to heterogeneous feature universes.

The OLS regression framework was informed by the limma methodology (Ritchie et al., 2015; Smyth, 2004) but was implemented independently in Python using statsmodels; it does not call the R limma package directly.

| **Dataset** | **Modality** | **DEGs at FDR<0.10** | **DEGs at FDR<0.05** | **Proportion at FDR<0.10** | **Primary FDR Cutoff** |
| --- | --- | --- | --- | --- | --- |
| GSE99039 | DEG | 2,670 | 1,596 | 59.8% | 0.10 |
| GSE6613-CEL | DEG | 2,104 | 779 | 37.0% | 0.10 |
| GSE6613 (t-test) | DEG | 5 | 0 | 0% | 0.10 |
| GSE72774 | DMR | 9,268 | 9,266 | 99.98% | 0.05 |
| GSE145361 | DMR | 12,917 | 12,915 | 99.98% | 0.05 |
| GSE111629 | DMR | 29 | 29 | 100% | 0.05 |
| GSE165083-GPL11154 | DEG | 0 | 0 | 0% | 0.05 |

**Table 2: Differential analysis summary per dataset**

*Table 2: Post-quality-control analytical universes, number of statistically significant features, and corresponding FDR thresholds. Differences in the proportion of significant features reflect platform-specific feature definitions rather than biological interpretation. The GSE6613 (t-test) row is a methodological diagnostic comparison included for illustrative comparison only.*

**Overlap Enrichment Analysis**

Within-dataset methodological comparisons demonstrated strong overlap enrichment, indicating internal analytical consistency. In contrast, independent cohort comparisons exhibited enrichment ratios generally close to unity, reflecting limited gene-identity replication across datasets. Methylation datasets showed near-complete overlap across comparisons, attributable to the expansive and deterministic nature of DMR-based gene mappings rather than shared biological signal (Figure 6, Table 3) (Henderson-Smith et al., 2019; Moore et al., 2014; Du et al., 2010). Although some overlaps reached nominal statistical significance, enrichment ratios were generally close to unity.

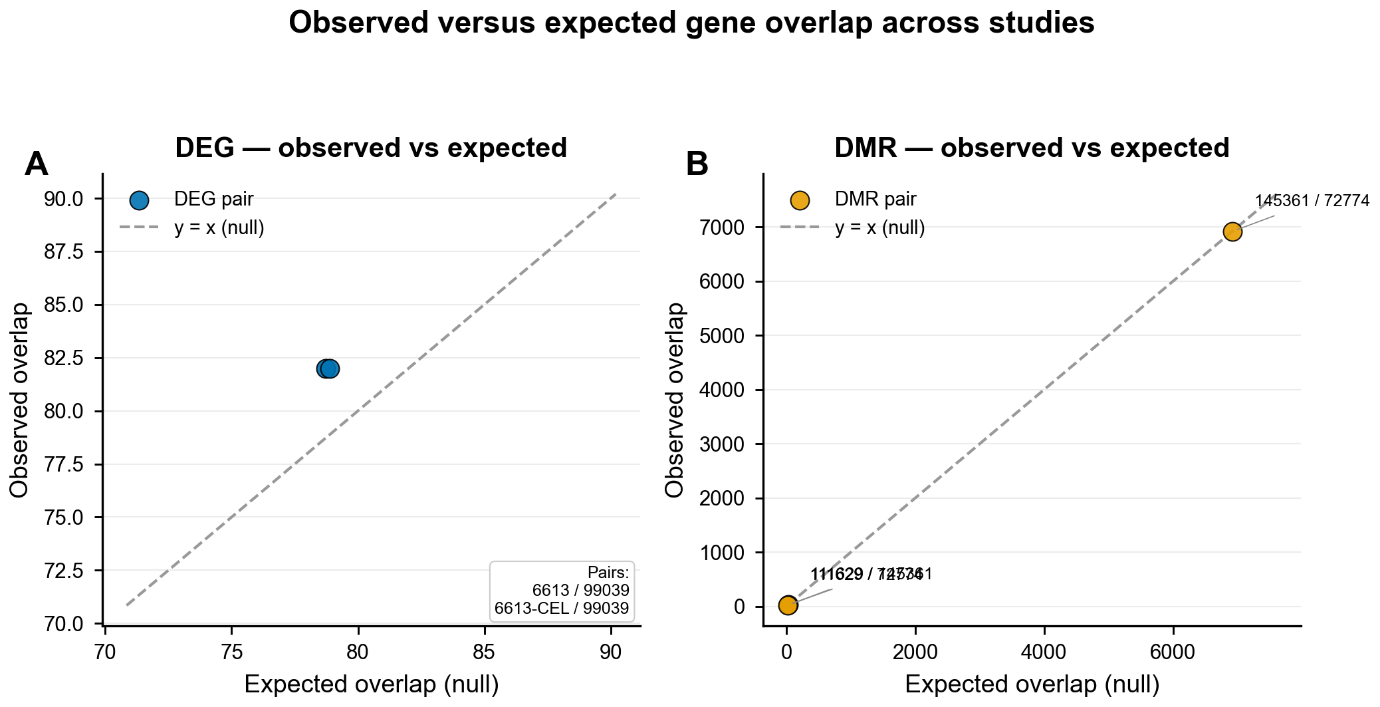

*Figure 6. Observed versus expected gene overlap across studies. Scatter plots comparing observed overlap of significant features to null-expected overlap derived from permutation-based models. (A) DEG overlap. (B) DMR overlap. Dashed line: y = x (null expectation). Methylation overlaps approach near-complete gene coverage due to DMR-to-gene mapping expansion.*

| **Comparison** | **Modality** | **Observed Overlap** | **Expected Overlap** | **Enrichment Ratio** | **Interpretation** |
| --- | --- | --- | --- | --- | --- |
| GSE6613 vs GSE6613-CEL | DEG | 779 | 289 | 2.70 | Methodological replication |
| GSE6613 vs GSE99039 | DEG | 82 | 79 | 1.04 | Near-null (independent) |
| GSE6613-CEL vs GSE99039 | DEG | 82 | 79 | 1.04 | Near-null (independent) |
| GSE145361 vs GSE72774 | DMR | 6,917 | 6,917 | 1.00 | Deterministic (artifact) |
| GSE111629 vs GSE145361 | DMR | 28 | 28 | 1.00 | Deterministic (artifact) |
| GSE111629 vs GSE72774 | DMR | 19 | 19 | 1.00 | Deterministic (artifact) |

**Table 3: Overlap enrichment analysis**

*Table 3: Observed and expected overlaps computed using dataset-specific feature universes. Enrichment ratios near unity indicate limited gene-identity replication across independent cohorts. Deterministic overlaps observed for methylation datasets reflect large and highly overlapping DMR-mapped gene universes rather than shared biological signal.*

**Directional Consistency Analysis**

Directional consistency analyses demonstrated substantially greater stability than gene-identity overlap. Alternative analytical methods applied to the same dataset showed complete directional consistency among significant features. Descriptive comparisons between independent cohorts also demonstrated high directional consistency (Table 4), despite limited overlap at the gene-identity level.

Inferential analysis involving at least three analytical results (including alternative pipelines applied to the same cohort) revealed significant excess directional consistency beyond random expectations. Because these comparisons combine two independent cohorts with one methodological reanalysis, they reflect analytical stability rather than three independent biological replications and should be interpreted accordingly.

Permutation testing confirmed that observed directional consistency significantly exceeded null expectations in all evaluable comparisons (aggregate permutation p < 0.001 based on 10,000 permutations). These p-values reflect aggregate directional consistency statistics across shared gene sets, not per-gene permutation tests, and support the statistical robustness of the directional consistency metric at the dataset level.

Directional consistency is interpreted exclusively as a measure of analytical stability and not as evidence of shared disease mechanisms or etiological consistency (Dayan et al., 2025; Cahill et al., 2018). Directional consistency reflects consistency in effect sign and does not imply concordance in effect magnitude or biological relevance.

| **Comparison** | **N Testable** | **Observed Agreement** | **Expected (Null)** | **Excess Agreement** |
| --- | --- | --- | --- | --- |
| GSE6613-CEL vs GSE99039 | 82 | 91.5% (95% CI: 83.4–95.8%) | 74.8% | 16.7% |
| n ≥3 DEG (combined) | 136 | 98.3% (95% CI: 94.8–99.6%) | 74.8% | 23.4% |

**Table 4: Directional consistency analysis results**

*Table 4: DEG, differentially expressed gene. Directional consistency was defined as concordant direction of effect (up–up or down–down) across datasets. Expected agreement was estimated from permutation-based null models. The n ≥3 dataset analysis includes two analytical pipelines applied to the same underlying cohort and therefore reflects analytical stability rather than independent biological replication. The two rows use different analytical frameworks and are not directly comparable.*

Because GSE6613 and GSE6613-CEL represent alternative analytical pipelines applied to the same underlying biological cohort, the n ≥ 3 directional stability estimate (98.3%) is partially driven by non-independence. A gene identified in both pipelines does not constitute independent biological replication. Therefore, this estimate reflects methodological robustness rather than three independent cohort confirmations.

When evaluated across the full shared gene universe without conditioning on joint statistical significance, directional consistency between GSE6613-CEL and GSE99039 was 30.5% (95% CI: 25.1–36.5%) (Table 7). The higher 91.5% estimate reflects conditioning on genes meeting significance criteria in both datasets and therefore represents a restricted intersection rather than a fixed-universe concordance rate.

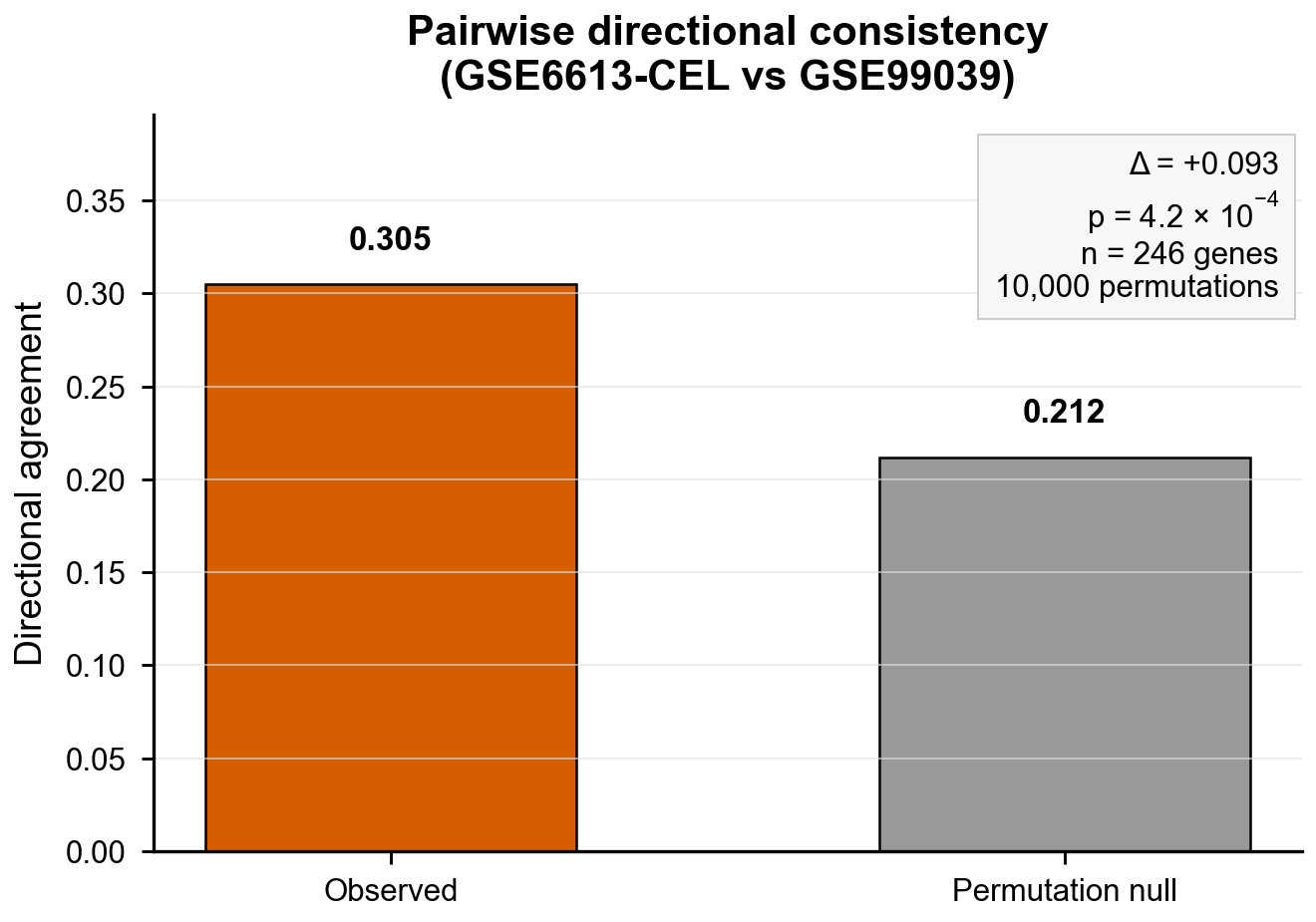

*Figure 7. Pairwise directional consistency between GSE6613-CEL and GSE99039. Observed directional consistency (0.305) across n = 246 testable genes exceeded the permutation-derived null expectation (0.212), yielding excess agreement Δ = +0.093 (p = 4.2 × 10⁻⁴, 10,000 permutations). Analysis retains one-sided cases in the denominator.*

**Robustness and Sensitivity Analyses**

Permutation-based resampling confirmed that observed directional consistency exceeded random expectation under null models with appropriate Type I error control. Directional consistency metrics remained stable across a wide range of analytical perturbations, including threshold variation and annotation strategy changes.

***Threshold Sensitivity***

Directional consistency remained high across multiple false discovery rate thresholds. For independent cohort comparisons between GSE6613-CEL and GSE99039, concordance was 90.0% at FDR < 0.05 (30 overlapping genes) and 82.8% at FDR < 0.10 (58 overlapping genes) (Figure 8, Table 5). Although overlap size increased with threshold relaxation, the persistence of high directional consistency indicates that concordance patterns are not driven by arbitrary threshold selection.

| **FDR Threshold** | **Overlap Size** | **Directional consistency** | **Change from FDR 0.05** |
| --- | --- | --- | --- |
| 0.01 | 0 | — | — |
| 0.05 | 30 | 90.0% (95% CI: 74.4–96.5%) | baseline |
| 0.10 | 58 | 82.8% (95% CI: 71.1–90.4%) | -7.2% |

**Table 5: Threshold sensitivity of directional consistency**

*Table 5: Sensitivity of directional consistency to FDR threshold selection for the independent cohort comparison. Relaxation of the FDR threshold from 0.05 to 0.10 resulted in near doubling of overlapping features (30 to 58), while directional consistency decreased modestly. No overlapping features were observed at the most stringent threshold (FDR < 0.01).*

In contrast, all methylation pairwise comparisons yielded zero overlapping genes meeting both false discovery rate and effect-size thresholds across FDR values of 0.01, 0.05 and 0.10. This absence of overlap, despite extreme inflation in individual datasets, underscores the structural asymmetry between within-dataset significance and cross-dataset replication in DMR-mapped analyses.

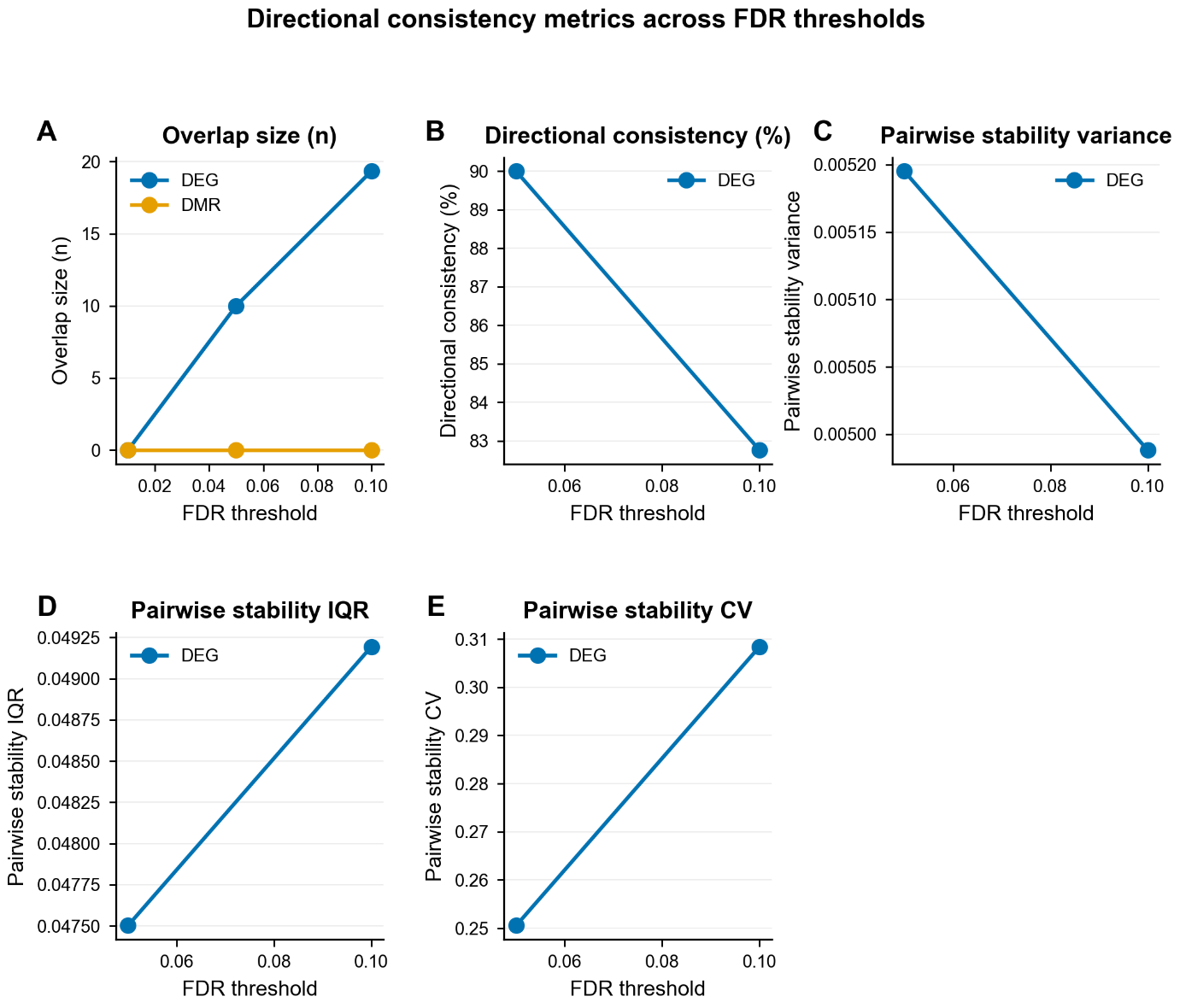

*Figure 8. Directional consistency metrics across FDR thresholds. Six-panel sensitivity analysis of pairwise concordance statistics across FDR thresholds (0.01–0.10) for DEGs (blue) and DMRs (orange). Panels: (A) overlap size, (B) directional consistency, (C) variance, (D) interquartile range, (E) coefficient of variation. DEG overlap grows with relaxed thresholds while consistency declines modestly (90% → 83%); DMR overlap is absent across all thresholds.*

***Gene Annotation Sensitivity***

Sensitivity analyses assessing probe-to-gene collapsing strategies demonstrated comparable directional consistency across methods. Maximum absolute log₂ fold-change selection, lowest adjusted p-value selection, and median log₂ fold-change approaches yielded directional consistency ranging from 30.5% to 32.4% (Figure 9, Table 6). These results indicate that directional consistency metrics are robust to reasonable variations in gene annotation methodology.

| **Collapsing Method** | **Description** | **Observed Directional Consistency** |
| --- | --- | --- |
| max_abs_log2fc | Maximum absolute log₂FC | 30.5% |
| lowest_padj | Lowest adjusted p-value | 32.4% |
| median_log2fc | Median log₂FC | 32.4% |

**Table 6: Gene annotation sensitivity analysis**

*Table 6: Robustness of directional consistency estimates to alternative probe-to-gene collapsing strategies. All three collapsing methods produced highly comparable directional consistency values (30.5%–32.4%), confirming that results are not driven by probe annotation or collapsing choices.*

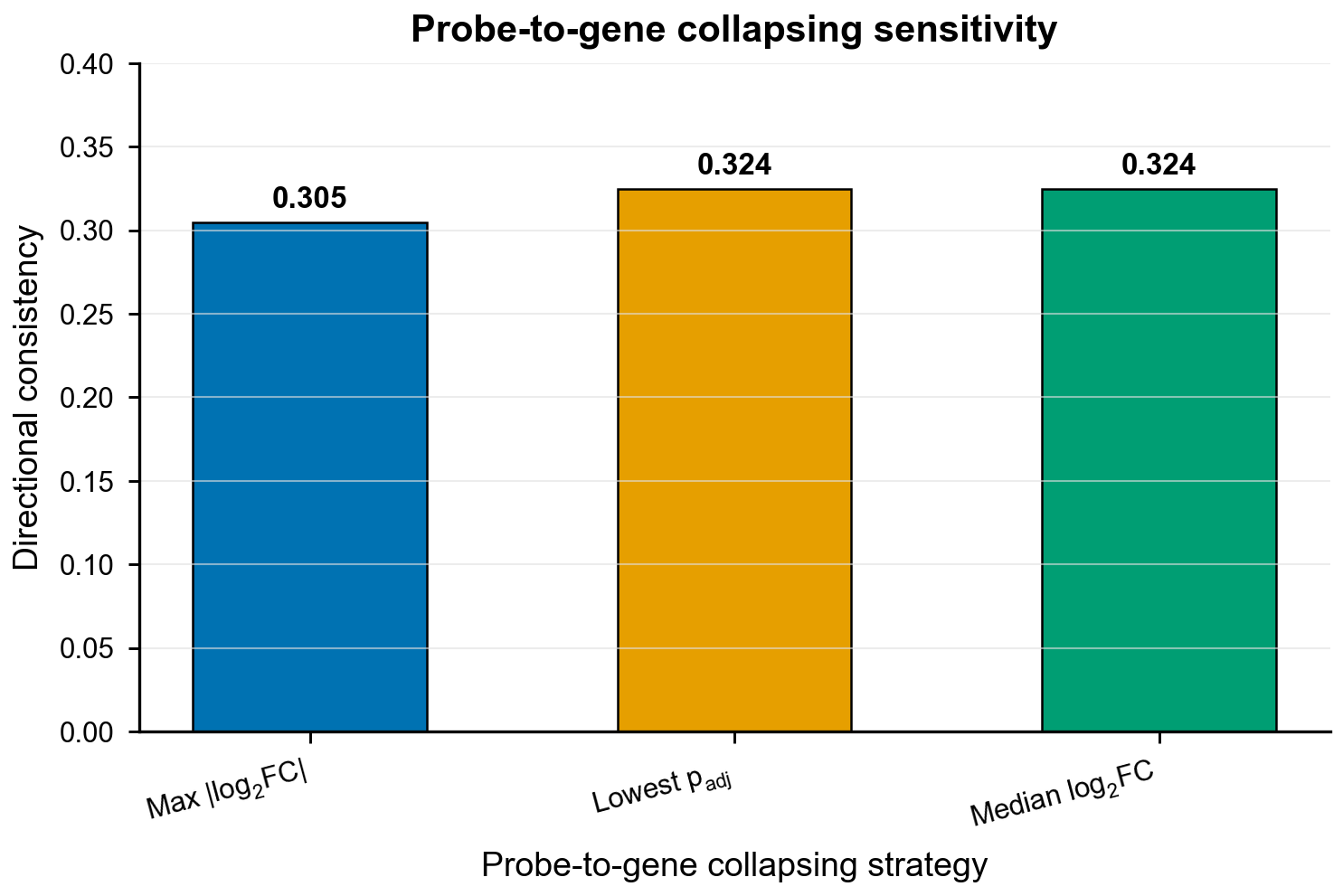

*Figure 9. Directional consistency is robust to probe-to-gene collapsing strategy. Observed directional consistency under three collapsing methods: maximum |log₂FC| (0.305), lowest adjusted p-value (0.324), and median log₂FC (0.324). Narrow range confirms robustness to annotation choice.*

| **Scenario** | **Gene-Identity Overlap** | **Directional Consistency** | **Recommendation** |
| --- | --- | --- | --- |
| Same platform, large sample size (n>200) | Informative | Informative | Both metrics appropriate |
| Cross-platform comparisons | Unreliable (sensitive to feature universe and probe definitions) | Robust (sign-based; insensitive to feature naming) | Prefer directional consistency |
| Small sample size (n < 50) | Unstable (few significant genes; high variance) | Cautious (wide confidence intervals) | Report both metrics with uncertainty estimates |
| Methylation (DMR- mapped to genes) | Structurally uninformative (overlap often deterministic due to mapping) | Partially informative (may be inflated without effect-size gating) | Use directional consistency with effect-size thresholding |
| Cross-modal comparisons (expression vs methylation) | Not applicable (different feature spaces) | Near-random (~50% agreement expected) | Neither metric informative |

**Table 7: Practical guidance for selecting reproducibility metrics under common cross-study comparison scenarios**

**FUNDINGS**

No external funding was received for this study.

**CONFLICT OF INTEREST DISCLOSURE**

The author(s) declared no potential conflicts of interest with respect to the research, authorship, and/or publication of this article.

**AUTHORS' CONTRIBUTIONS**

Inayat Chauhan designed the study, acquired raw data, and prepared the initial draft of the manuscript. Chiragh Dewan performed the bioinformatics analysis. Kanika Sharma provided valuable inputs for compiling the manuscript, editing, and critical review of the study. Sangeeta Sharma and Rupinderjeet Kaur supervised the study.

**DATA, SCRIPT, CODE, AND SUPPLEMENTARY INFORMATION AVAILABILITY**

All analysis scripts were executed in R (version 4.5.1) and Python (version 3.13.7). Reproducible workflows and configuration files are [PLACEHOLDER: Scripts to be deposited on Zenodo prior to submission. Replace this text with: Scripts available at https://doi.org/10.5281/zenodo.XXXXXXX (Zenodo DOI to be added before submission)]. All analyses were performed with fixed random seeds (seed = 42) to ensure reproducibility.

**SUPPLEMENTARY MATERIALS**

Supplementary figures and tables provide detailed quality control diagnostics, calibration assessments, direction-stratified overlap statistics, and robustness analyses that support the main methodological conclusions of the study. These materials are intended to document analytical validity and parameter sensitivity rather than biological interpretation.

**Supplementary Table S1: Covariate Availability Across Datasets**

| **Dataset** | **Age** | **Sex** | **Batch** | **Cell Fractions** |
| --- | --- | --- | --- | --- |
| GSE99039 | — | ✓ | ✓ |  |
| GSE6613-CEL | — | ✓ | — | — |
| GSE165083-GPL11154 | — | ✓ | — | — |
| GSE145361 | — | ✓ | — | — |
| GSE72774 | ✓ | ✓ | — | — |
| GSE111629 | ✓ | ✓ | — | — |
| GSE165083-methylation | — | ✓ | — | ✓ |

*Supplementary Table S1: Availability of demographic and technical covariates. ✓ indicates valid numeric covariate values present for the majority of samples. — indicates absent or missing values. Cell-type fraction estimates (EpiDISH-derived) were available for GSE165083-methylation only but were not included as covariates in any differential analysis.*

**Supplementary Table S2: Genomic Inflation Factors and Calibration Diagnostics**

| **Dataset** | **Modality** | **λGC** | **N P-values** | **Interpretation** |
| --- | --- | --- | --- | --- |
| GSE165083 – GPL11154 | Expression | 4.15 | 21,062 | Substantial Inflation |
| GSE6613-CEL | Expression | 5.03 | 8,064 | Substantial Inflation |
| GSE99039 | Expression | 3.95 | 9,779 | Substantial Inflation |
| GSE145361 | Methylation | 123.1 | 20,564 | Substantial Inflation |
| GSE72774 | Methylation | 114.7 | 12,404 | Substantial Inflation |

*Supplementary Table S2: For methylation datasets, region-level λGC values are shown; CpG-level λGC values (1.04 and 0.99 respectively) are reported in the main text. GSE111629 and GSE165083-methylation produced insufficient region-level p-values for stable λGC estimation and are omitted from region-level diagnostics.*

**Supplementary Table S3: Direction-Stratified Overlap Counts**

| **Comparison** | **Direction Type** | **N Overlap** |
| --- | --- | --- |
| GSE6613-CEL vs GSE99039 | Up – Up | 30 |
| GSE6613-CEL vs GSE99039 | Down – Down | 45 |
| GSE6613-CEL vs GSE99039 | Discordant | 7 |

*Supplementary Table S3: Counts of overlapping features stratified by direction of effect for pairwise dataset comparisons. These counts underlie the directional consistency metrics reported in the main text.*

**Supplementary Table S4: Pairwise Null-Model Context for Directional Consistency**

| **Comparison** | **N Testable Features** | **Observed Agreement (%)** | **Expected Agreement (%)** | **Excess Agreement (%)** | **Interpretation** |
| --- | --- | --- | --- | --- | --- |
| GSE6613 vs GSE99039 | 82 | 91.5 | 74.8 | 16.7 | Modest directional consistency |
| n ≥ 3 DEG (combined) | 136 | 98.3 | 74.8 | 23.4 | Strong excess agreement |
| Null Expectation | — | — | 74.8 | — | Random-direction baseline |

**Supplementary Table S4: Pairwise Null-Model Context for Directional Consistency**

*Supplementary Table S4: Permutation-based null-model statistics. Expected agreement was derived from permutation procedures using fixed random seeds and 10,000 iterations.*

Supplementary Figures S1–S12: Quality control diagnostics, calibration QQ plots, gene-mapping expansion plots, distributional properties of directional consistency metrics, direction-stratified overlap heatmaps, and rank concordance heatmaps (to be provided as separate figure files).
