## Supplementary material for "Directional Gene-Level Concordance and Methodological Constraints in Blood Transcriptomic and DNA Methylation Studies of Parkinson’s Disease": Code files: Supplementary Figures PCI.docx

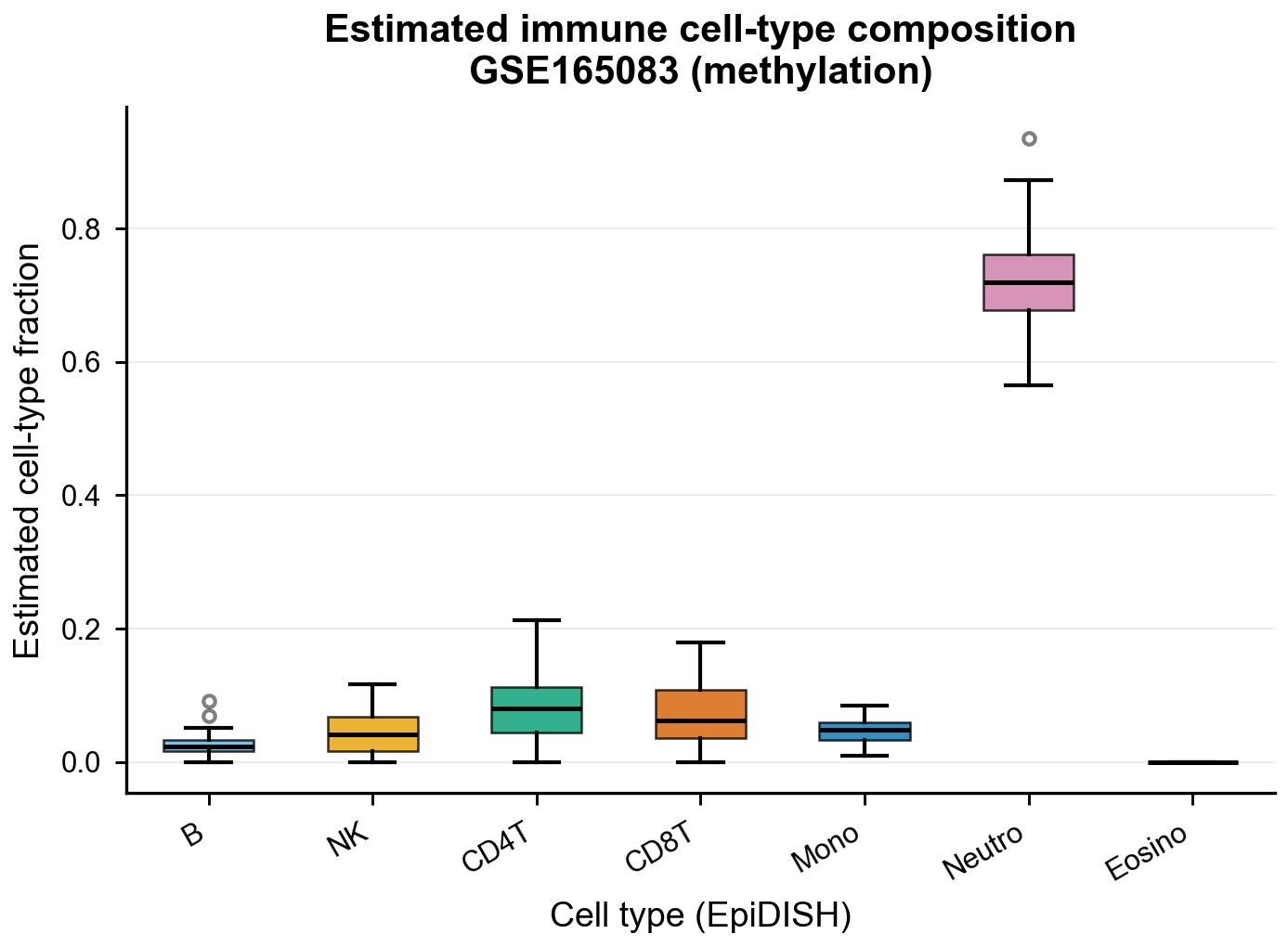
***Supplementary Figure S1. Estimated immune cell-type composition for GSE165083 (methylation):*** *Boxplots show EpiDISH-estimated proportions of major immune cell populations across samples. Cell-type fractions were used solely for quality control and descriptive assessment; no covariate adjustment using these estimates was performed in the primary differential analyses.*

.

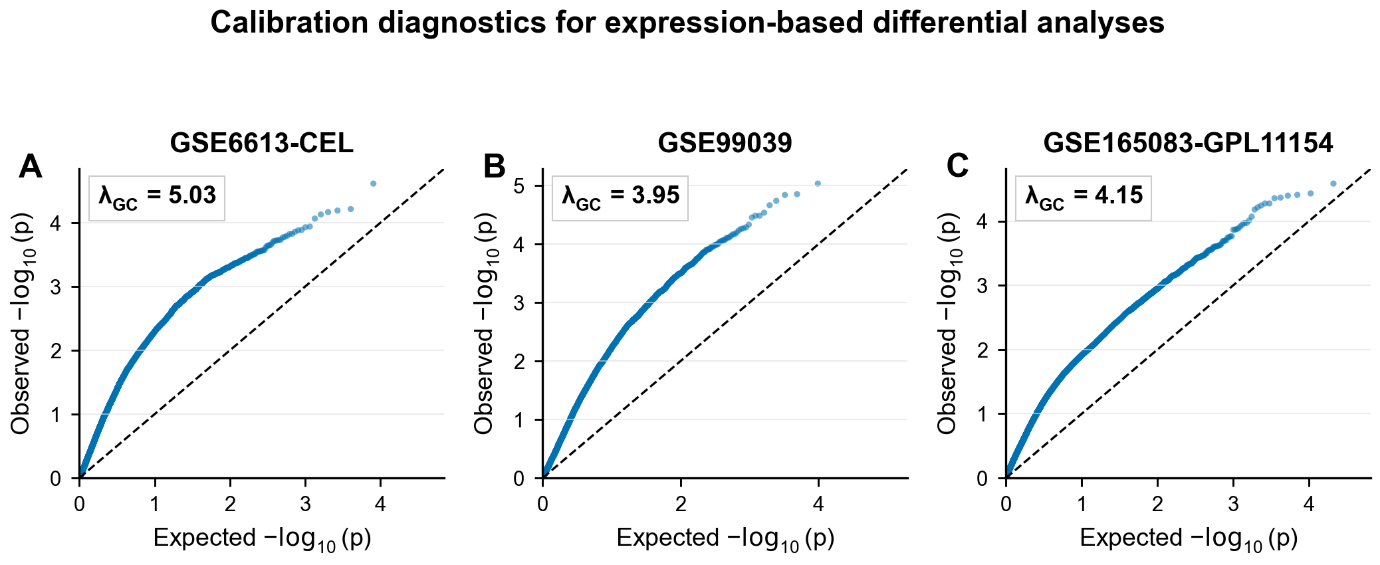

***Supplementary Figure S2. Calibration diagnostics for expression-based differential Analyses:*** *QQ plots of gene-level p-values against the uniform null for all three expression datasets. Genomic inflation factors (λ_GC) are annotated on each panel. Substantial deviation from y = x motivates emphasis on relative and directional metrics rather than absolute significance counts.*

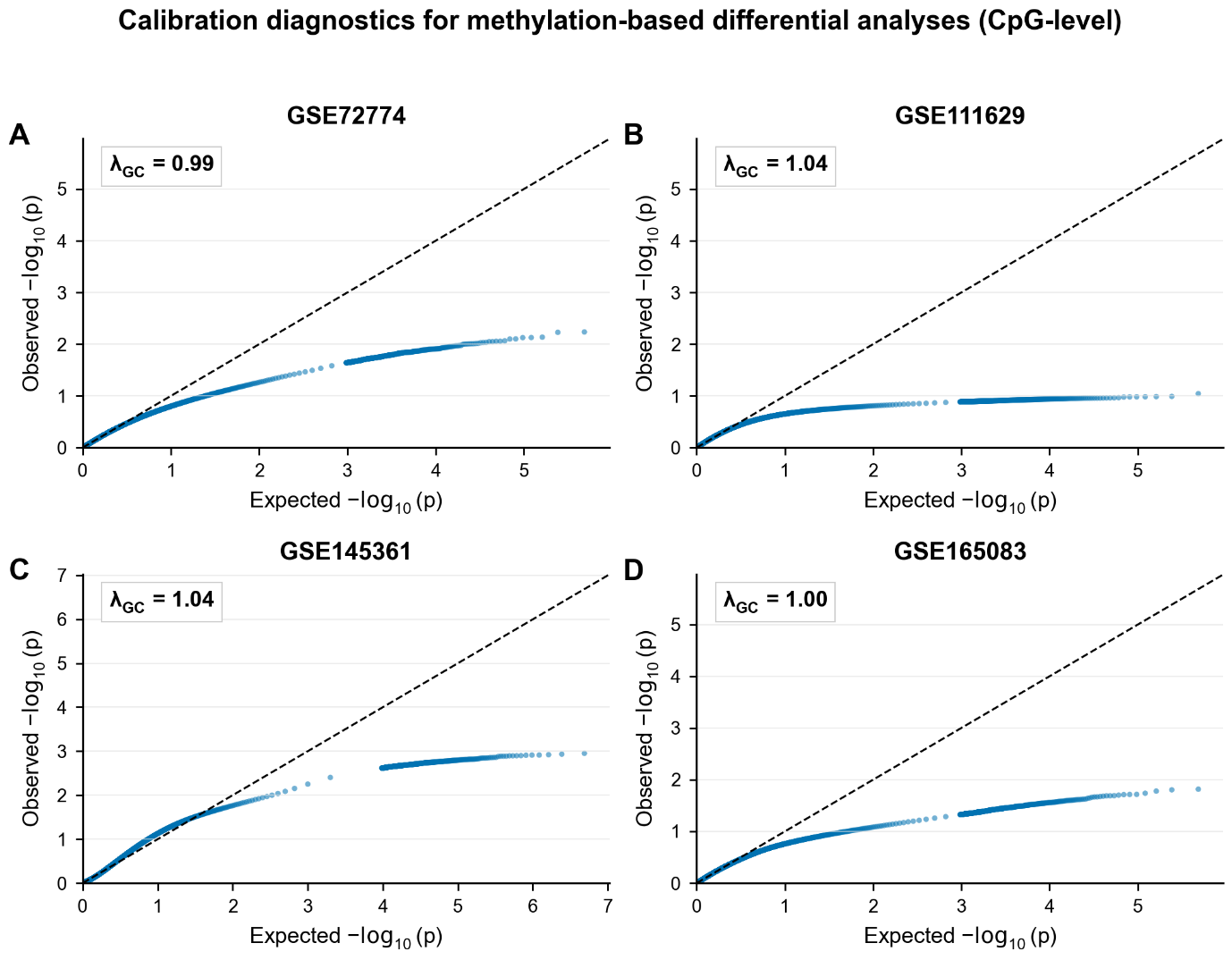

***Supplementary Figure S3. Calibration diagnostics for methylation-based differential analyses:*** *CpG-level QQ plots across four methylation datasets. CpG-level λ_GC values are close to 1.00 (0.99–1.04), confirming per-probe calibration; the substantially elevated region-level λ_GC values (> 100) reflect aggregation of correlated CpG signals within DMRs, not probe-level miscalibration.*

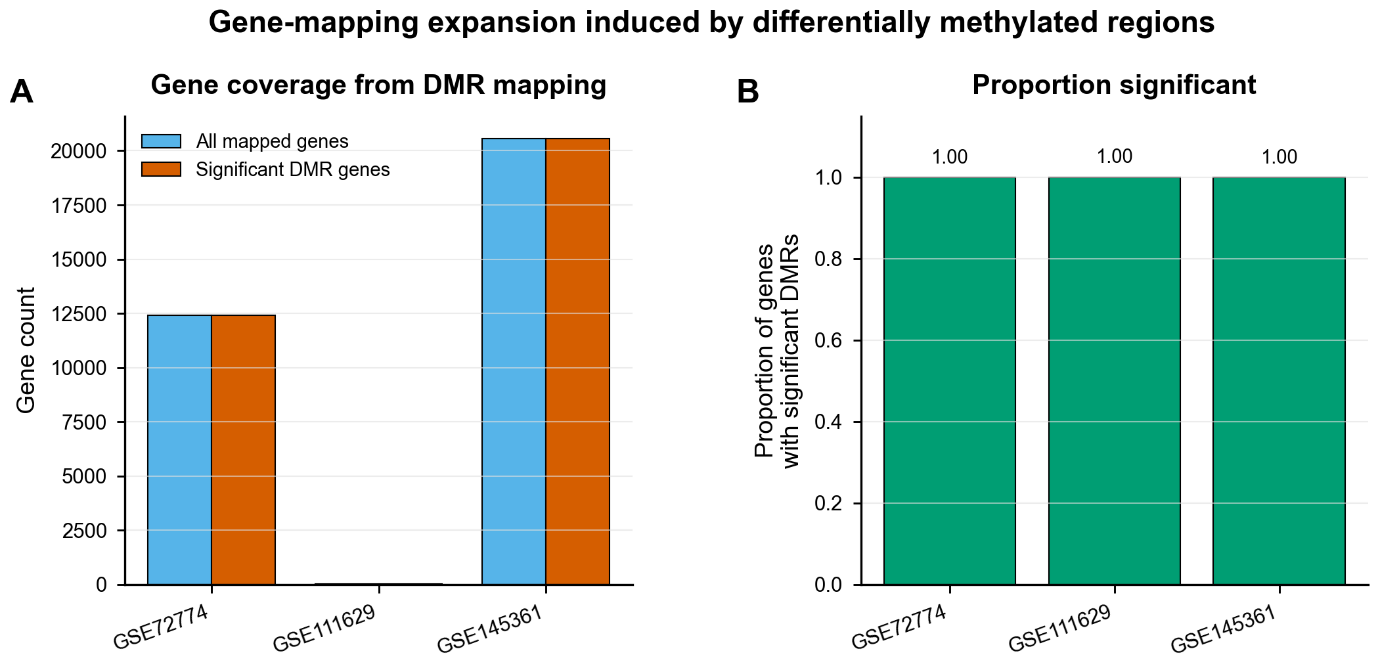

***Supplementary Figure S4. Gene-mapping expansion induced by differentially methylated regions:*** *(A) Total number of genes covered by any DMR mapping versus genes covered by significant DMRs. (B) Proportion of mapped genes that harbor a significant DMR. Near-complete coverage demonstrates that overlap and enrichment metrics for methylation data are driven by mapping structure rather than independent biological replication.*

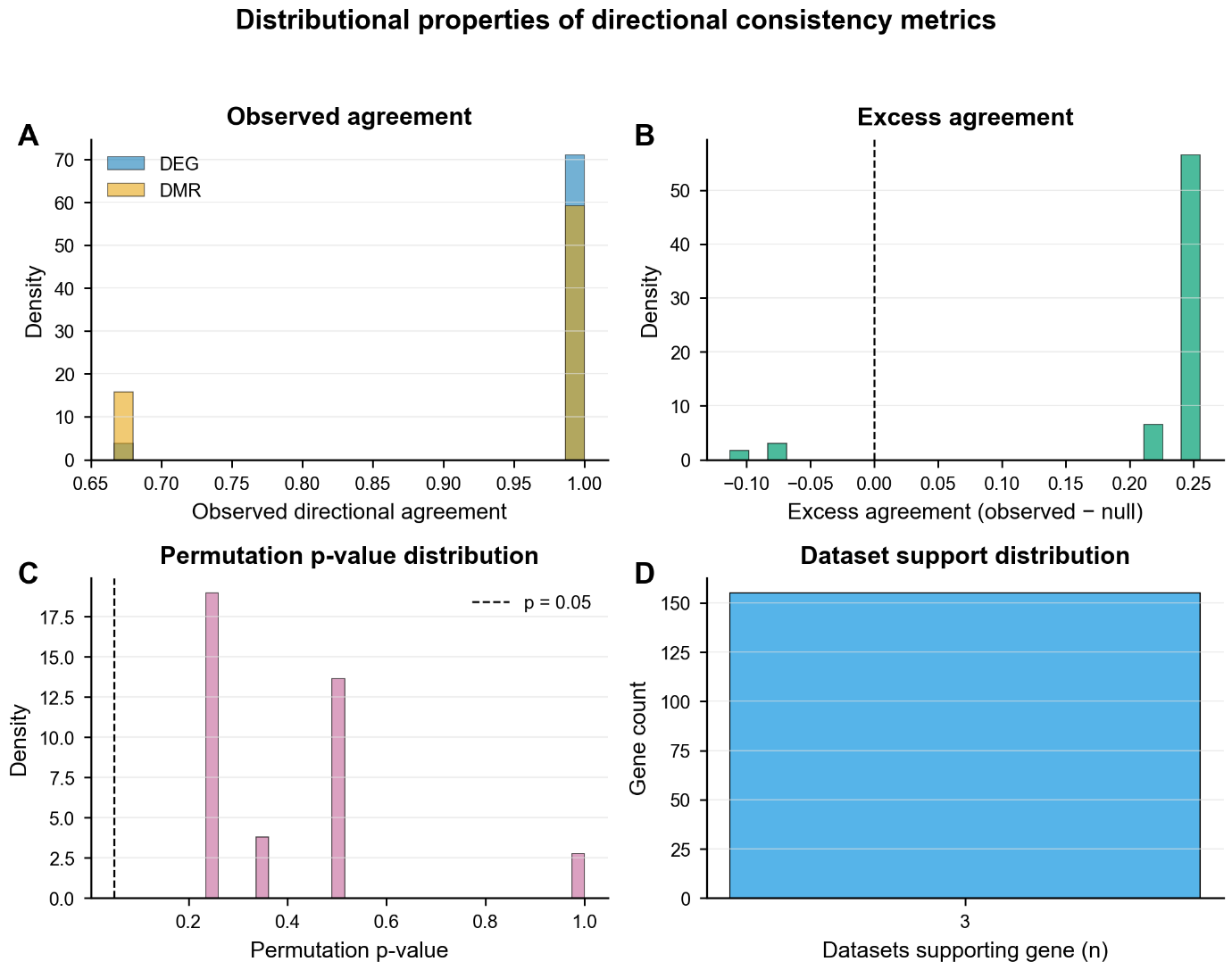

***Supplementary Figure S5. Distributional properties of directional consistency metrics:*** *(A) Observed agreement density stratified by modality. (B) Excess agreement (observed − null). (C) Permutation p-value distribution with p = 0.05 reference line. (D) Dataset-support histogram (genes by number of contributing datasets).*

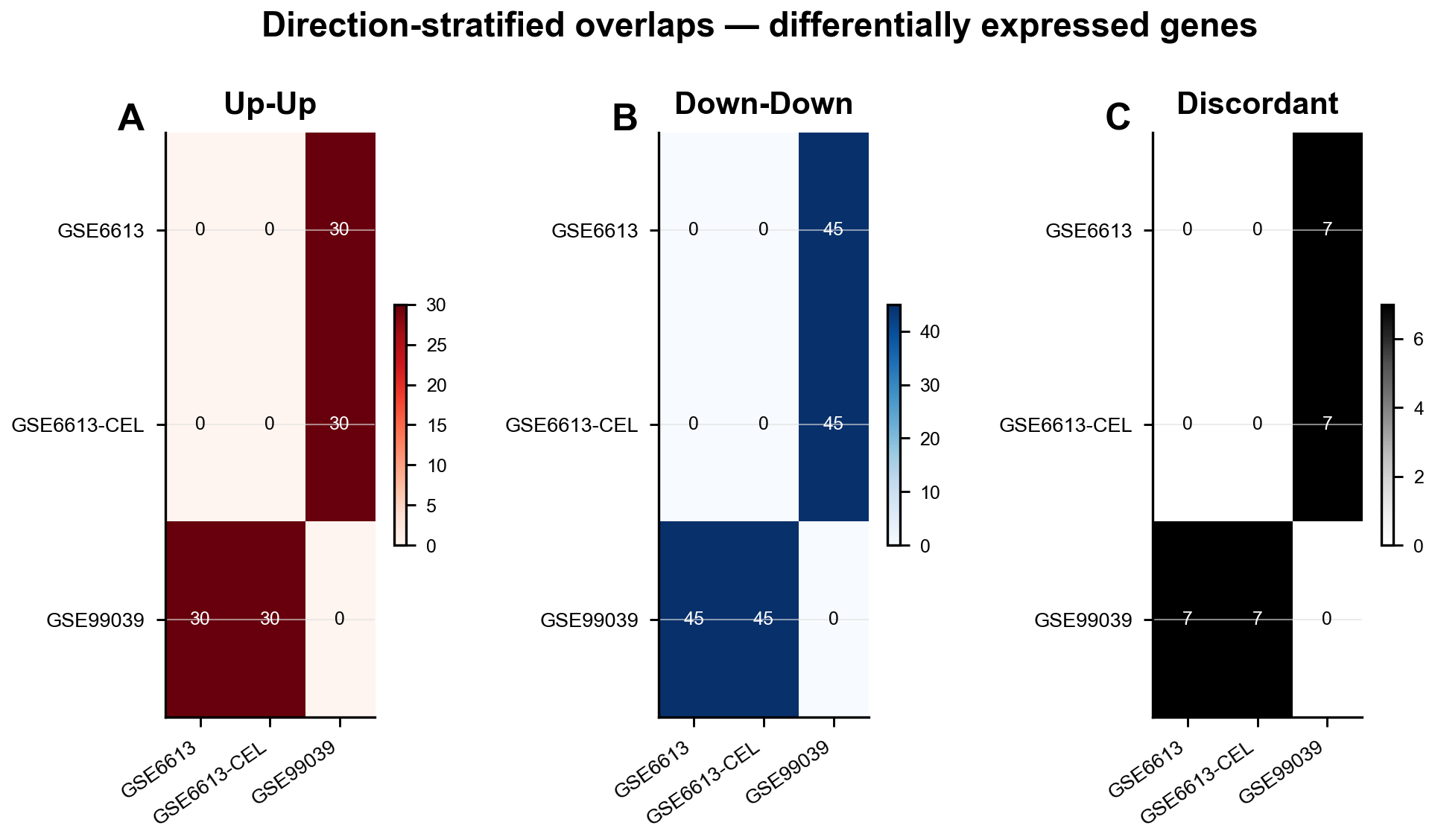

***Supplementary Figure S6. Direction-stratified overlaps for differentially expressed genes:*** *Heatmaps of raw pairwise overlap counts stratified by effect direction: (A) up-up, (B) down-down, (C) discordant. Values are not normalized by feature universe size. Cell values are annotated****.***

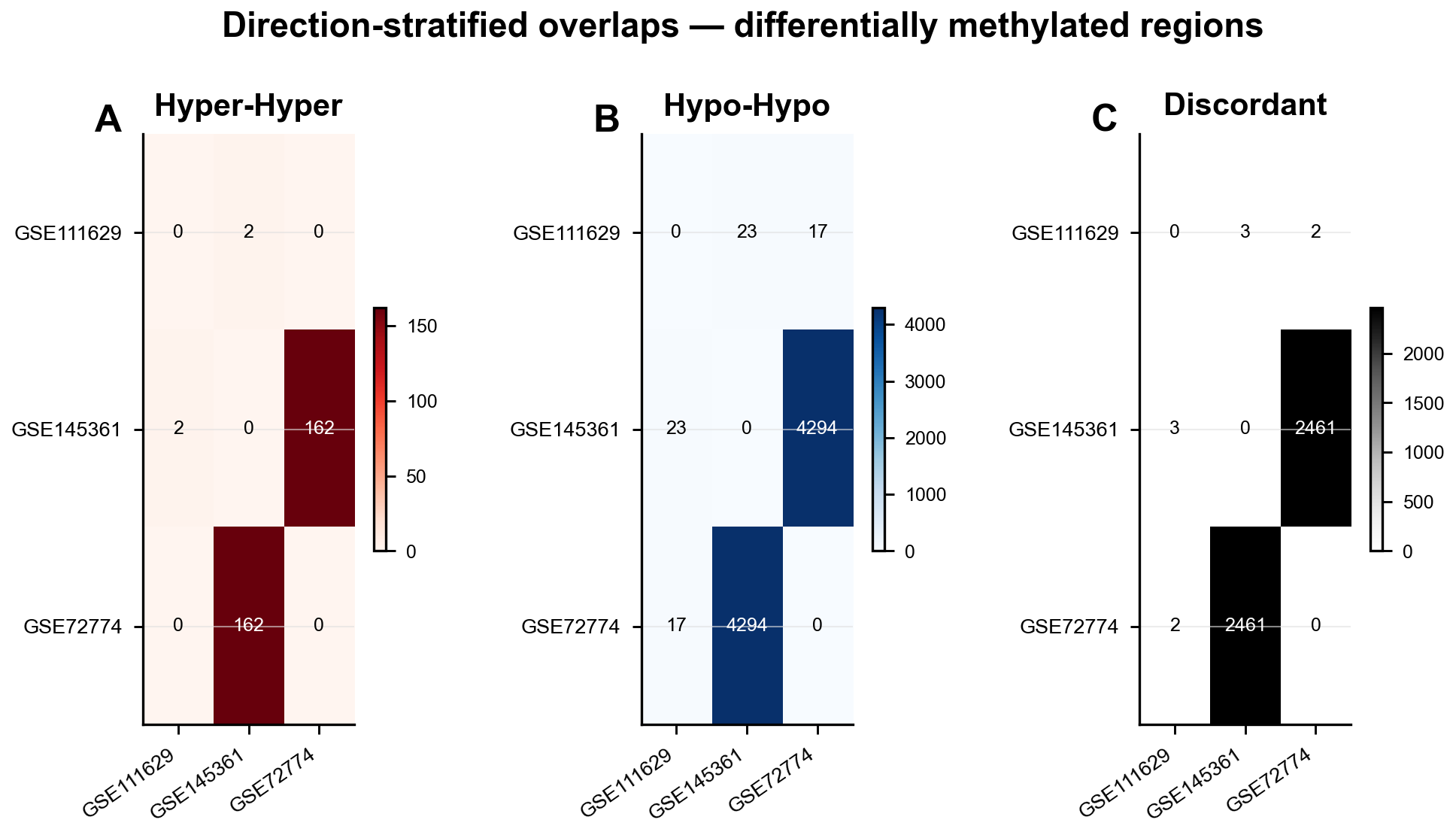

***Supplementary Figure S7. Direction-stratified overlaps for differentially methylated regions:*** *Heatmaps of raw overlap counts stratified by methylation direction: (A) hyper-hyper, (B) hypo-hypo, (C) discordant. High absolute values reflect deterministic mapping and feature-space properties rather than independent biological replication.*

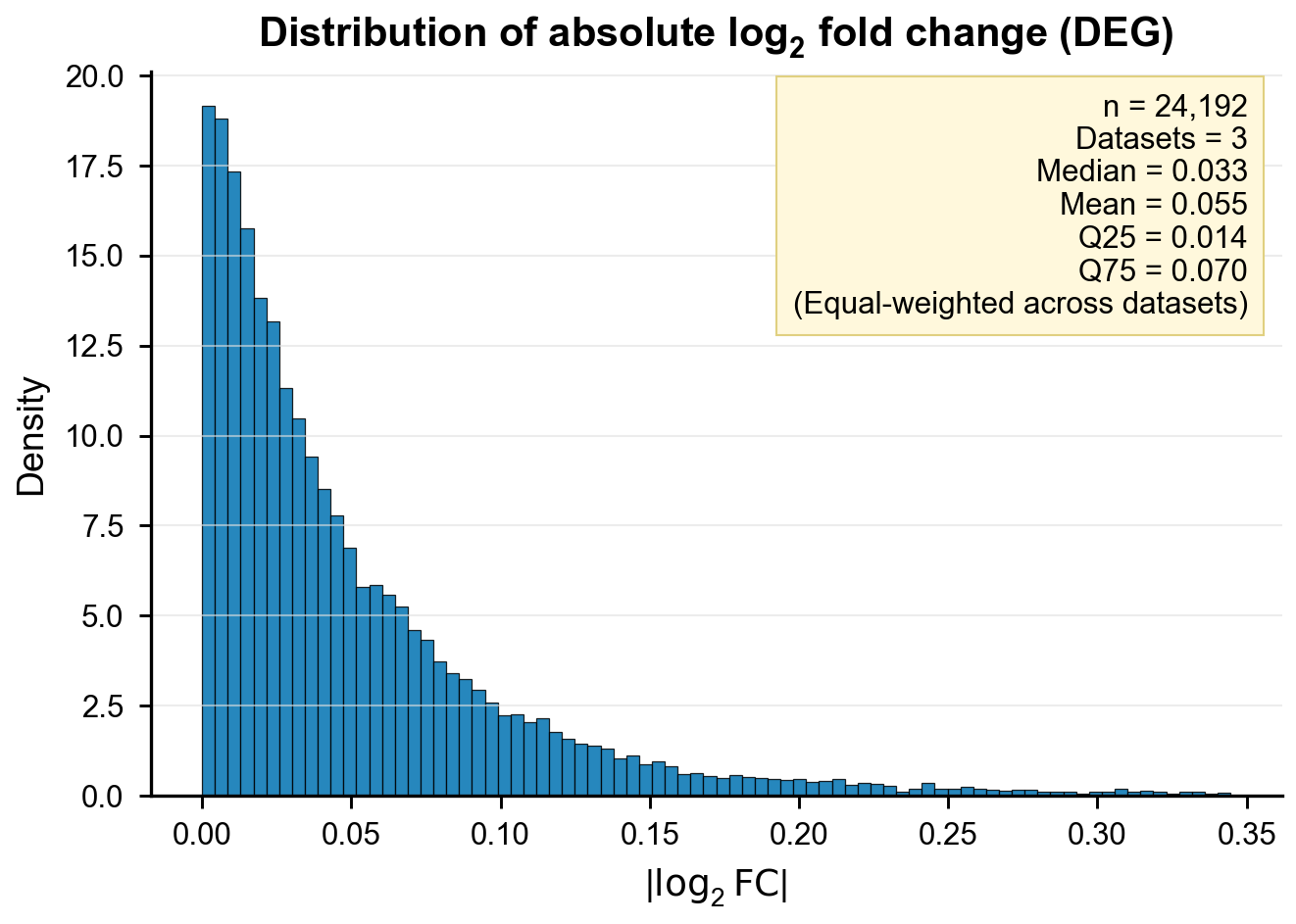

***Supplementary Figure S8. Distribution of absolute log$_2$ fold-change values for differentially expressed genes****: Histogram of |log$_2$FC| across three expression datasets (equal-weighted by sorted gene identifier). Summary statistics (median, mean, IQR) shown in the inset. Descriptive only; no statistical comparison across modalities was performed.*

*.*

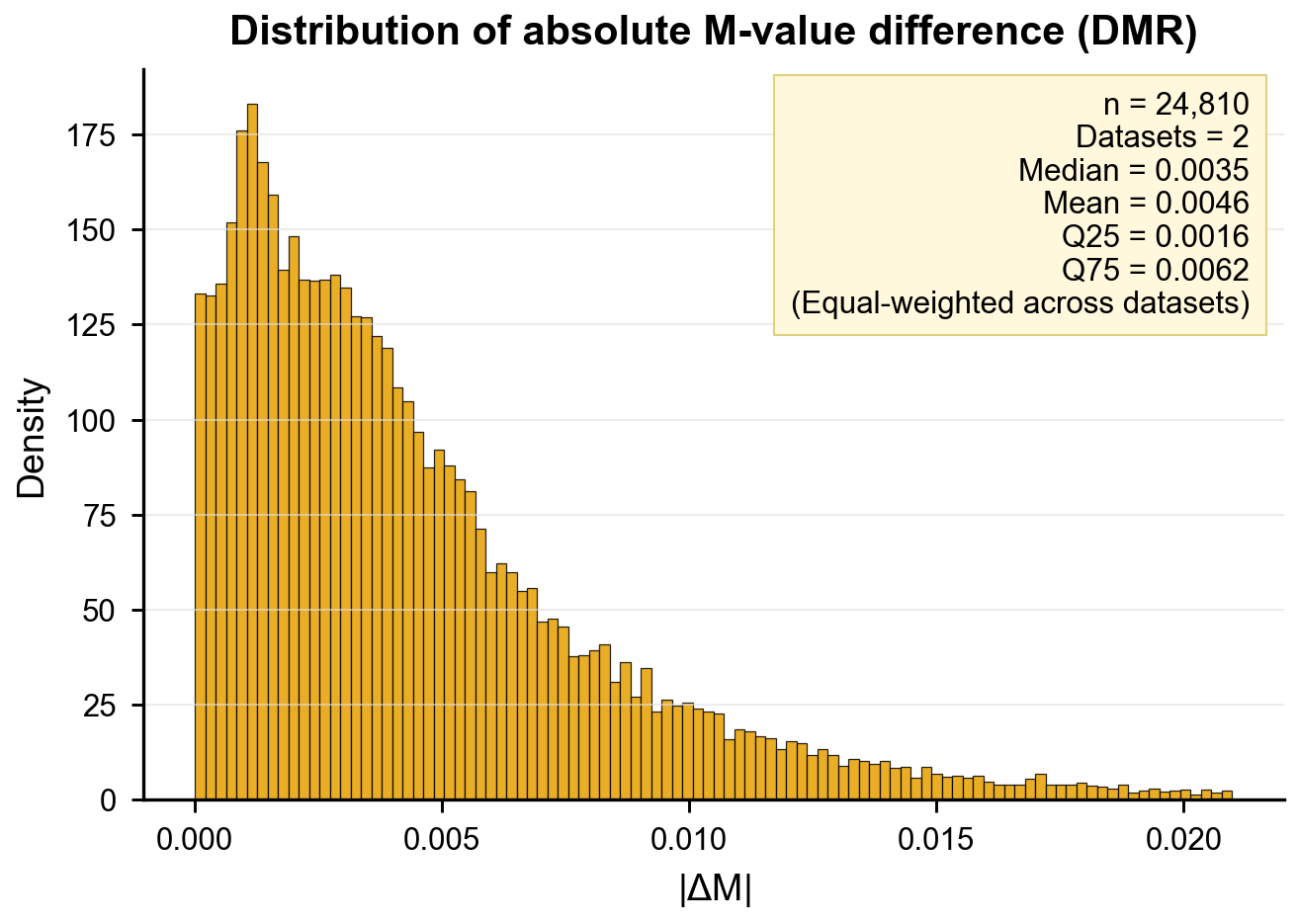

***Supplementary Figure S9. Distribution of absolute M-value differences for differentially methylated regions:*** *Histogram of |ΔM| across DMRs in two datasets (GSE72774, GSE145361; GSE111629 excluded as below the equal-weighting minimum). Summary statistics shown in the inset. Descriptive only.*

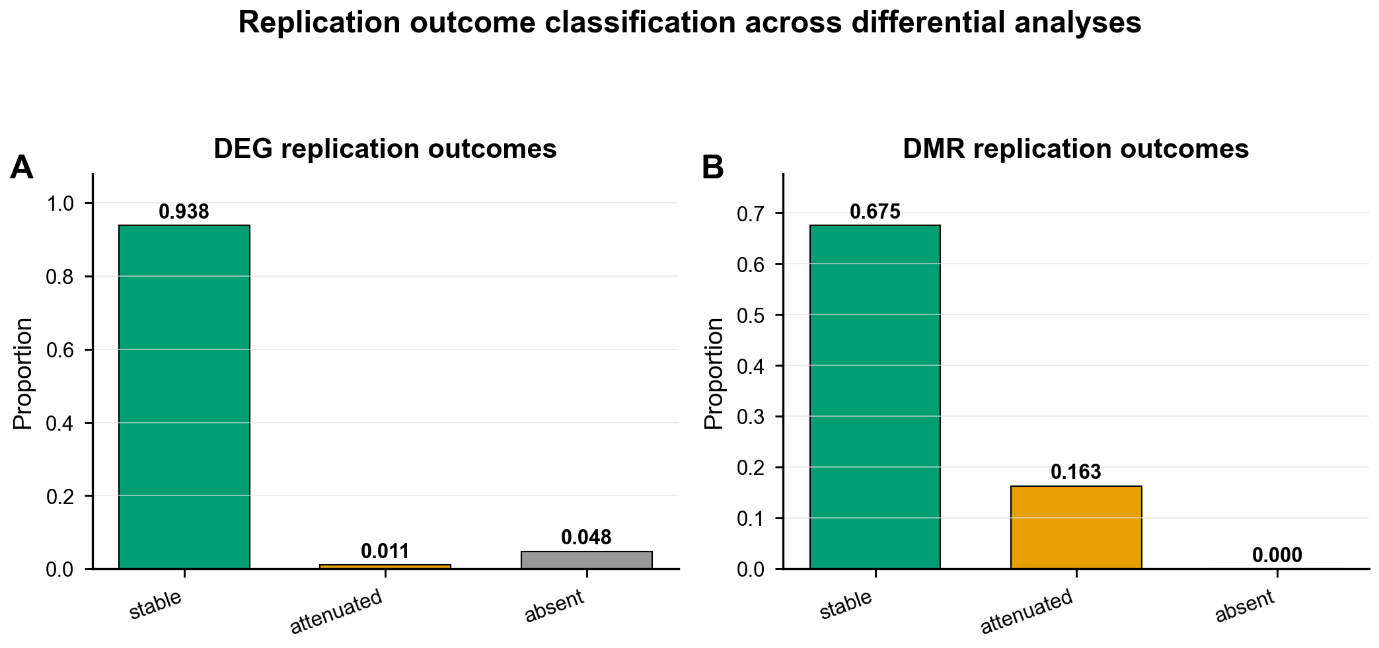

***Supplementary Figure S10. Replication outcome classification for differential expression and differential methylation:*** *Features categorized as stable, attenuated, direction-discordant, or absent across datasets. Proportions shown separately for (A) DEG and (B) DMR analyses.*

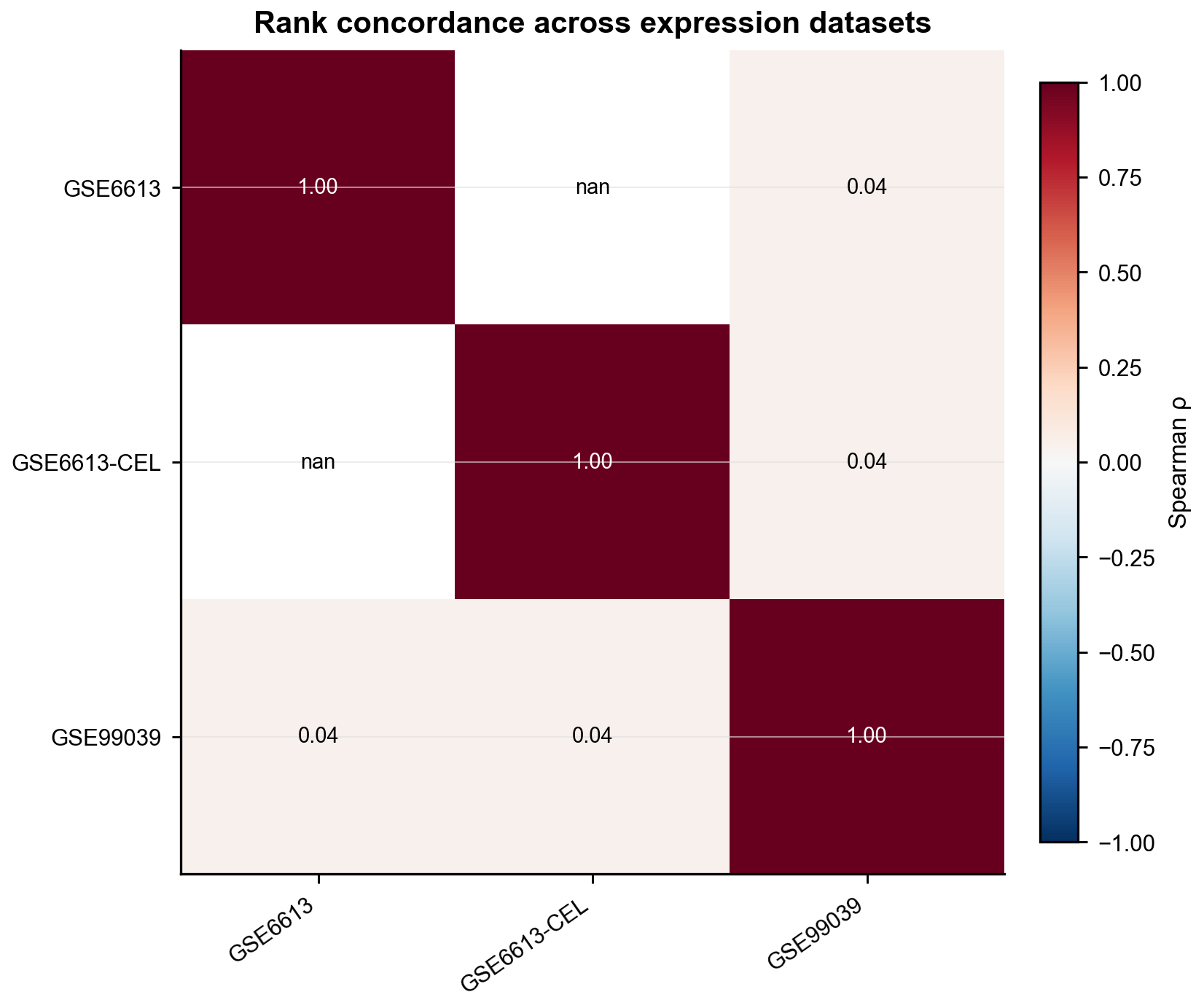

***Supplementary Figure S11. Rank concordance heatmap across expression datasets:*** *Pairwise Spearman rank correlations between gene-level differential expression statistics. Diagonal = self-comparisons. Cell values annotated.*

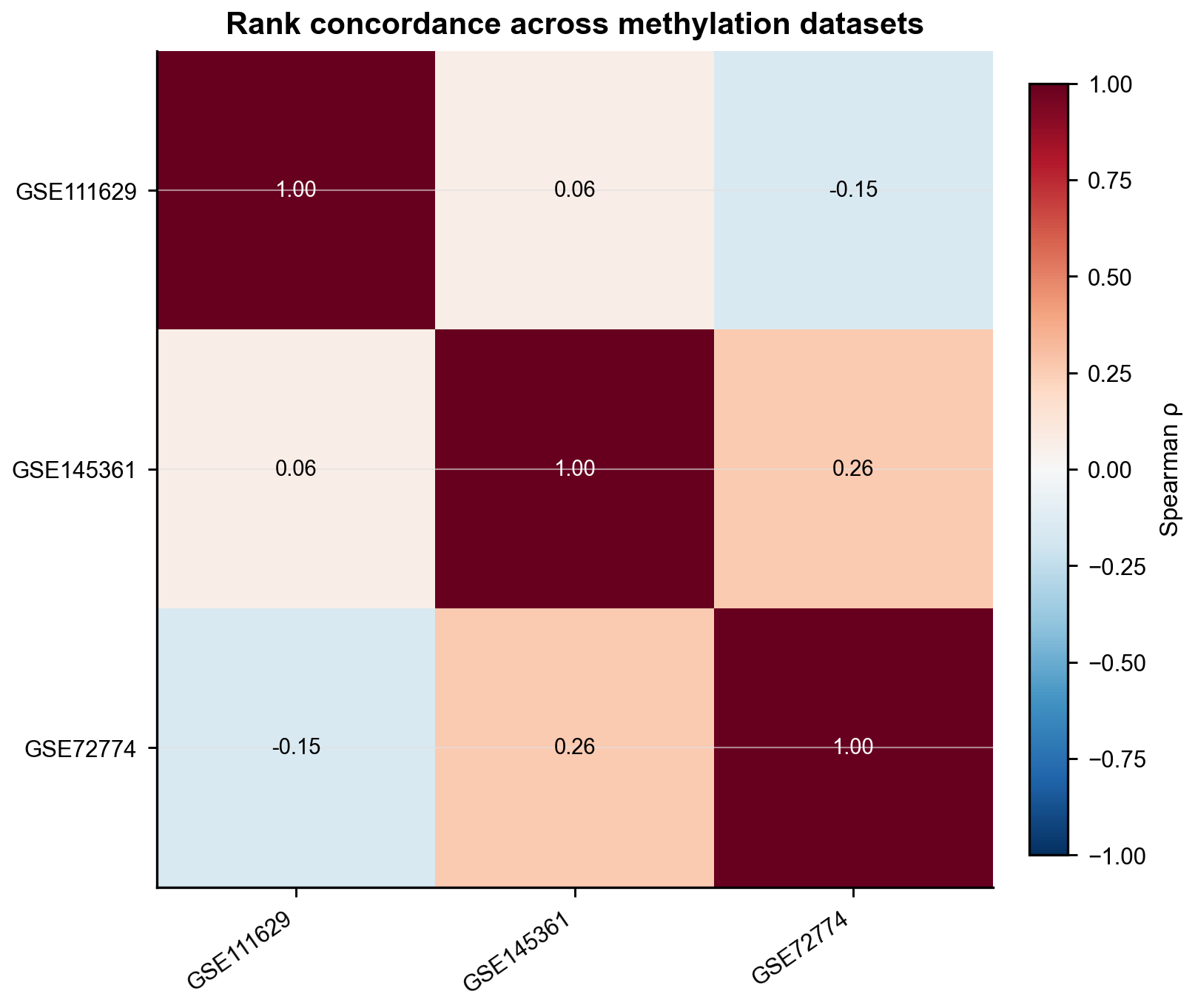

***Supplementary Figure S12. Rank concordance heatmap across methylation datasets****: Pairwise Spearman rank correlations between DMR-level statistics. Off-diagonal values vary with annotation structure and dataset-specific variability.*
