## Supplementary figures and images for "Directional Gene-Level Concordance and Methodological Constraints in Blood Transcriptomic and DNA Methylation Studies of Parkinson’s Disease"

### d7c0e1d3-23c4-402a-a867-f3c095c64fc1-0.jpg

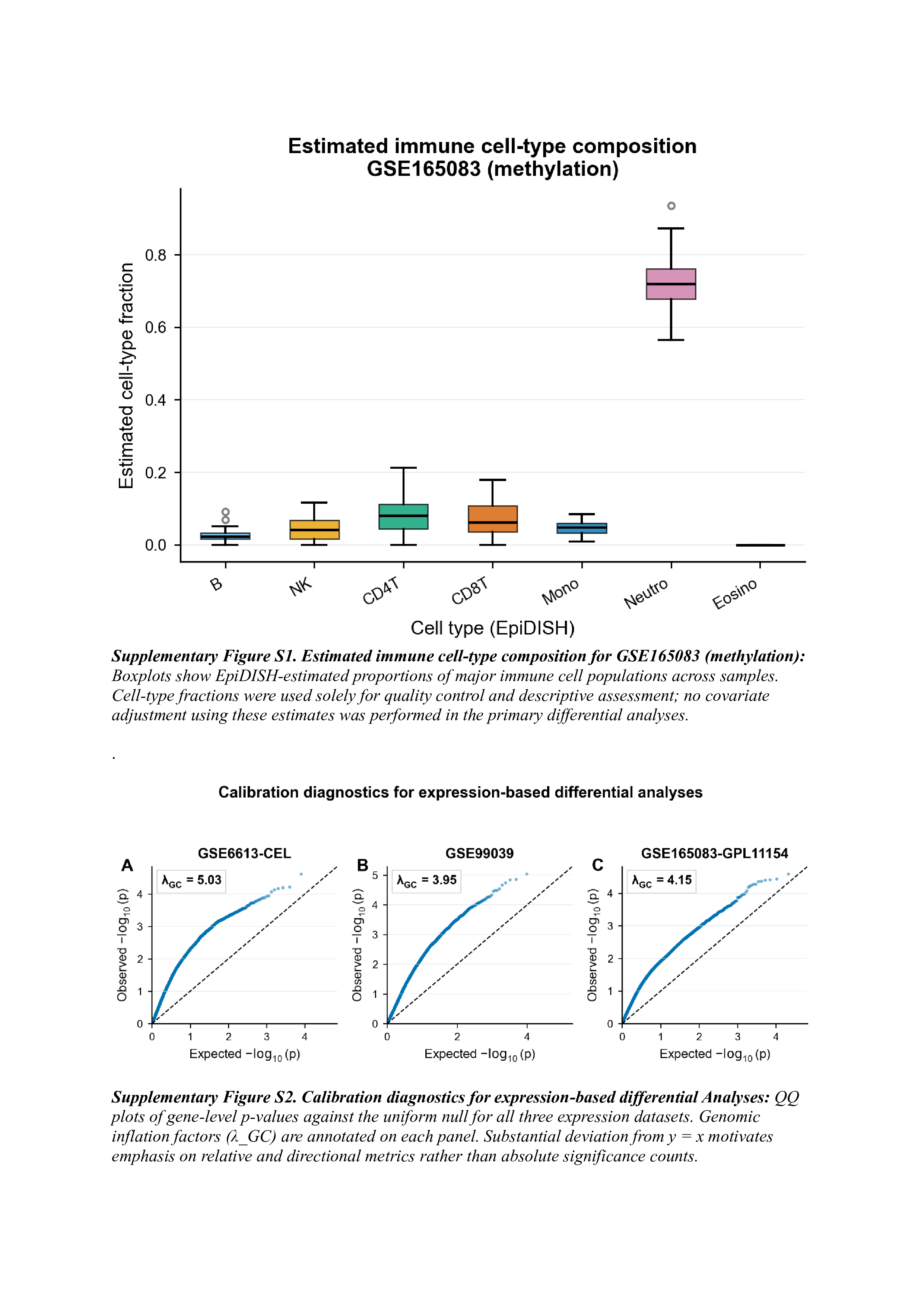

### d7c0e1d3-23c4-402a-a867-f3c095c64fc1-1.jpg

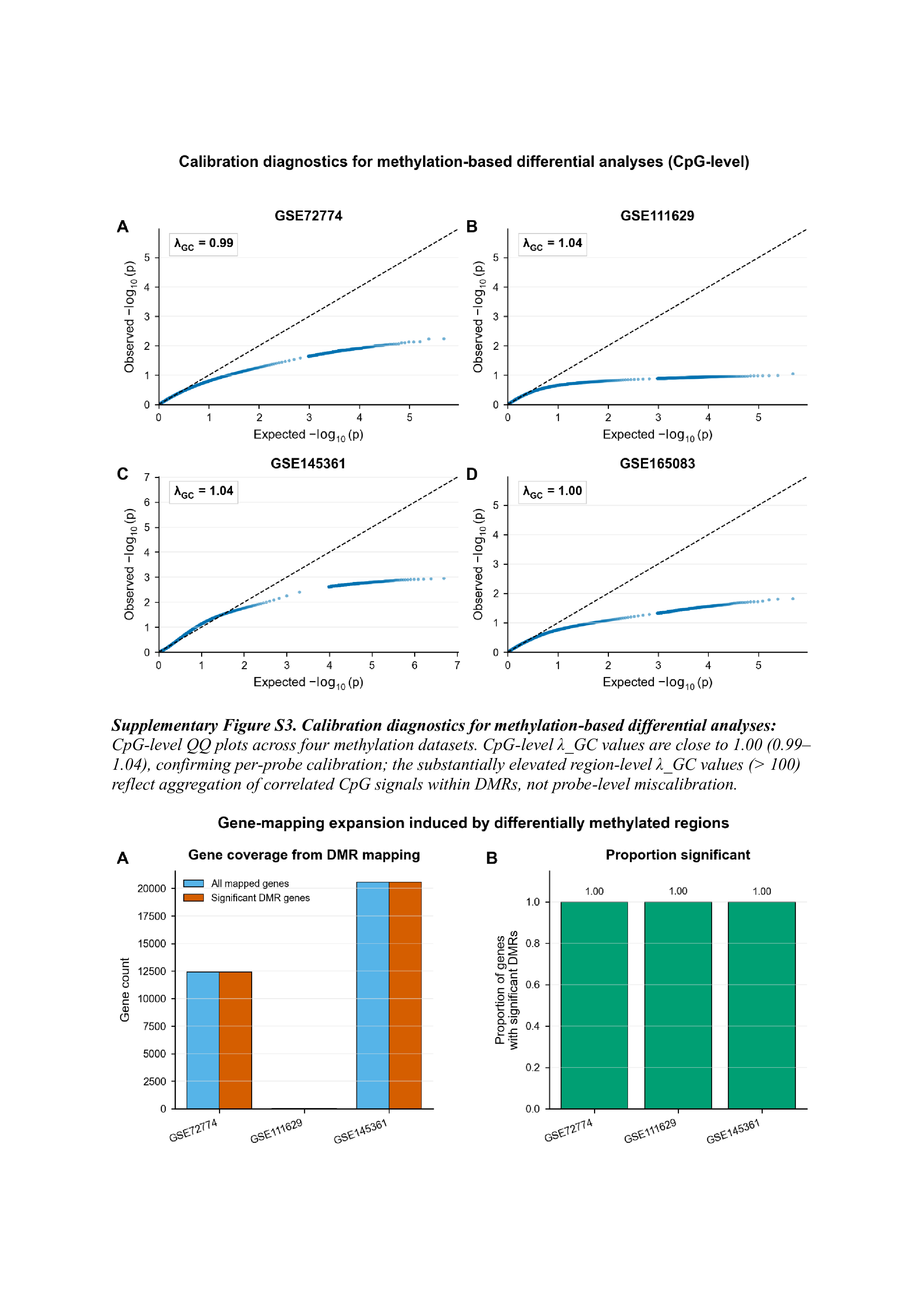

### d7c0e1d3-23c4-402a-a867-f3c095c64fc1-2.jpg

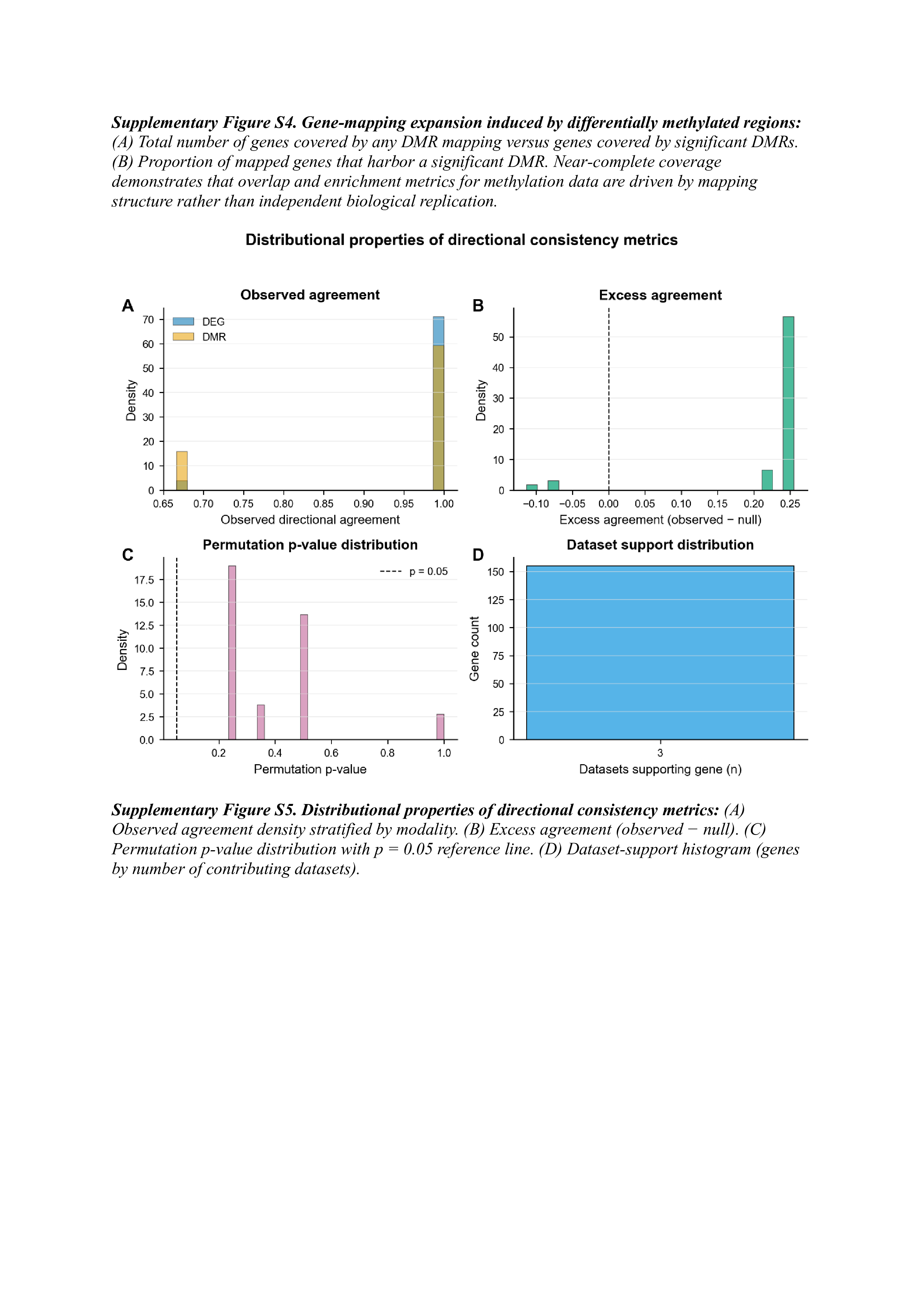

### d7c0e1d3-23c4-402a-a867-f3c095c64fc1-3.jpg

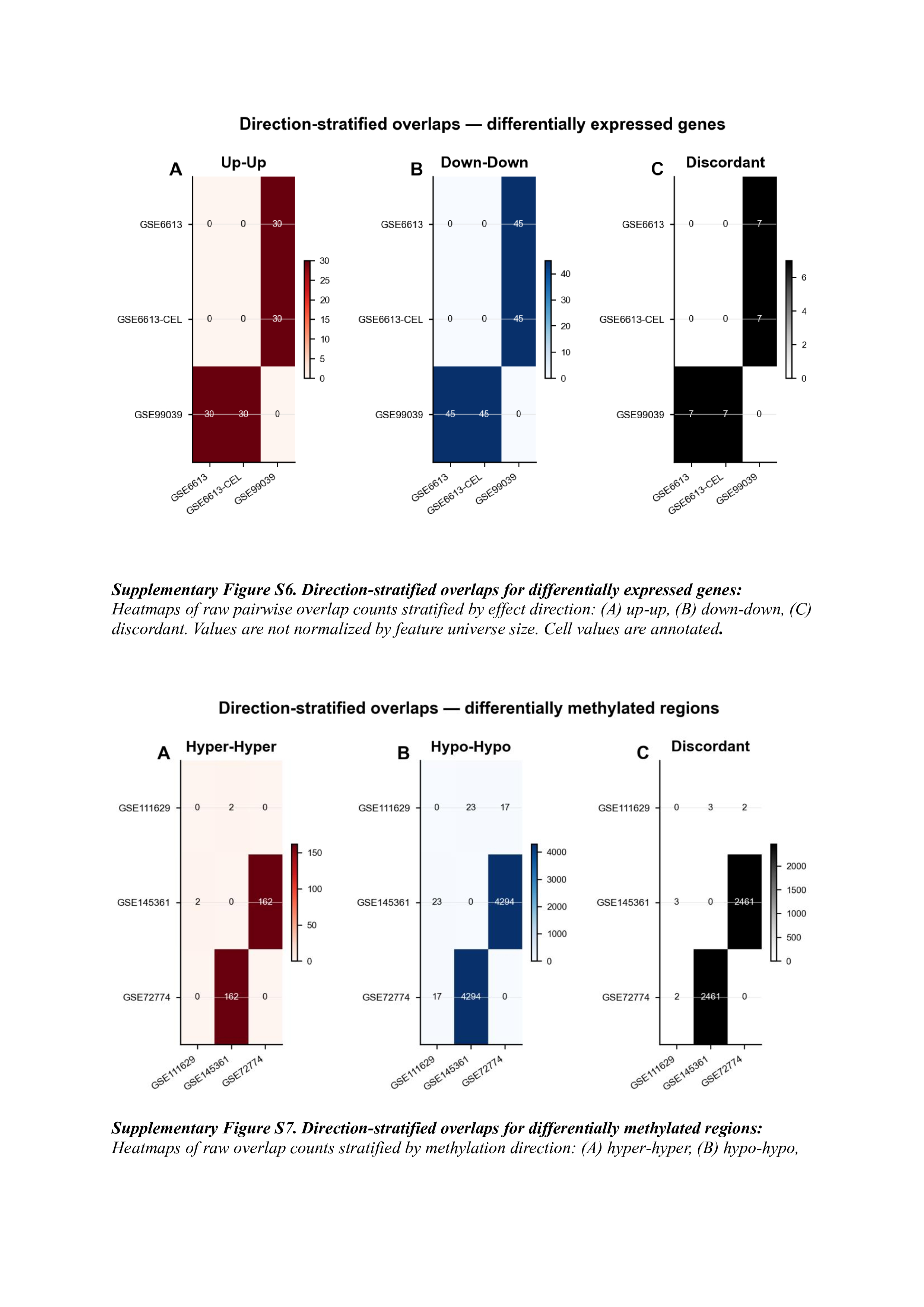

### d7c0e1d3-23c4-402a-a867-f3c095c64fc1-4.jpg

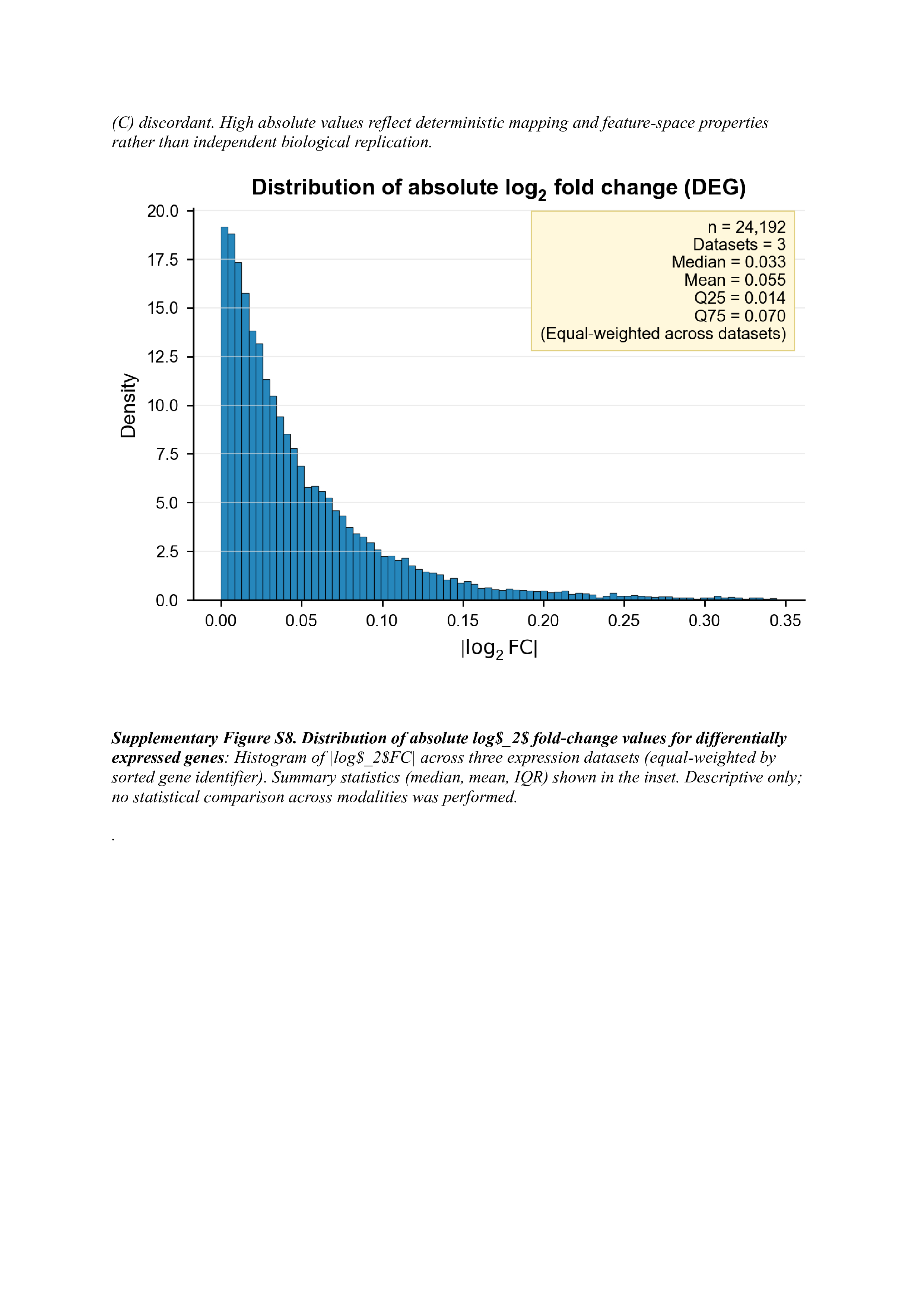

### d7c0e1d3-23c4-402a-a867-f3c095c64fc1-5.jpg

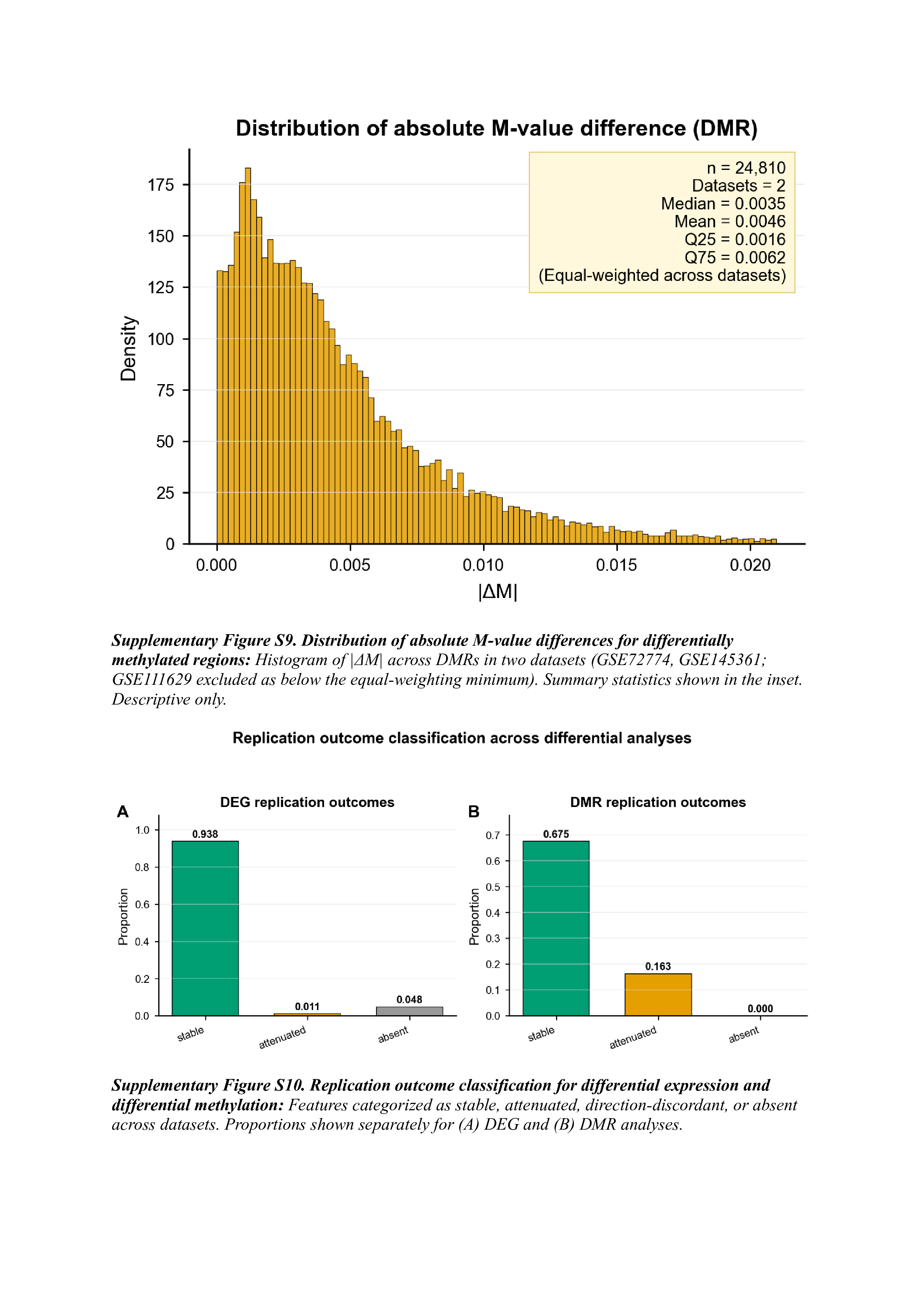

### d7c0e1d3-23c4-402a-a867-f3c095c64fc1-6.jpg

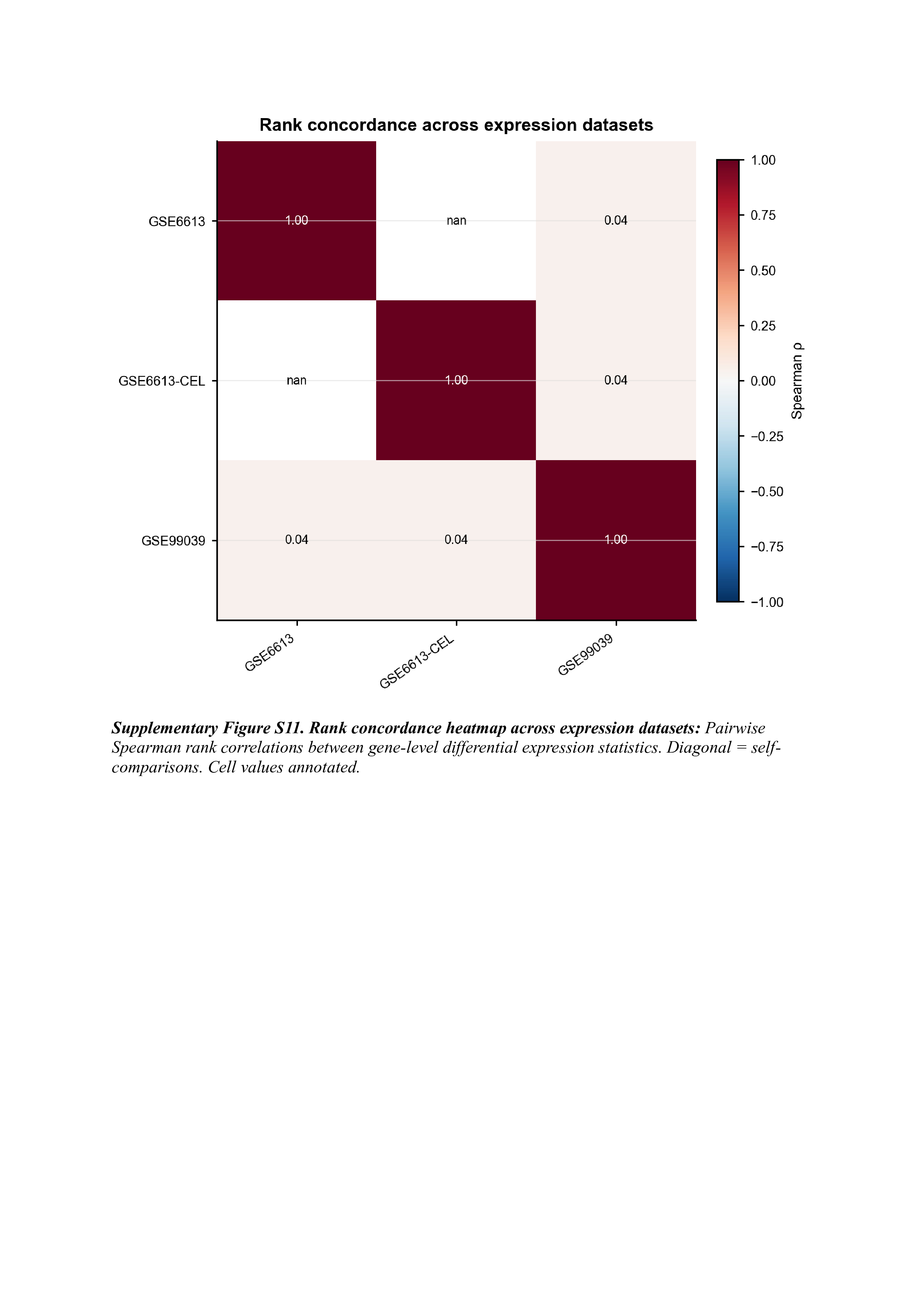

### d7c0e1d3-23c4-402a-a867-f3c095c64fc1-7.jpg

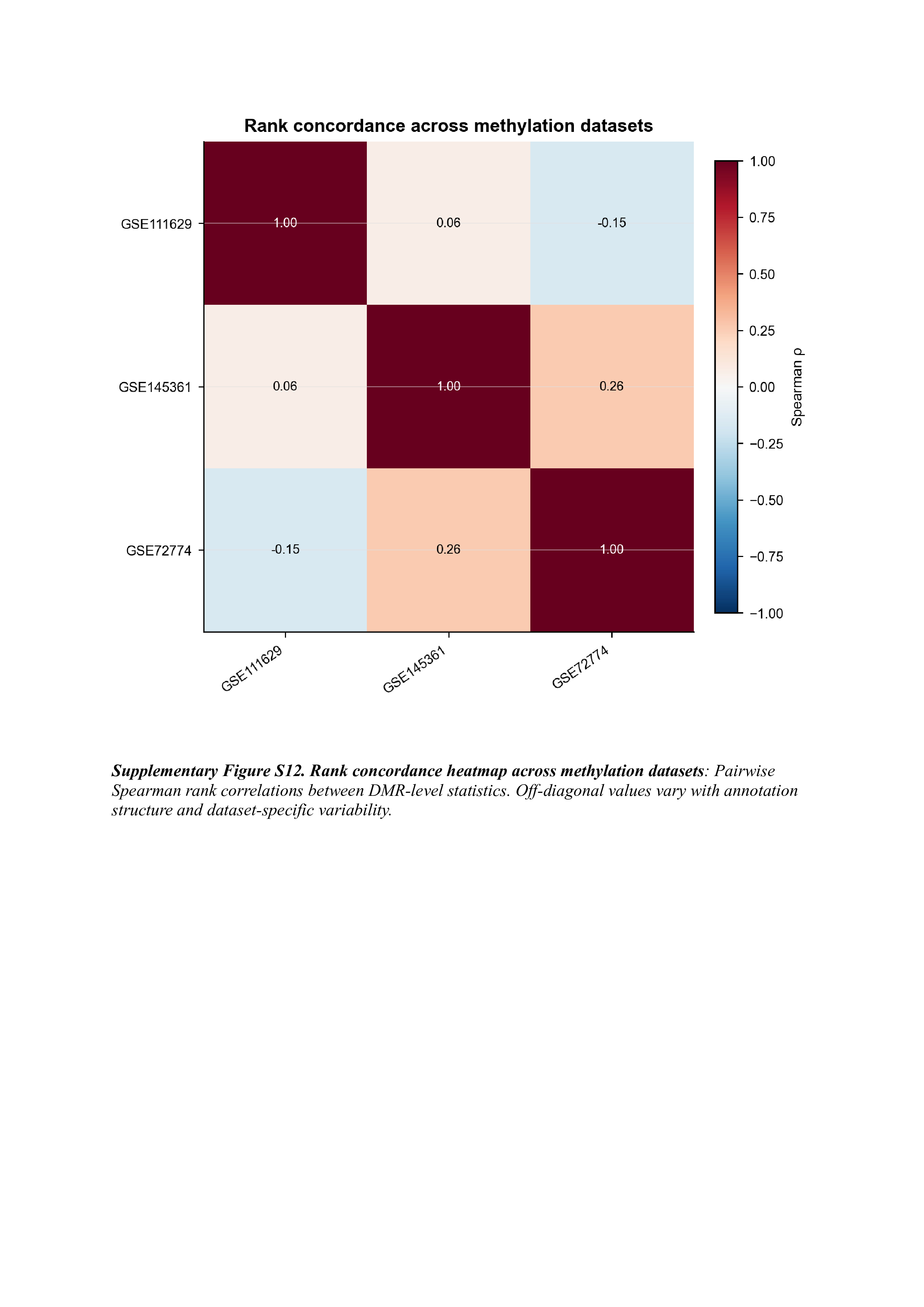
